# Thermotolerance in Chia (*Salvia hispanica* L.) is Mediated by Rapid Heat-Induced Transcriptomic Reprogramming and Lipid Remodelling in Leaves

**DOI:** 10.1101/2025.05.30.656991

**Authors:** Tannaz Zare, Cheka Kehelpannala, Atul Bhatnagar, Thusitha W. Rupasinghe, Berit Ebert, Alexandre Fournier-Level, Ute Roessner

## Abstract

Heat stress poses a significant threat to crop productivity; however, the thermotolerance mechanisms in underutilised oilseed crops, such as chia (*Salvia hispanica* L.), remain poorly understood. Despite the growing interest in chia as a rich source of ω-3 fatty acids, its molecular response to heat stress, particularly in vegetative tissues, has not been explored. We conducted transcriptomic and lipidomic profiling to examine how chia leaves respond to short-term (3 h) and prolonged (27 h) heat stress, followed by recovery under ambient conditions. Heat stress induced differential expression in over 20% of transcripts in chia leaves, with distinct patterns involving Ca²⁺ signalling, heat shock factors, and other biological pathways contributing to cellular homeostasis. Gene expression and lipid profiles in chia leaves responded dynamically to both short-term (3 h) and prolonged (27 h) heat stress (38°C/20°C). An almost complete return to baseline was observed, with all but 0.3% of heat-responsive genes reverting to control expression levels after 24 h. Our analysis confirms the role of Ca²⁺-mediated signalling pathways and molecular chaperones, including heat shock proteins and heat shock factors, which have been previously shown to contribute to maintaining cellular function during heat stress in other plant species. Among 287 annotated lipid species, TGs exhibited the most significant and reversible changes (>2-fold), suggesting their involvement in membrane remodelling. Our findings reveal adaptive mechanisms in chia that may open avenues for enhancing thermotolerance in other heat-sensitive oilseed crops.

**SIGNIFICANCE STATEMENT:** Understanding how crops respond to heat stress is crucial as global temperatures rise. This study indicates that in chia (*Salvia hispanica*), pathways responding to heat stress, like calcium signalling and heat shock proteins, are rapidly activated, and leaf triacylglycerol levels rise under heat stress before returning to baseline during recovery. By demonstrating near-complete recovery of gene expression following heat exposure, these findings highlight mechanisms of thermotolerance that may support improved stress tolerance in other crops.

## INTRODUCTION

Climate change poses a significant threat to agriculture, with global temperatures projected to rise by 1.0–5.7 °C by 2100 (Lee et al., 2024), thereby placing food production and security at risk (Bibi and Rahman, 2023). Among environmental stressors, heat stress impairs plant development by altering gene expression, protein synthesis, and metabolism (Jagadish et al., 2021). Prolonged high temperatures reduce photosynthesis and water potential, increase respiration and transpiration, and cause oxidative damage (Lippmann et al., 2019, Jagadish et al., 2021). Elevated heat also disrupts membrane stability, triggering signalling networks and adaptive responses to restore homeostasis (Chaudhry and Sidhu, 2022, Ding and Yang, 2022). Understanding these molecular responses is crucial for enhancing crop resilience to heat stress.

Heat stress impairs photosynthesis by damaging thylakoid membranes and disrupting electron transport (Zhao et al., 2020). It also triggers a rapid influx of cytosolic calcium ions (Ca²⁺), an early signalling event in the heat stress response (Saidi et al., 2010). This Ca²⁺ influx is mediated by several protein channels, including cyclic nucleotide-gated channels (CNGCs), glutamate receptor-like channels (GLRs), and mechanosensitive (MS) channels (Ghosh et al., 2022, Park and Shin, 2022, Kang et al., 2023, Oranab et al., 2023). In Arabidopsis (*Arabidopsis thaliana*) and rice (*Oryza sativa*), specific CNGC proteins, such as AtCNGC6, OsCNGC14, and OsCNGC16, promote thermotolerance by modulating Ca²⁺ signalling and activating heat-responsive genes (Gao et al., 2012, Cui et al., 2020).

Ca²⁺ ions act as key secondary messengers in the heat stress signalling pathway. Plants sense changes in cytosolic Ca²⁺ concentrations through calcium-binding sensor proteins, including calmodulin (CaM), calmodulin-like proteins (CMLs), Ca²⁺-dependent protein kinases (CDPKs), and calcineurin B-like proteins (CBLs), which interact with CBL-interacting protein kinases (CIPKs) to activate downstream stress-responsive signals (Pan et al., 2019, Jarratt-Barnham et al., 2021). In Arabidopsis, the loss of *AtCaM3* reduces thermotolerance, whereas its overexpression improves it (Zhang et al., 2009). Similar effects are seen with *CsCaM3* in cucumber (*Cucumis sativus*) (Yu et al., 2018) and *OsCaM1-1* in rice (Wu et al., 2012). CDPKs and CBL-CIPK complexes also contribute to heat tolerance and adaptation. Overexpressing *VaCPK29* in grapevine (*Vitis amurensis*) and *ZmCDPK7* in maize (*Zea mays*) improves heat tolerance by maintaining photosynthesis or enhancing signalling (Dubrovina et al., 2017, Zhao et al., 2021). In rice, *OsCBL8–OsCIPK17* enhances heat resilience by phosphorylating the transcription factor *OsNAC77* (Gao et al., 2022).

Heat shock rapidly elevates cytosolic Ca²⁺ levels and disrupts normal Ca²⁺ oscillations (Weigand et al., 2021). This influx is detected by membrane-localised proteins that activate heat shock protein (HSP) genes (Bourgine and Guihur, 2021, Haider et al., 2022). The HSPs, classified into five major families: HSP100, HSP90, HSP70, HSP60, and small HSPs, act as molecular chaperones to stabilise proteins and maintain cellular integrity under stress (Kang et al., 2023, Kumar et al., 2024). Their expression is regulated by heat stress transcription factors (HSFs), which are classified into three classes: HSFA, HSFB, and HSFC (Haider et al., 2022). Under normal conditions, HSFs are sequestered in cytosolic HSP complexes; heat stress triggers their release, phosphorylation, and nuclear translocation, leading to transcriptional activation of HSP genes (Swindell et al., 2007, Andrási et al., 2021).

HSFA1 proteins are master regulators of the heat shock response, activating downstream HSFs such as HSFA2, HSFA3, HSFA7, and HSFBs (Haider et al., 2022). Overexpression of HSFA1 enhances basal thermotolerance in tomato (*Lycopersicon esculentum*), while suppression of HSFA2 and HSFB1 increases heat sensitivity (Mishra et al., 2002). HSFA3 supports thermomemory in Arabidopsis by interacting with HSFA2 and recruiting H3K4 methylation (Friedrich et al., 2021). Overexpression of wheat (*Triticum aestivum*) TaHsfA5 enhances thermotolerance in transgenic Arabidopsis and rice plants during both stress and recovery phases (Samtani et al., 2023). Similarly, TaHSF6b overexpression in barley (*Hordeum vulgare*) activates the transcription of major heat shock proteins (HSP70-2, HSP70-5, HSP90-2, and HSP18.1), thereby improving reactive oxygen species (ROS) homeostasis (Poonia et al., 2020). In tomato, HSFA7 regulates HSFA1a to support long-term adaptation (Mesihovic et al., 2022). Although most HSFCs respond to oxidative or salinity stress, HSFC2a enhances grain thermotolerance in wheat through ABA-mediated signalling (Hu et al., 2018). The HSPs further support thermotolerance by stabilising membranes and limiting lipid peroxidation (Bourgine and Guihur, 2021).

Heat stress disrupts lipid bilayers, compromising membrane integrity, altering permeability, and reducing oil accumulation in seeds (Higashi and Saito, 2019). Moderate temperatures (30-45 °C) affect thylakoid membrane stacking and chloroplast morphology, while temperatures above 45 °C induce glycerolipid phase separation and protein denaturation (Higashi and Saito, 2019, Zeng et al., 2021). Heat also increases the ratio of unsaturated to saturated fatty acids (FAs) in membrane lipids, thereby enhancing fluidity and promoting lipid transfer, as well as ROS-induced lipid peroxidation (Higashi et al., 2015, Shiva et al., 2020, Bourgine and Guihur, 2021). Lipids form the structural foundation of cellular membranes and play central roles in plant stress responses (Higashi and Saito, 2019, Bourgine and Guihur, 2021). Lipid classes such as phosphatidylcholine (PC), phosphatidic acid (PA), lysophosphatidylcholine (LPC), inositol phosphate (IP), diacylglycerol (DG), triacylglycerol (TG), sphingolipids (SLs), N-acylethanolamines (NAEs), sterols, and oxylipins are involved in plant adaptation to heat (Higashi et al., 2015, Higashi and Saito, 2019, Shiva et al., 2020, Bourgine and Guihur, 2021).

TGs accumulation and lipid remodelling contribute to thermotolerance by stabilising membranes under heat stress (Mueller et al., 2015, Lu et al., 2020, Zhang et al., 2022). For example, TG content increases under heat stress in Arabidopsis (Mueller et al., 2017, Shiva et al., 2020), wheat (Djanaguiraman et al., 2018), peanut (*Arachis hypogaea*) (Zoong Lwe et al., 2020), and castor bean (*Ricinus communis*) (Zhang et al., 2022). In Arabidopsis, *pdat1* mutants lacking phospholipid:diacylglycerol acyltransferase 1 (PDAT1) exhibited impaired TGs accumulation and heat sensitivity (Mueller et al., 2017). Heat stress promotes the conversion of DGs and PC-derived FAs into TGs and LPCs, redirecting lipids from galactolipids into extraplastidic pools (Mueller et al., 2017, Shiva et al., 2020). In wheat and castor bean, stress also alters thylakoid lipids and enhances DGAT-dependent TG biosynthesis (Djanaguiraman et al., 2018, Zhang et al., 2022). While progress has been made in understanding how plants adapt lipid metabolism under heat stress, further research is needed to clarify how such mechanisms support homeostasis in oilseed crops.

*Salvia hispanica (chia)*, a Lamiaceae species increasingly recognised for its high ω-3 FAs content and potential as a nutraceutical, has sparked growing interest in its application as a health-promoting oilseed crop (Zare et al., 2024a). Chia is commercially cultivated in Australia’s Kimberley region, where projected climate scenarios indicate that the number of days exceeding 35 °C will rise dramatically in the coming decades (Crawford et al., 2012, Quicke, 2019). Although several studies have advanced chia’s genomic and molecular resources (Sreedhar et al., 2015, Peláez et al., 2019, Wimberley et al., 2020, Gupta et al., 2021, Wang et al., 2022, Alejo-Jacuinde et al., 2023, Zare et al., 2024b), they have not addressed heat stress responses in leaves. A recent proteomic analysis examined seed viability loss in chia under heat and humidity stress (Rodríguez et al., 2024); however, no transcriptomic or lipidomic studies have investigated vegetative tissues exposed to acute or sustained heat stress.

To address this gap, we examined how chia leaves respond to heat shock, prolonged heat stress, and subsequent recovery using transcriptomic and lipidomic analyses. Leveraging a recently published reference genome (Zare et al., 2024b), we employed RNA sequencing to assess changes in gene expression and liquid chromatography-tandem mass spectrometry (LC–MS) to profile lipid composition. This integrative study aims to improve our understanding of the impact of heat stress on gene regulation and lipid metabolism in chia, a nutritious yet underexplored oilseed crop.

## RESULTS

### Transcriptomic profiling shows distinct temporal responses to heat stress in chia

The Multidimensional scaling (MDS) and principal component (PC) analyses of transcriptomic data revealed a strong transcriptional shift in response to heat stress. Samples from heat-treated plants (T1.H and T2.H) were distinctly separated from both control and recovery samples along the first dimension (Dim1) (Figures 1a, 1b), indicating substantial transcriptomic reprogramming during stress and a return to baseline during recovery. The separation between T1.H (3 hr heat shock) and T2.H (27 hr prolonged heat stress) implies an enhanced and evolving response to prolonged heat exposure. Hierarchical clustering (Figure 1d) further supported these patterns, indicating that the variation between treatment groups exceeded the variation within groups. Notably, the T3.R (recovery) group clustered closely with the control samples (T0, T1C, T2C, T3C), suggesting that gene expression levels returned to initial control levels after 24 hr of recovery. Additionally, the second dimension (Dim2) of the MDS plot appeared to capture variation related to plant growth, although some differences were also noted among the control samples (Figure 1b).

**Figure 1.**
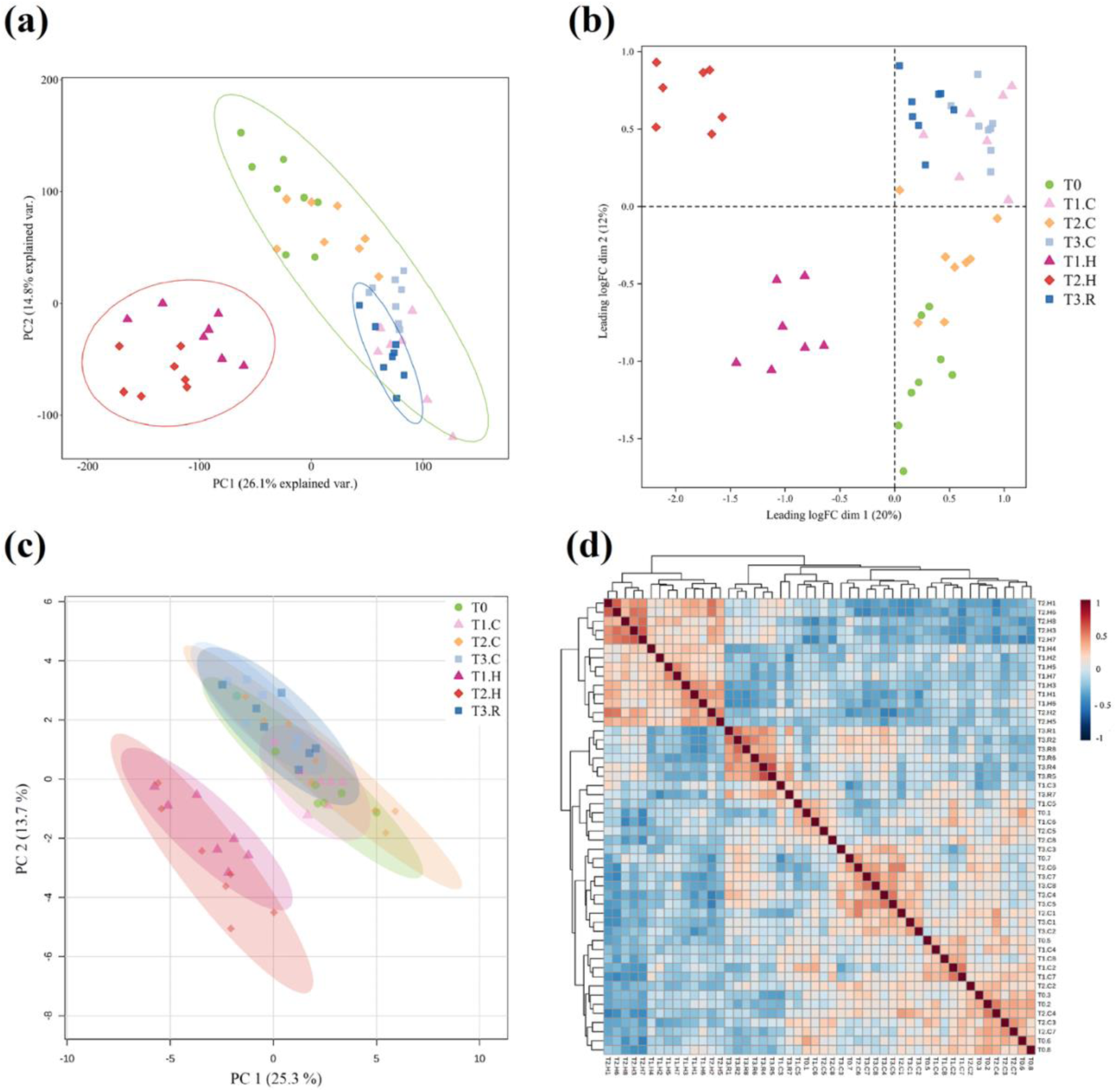
Transcriptomic and lipidomic responses of chia leaves to heat stress and recovery. Analysis was performed with n = 8 biological replicates per group per time point. (a) Principal component analysis (PCA) of RNA-seq data (ellipses indicate 95% normal probability regions for control, heat shock, and recovery groups); (b) Multidimensional scaling (MDS) plot of RNA-seq data based on leading log-fold changes; (c) PCA of lipidomic profiles (ellipses show 95% normal probability regions); and (d) Heatmap and hierarchical clustering of Pearson correlation coefficients for lipid profiles across all samples.

### Short-term heat stress induces lipidomic remodelling and partial recovery in chia

The strong separation between the heat-treated (T1.H and T2.H) and control groups, observed at the transcriptomic level, was also evident in the lipid composition of chia leaves (Figure 1c). Lipid levels in the recovery group (T3.R) returned to control levels, indicating near-complete recovery consistent with the transcriptomic response. Lipid levels at T1.H and T2.H were also distinct from those in other groups (Figure 1d), confirming a strong lipidomic response to heat stress. Although T3.R samples clustered with the control groups, they formed a separate branch, suggesting that recovery at the lipid level may be slower than at the transcriptomic level. The similarity in lipid and transcript profiles points to a coordinated response of gene expression and lipid metabolism in chia leaves under short-term heat stress.

### Short-term and prolonged heat stress elicit distinct transcriptomic responses in chia

To identify differentially expressed genes (DEGs) under heat stress, the transcriptomes of the T1.H and T2.H groups were compared with the control group at T0 (Figure 2). The large number of DEGs in both comparisons (23% and 20% of the 27,151 filtered genes, respectively; QL F-test, FDR = 0.1%, *P* < 0.001) indicates a whole-transcriptome response to heat stress, affecting multiple biological processes. In response to short-term heat shock (T1.H), 2998 genes were up-regulated (including 934 with log_2_FC > 2), while 3148 were down-regulated (751 with log_2_FC < –1) (Figures 2a & c). After 24 additional hours of heat exposure (T2.H), 2476 genes were up-regulated (964 with log_2_FC > 2) and 2909 were down-regulated (699 with log_2_FC < – 1) compared with the T0 samples (Figures 2b & d). While the short-term heat stress response involved a balanced pattern of gene regulation, prolonged exposure resulted in predominantly down-regulated transcripts. This suggests that distinct sets of genes may be involved in the early versus prolonged phases of chia’s response to heat stress.

**Figure 2.**
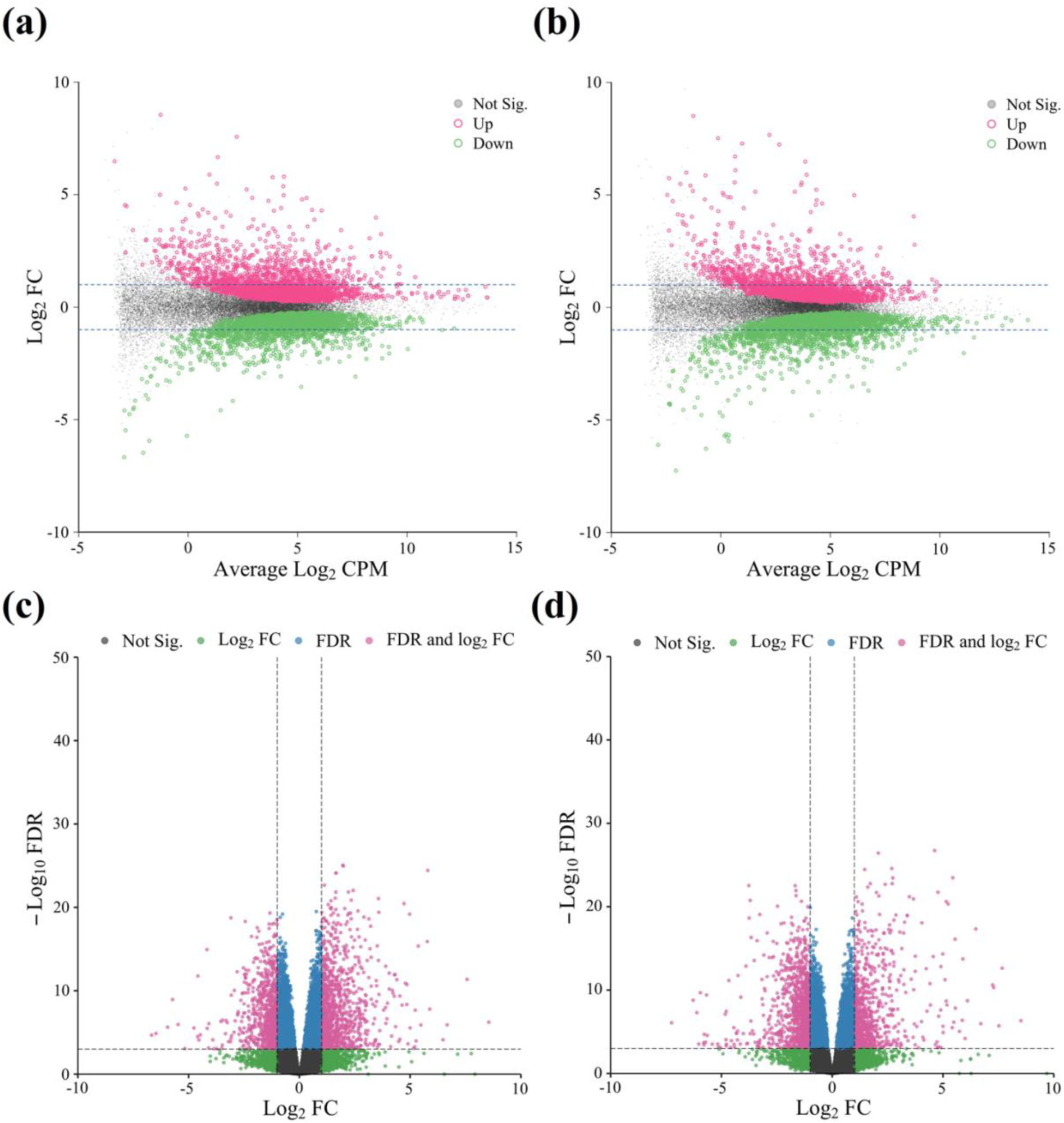
Differential gene expression in chia leaves under short-term and prolonged heat stress. Mean–difference plots for (a) T1.H (3 hr heat shock) and (b) T2.H (27 hr prolonged heat stress) show log₂ fold change (log₂FC) against average gene abundance (log₂CPM). Significantly up-regulated and down-regulated genes (FDR = 0.1%, *P* < 0.001) are highlighted in pink and green, respectively. Volcano plots for (c) T1.H (3 hr heat shock) and (d) T2.H (27 hr prolonged heat stress) display log₂FC on the x-axis and –log₁₀(FDR) on the y-axis. Genes are highlighted based on FDR < 0.1% and absolute log₂FC ≥ 1.

### Transcriptomic profile of chia leaves returns to baseline after 24 hr recovery from heat stress

To determine whether chia could maintain transcriptional homeostasis following heat stress, gene expression in recovery samples (T3.R) was compared with that in control samples (T0). Of the 27,151 genes analysed, only 13 showed differential expression (QL F-test, FDR = 0.1%, *P* < 0.001). This represents less than 0.3% of the differentially expressed genes identified in the heat-stress versus control comparisons. The extremely low number of DEGs following recovery indicates that transcriptomic profiles largely returned to baseline within 24 hours.

### Identification of heat-responsive genes in chia leaves during heat stress and recovery phases

Exposure to heat stress in chia leaves triggered differential expression of multiple gene families, including *HSFs*, *HSPs*, Ca²⁺-signalling genes, and molecular chaperones (QL F-test, FDR = 0.1%, *P* < 0.001; Supplemental Figure S6). The *HSFs* belonged to class A (A2, A4, A5, A7), class B (B2), and class C (C1) gene families. All were up-regulated during heat stress, with stronger responses during heat shock (T1.H) than during prolonged heat (T2.H) and returned to control levels upon recovery (Supplemental Figure S6a).

Heat shock proteins (*HSPs*) were more strongly induced by prolonged heat exposure (T2.H), with most genes being up-regulated (Supplemental Figure S6b). These included high and medium molecular weight members such as HSP70, HSP80, HSP83, and HSP90, and a broad range of small *HSPs* (*sHSPs*) including *HSP1*, *HSP15.7*, *HSP17*, *HSP17.3*, *HSP17.4*, *HSP17.8*, *HSP18.2*, *HSP21.7*, *HSP21.9*, *HSP22* and *HSP26.5*. Notably, the majority of up-regulated *HSP* genes belonged to the *HSP70* family.

Genes associated with Ca²⁺ influx and signalling pathways showed a stronger transcriptional response to heat shock than to prolonged heat exposure (Supplemental Figure S6d). This highlights the role of Ca²⁺ as an early signal transducer in how plants respond to heat stress, further emphasising its importance in plant responses to abiotic stress (Li et al., 2022, Kan et al., 2023). Differential expression was observed across a wide range of calcium-related genes encoding calcium-binding proteins (*CBP*), calcium-dependent lipid-binding proteins (*CLB*), and various calcium-dependent protein kinases (*CDPK1*, *CDPK4*, *CDPK-SK5*, *CDPK10*, *CDPK11*, *CDPK13*, *CDPK28*, and *CDPK32*). Other up-regulated genes included calcium-transporting ATPase genes (*ATPase1*, *ATPase2*, and *ATPase13*), genes encoding the calcium-binding mitochondrial carrier protein (*SCaMC*), calcium-permeable stress-gated cation channel 1 (*CSC1*), calcium-sensing receptor (*CaSR*), mitochondrial calcium uniporter 2 (*MCU2*), calcium uptake protein (*CUP*), calmodulin calcium-dependent NAD kinase (*CaMK*), cation/calcium exchangers (*CCX1*, *CCX4*), and calmodulin-like proteins (*CML1*, *CML13*, *CML36*, *CML48*). Additional responsive genes included a sodium/calcium exchanger (*NCL*), a calcium-binding allergen gene, and two-pore calcium channel protein 1-A (*TPC1A*).

Chaperone proteins included members of the Bcl-2-associated athanogene (BAG) gene family (*BAG1*–*BAG6*), caseinolytic protease ClpB chaperones (*ClpB1*, *ClpB3*), and multiple *DnaJ*/Hsp40 genes (including *DnaJ1*, *DnaJ6*, *DnaJ8*, *DnaJ10*, *DnaJ11*, *DnaJ15*, *DnaJ16*, *DnaJ20*, *DnaJ49*, *DnaJ50*, and *DnaJ76*). Additional heat-responsive chaperones included copper chaperone for superoxide dismutases (*CCS*), the iron–sulfur cluster co-chaperone protein *HSCB*, the histone chaperone *ASF1B*, and *proteasome assembly chaperone 2*. These transcripts showed a mixture of up- and down-regulation and were primarily detected after the immediate heat shock (Supplemental Figure S6c). Molecular chaperones belonging to the FK506-binding protein (FKBP) family were mostly up-regulated and showed strong responses to both heat treatments (Supplemental Figure S6e). These included *FKBP12*, *FKBP16*–*FKBP20*, *FKBP43*, and *FKBP62*. Additional responsive genes included the peptidyl-prolyl cis– trans isomerases and NIMA-interacting protein 4 (*Pin1* and *Pin4*, respectively), along with multiple cyclophilin genes, including *CYP21*, *CYP26*, *CYP28*, *CYP37*, *CYP38*, *CYP40*, *CYP59*, and *CYP90*.

In addition to the gene families summarised in Figure S6, one multiprotein bridging factor 1b (*MBF1b*) gene showed increased expression during both heat shock (T1.H) and prolonged heat stress (T2.H). At T1.H, two dehydration-responsive element-binding factor (*DREB*) genes were down-regulated, whereas two others *DREB1b* and *DREB2c* were up-regulated at T2.H. These transcription factors play a crucial role in abiotic stress, and their varied expression patterns underscore the complex regulation of gene expression under heat stress (Jaimes-Miranda and Chávez Montes, 2020, Singh and Chandra, 2021).

A set of 100 lipid-related genes, selected from existing literature (Rawsthorne, 2002, Li-Beisson et al., 2013, He et al., 2020), was used to explore the transcriptional responses of lipid metabolism pathways to heat stress. These genes represent key steps in plastidial FA synthesis, FA elongation/desaturation, chloroplast export, plastidial and microsomal glycerolipid synthesis, galactolipid and sulfolipid synthesis, eukaryotic phospholipid synthesis, TG synthesis, lipid degradation, and β-oxidation (Supplemental Table S3). Of these, 95 genes were differentially expressed under heat stress (T1.H and/or T2.H), while none were differentially expressed during recovery compared to control (T3.R) (Supplemental Table S4).

### Clustering of DEGs shows distinct expression patterns during heat stress and recovery in chia

DEGs with log₂FC > 1, including 1685 genes in T1.H (3 hr heat shock) and 1663 genes in T2.H (27 hr prolonged heat stress), were grouped into four clusters (A to D) based on their temporal expression profiles (Figure 3). Under heat shock (T1.H) (Figure 3a and 3b), cluster A genes were significantly down-regulated after 3 hours but gradually returned to near control levels after prolonged heat exposure (T2.H) and showed a slight up-regulation during the recovery phase (T3.R) (Figure 3b). In contrast, the DEGs in cluster D exhibited a reverse pattern to those in cluster A; their expression increased sharply at 3 hr (T1.H), returned to control levels after 27 hr of continued heat exposure (T2.H), and was slightly down-regulated during recovery (T3.R). These contrasting temporal patterns suggest that clusters A and D may be involved in different stages of the transcriptional response to acute heat stress (Figure 3b).

**Figure 3.**
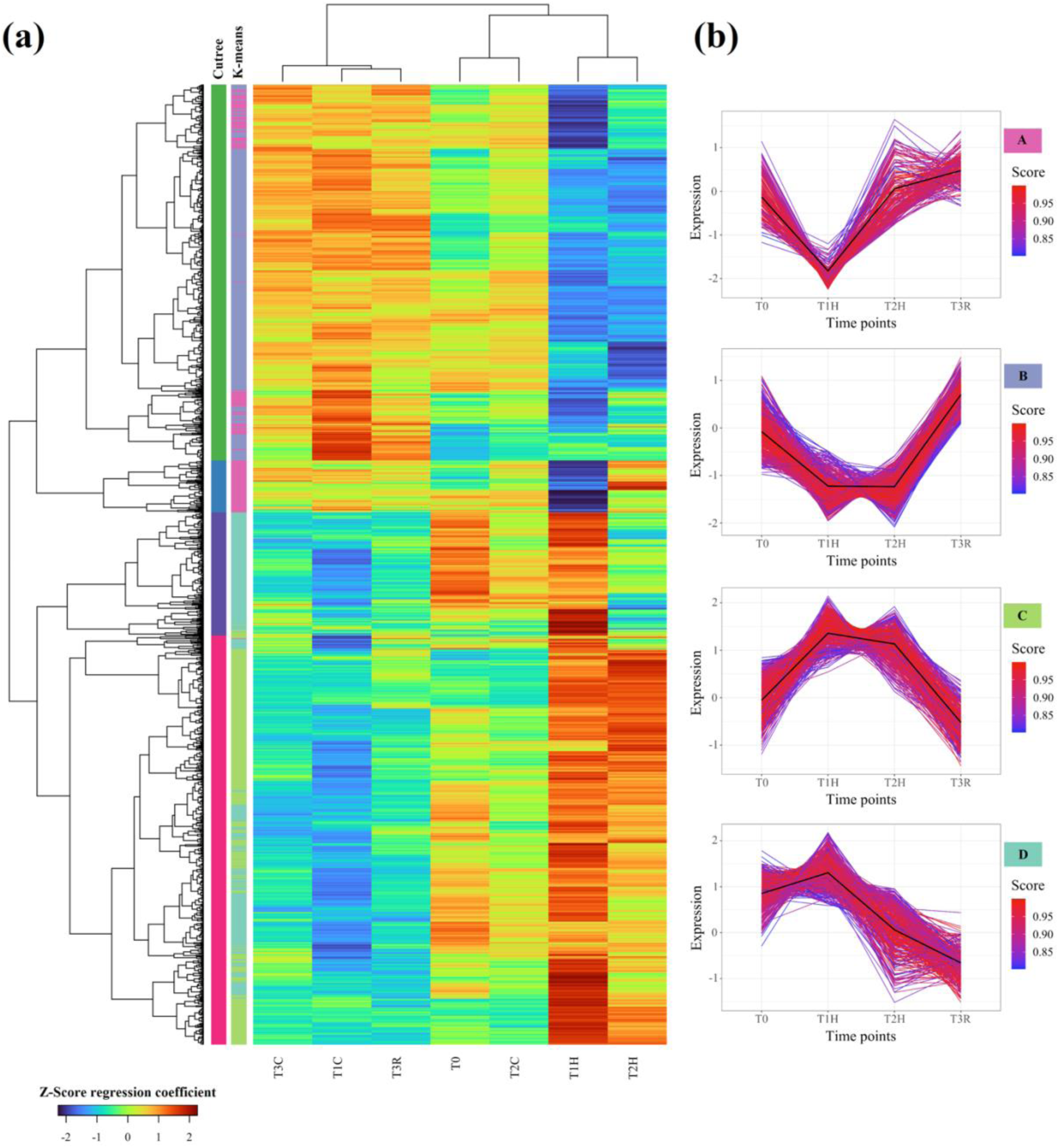
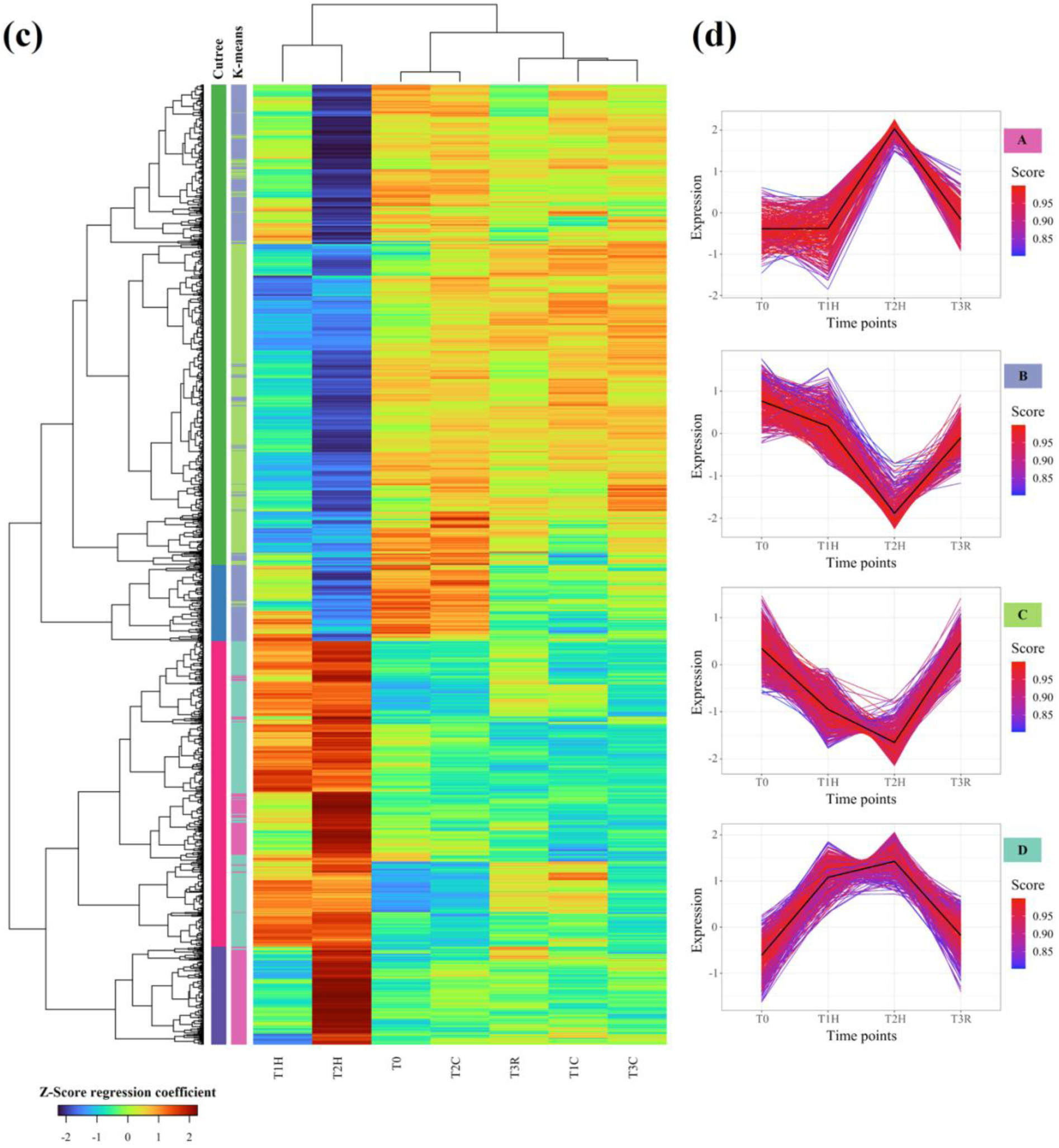
Clustering of differentially expressed genes (DEGs) during heat stress in chia. Heat-responsive DEGs were clustered based on expression profiles under two conditions: (a, b) heat shock (T1.H vs T0) and (c, d) prolonged heat stress (T2.H vs T0). (a, c) Heatmaps display z-scores of normalised regression coefficients from the NB-QL model. Each row represents a DEG, and each column corresponds to a specific treatment or time point. Hierarchical clustering was performed using 1-correlation (Z-score) with the Canberra distance metric. The left vertical colour bar indicates clusters defined by hierarchical tree cutting (“cutree”), while the right colour bar shows groups derived from K-means clustering (k = 4). (b, d) Line plots illustrate temporal expression trends for the four K-means clusters (A–D). Only DEGs with a membership score ≥ 0.8, indicating strong association with a cluster centroid, were included. Coloured boxes label the cluster identities (A–D), and black lines represent the average expression trend within each cluster.

Clusters B and C exhibited different transcriptional responses, showing initial up- or down-regulation after 3 hr of heat exposure (T1.H). Expression levels then remained relatively stable during prolonged exposure (T2.H), before returning to control levels after 24 hr of recovery (T3.R) (Figure 3b). This pattern suggests that genes in clusters B and C are involved in maintaining heat stress responses over time, possibly contributing to thermotolerance or sustained adaptation. The expression analysis of DEGs at T2.H (Figure 3c and 3d) revealed different temporal expression profiles from those observed at T1.H (Figure 3a).

Genes in clusters A and B (Figure 3d) showed little to no change after 3 hr of heat exposure but showed a marked shift in expression after 27 hr, suggesting a delayed transcriptional response. Notably, their expression returned to control levels after 24 hr of recovery (T3.R), indicating that the response was reversible. This contrasts with clusters C and D (Figure 3d), which showed a rapid transcriptional response that was evident after 3 hr and continued to rise, albeit more gradually, after 27 hr. After recovery, clusters A and B returned to control levels, while clusters C and D remained overexpressed relative to baseline (T0) (Figure 3d).

### Gene Ontology enrichment reveals distinct transcriptional and cellular responses during heat stress and recovery

At T1.H (3 hr heat shock), DEGs were significantly enriched in 24 biological processes, 27 molecular functions, and 14 cellular compartments (FDR ≤ 5%, *P* < 0.05; Supplemental Tables S5–S7). In contrast, at T2.H (prolonged heat stress), there were fewer enriched terms, including 18 biological processes, 19 molecular functions, and 10 cellular compartments (Supplemental Tables S8–S10), indicating a shift in the transcriptional response. Although only 13 genes were differentially expressed at T3.R (recovery), they were associated with 20 biological processes, 19 molecular functions, and 11 cellular compartments, suggesting transcriptional fine-tuning and restoration of cellular balance. DEGs at T1.H were also enriched in the Cul4A-RING E3 ubiquitin ligase complex, calcium and cation channel complexes, and endoplasmic reticulum (ER)-Golgi intermediate compartments (Supplemental Table S7), suggesting activation of ubiquitin-mediated protein regulation, calcium-mediated signalling, and vesicle trafficking in response to heat stress (Li et al., 2022, Sato et al., 2024, Wang and Xie, 2024).

At T2.H, enriched biological processes reflected detoxification and transcriptional adaptation, including formaldehyde metabolism (GO:0046292), formaldehyde catabolic process (GO:0046294), aldehyde detoxification (GO:0110095), cellular response to aldehyde (GO:0110096) and transcription elongation (GO:0034243) (Supplemental Table S8). Enriched molecular functions under prolonged stress included clathrin heavy chain binding (GO:0032050) and phosphatidylinositol binding (GO:0005545), pectate lyase activity (GO:0030570), glucan endo-1,3-beta-D-glucosidase activity (GO:0042973), and polygalacturonase activity (GO:0004650), indicating vesicle trafficking and membrane signalling were also involved (Supplemental Table S9). Enriched cellular compartments at T2.H included the Cul4A-RING E3 ubiquitin ligase complex (GO:0031464), clathrin-coated pit (GO:0005905), nucleosome (GO:0000786), DNA packaging complex (GO:0044815), and anchored component of plasma membrane (GO:0046658), suggesting structural and chromatin-level responses to prolonged heat stress (Supplemental Table S10).

These findings suggest that early heat stress responses in chia primarily involve calcium signalling and structural remodelling, while prolonged exposure activates detoxification and chromatin-associated pathways. The limited but targeted enrichment seen after recovery indicates a rapid return to transcriptional homeostasis.

### Triacylglycerols exhibit the strongest lipidomic response to heat stress in chia leaves

To assess the effects of heat stress on lipid composition, we profiled the lipidome of chia leaves using mass spectrometry. A total of 325 lipid species were detected across all samples, of which 287 were annotated (Supplemental Table S11). We focused on 14 major lipid classes: (i) glycerolipids — TG, DG, monogalactosyldiacylglycerol (MGDG), digalactosyldiacylglycerol (DGDG), and sulfoquinovosyldiacylglycerol (SQDG); (ii) glycerophospholipids — PC, phosphatidylglycerol (PG), phosphatidylinositol (PI), phosphatidylethanolamine (PE), phosphatidylserine (PS), and LPC; (iii) sphingolipids — ceramide (Cer) and hexosylceramide (HexCer); and (iv) cardiolipin (CL).

A pairwise comparison of average lipid peak areas revealed clear separation between heat-treated and control samples along PC1 for both T1.H (3 hr heat shock) and T2.H (27 hr prolonged heat stress) (Figure 4a & c). Additionally, PC2 separated T3.R (recovery) samples from their respective controls (T3.C) (Figure 4e), highlighting temporal changes in lipid composition across treatments. Heat shock primarily led to a significant increase in lipid species abundance (*t*-test, FDR = 5%, *p*-adjusted < 0.05, FC > 1.2), with only one species, TG 50:2|TG 16:0/16:0/18:2, showing a significant reduction (Figure 4b; Supplemental Figures S7a and S8). A similar trend was observed under prolonged heat stress, with only eight lipid species showing a significant decrease in abundance, including TG 50:3|TG 16:0/16:0/18:3, PI 36:4, PG 36:4|PG 18:2/18:2, DG 36:4|DG 18:2/18:2, DG 36:2|DG 18:0/18:2, DG 34:2|DG 16:0/18:2, DG 32:1|DG 16:0/16:1, and Cer 44:1;4O|Cer 18:1;3O/26:0;(2OH)(a) (Figure 4d; Supplemental Figures S7b and S9). Conversely, lipid levels largely remained unchanged during the recovery phase (Figure 4f), indicating a reduction in heat-induced lipid biosynthesis.

**Figure 4.**
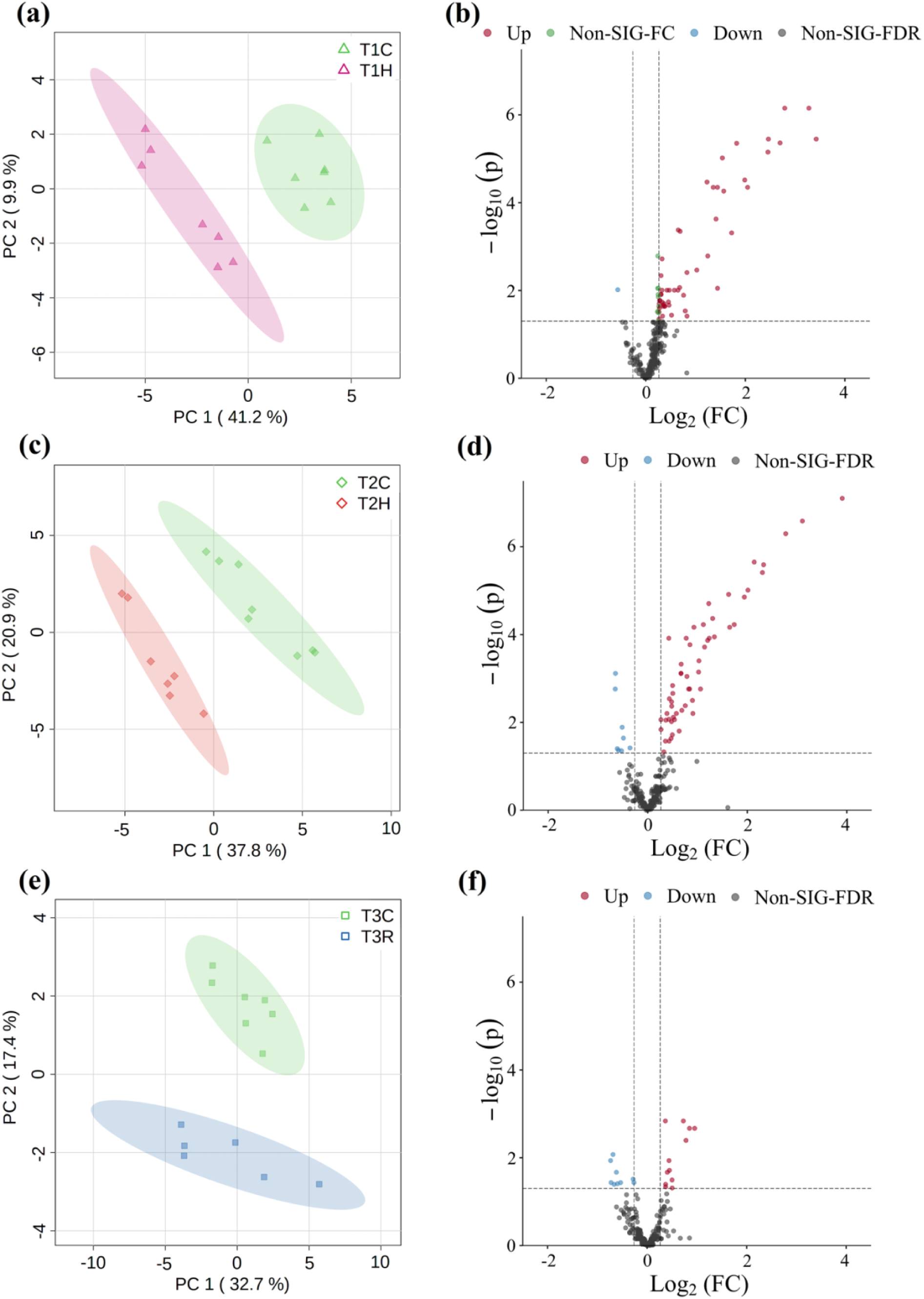
Pairwise comparisons of lipid profiles between treatment and control groups at each time point. (a, c, e) Principal component analysis (PCA) plots showing separation between control and treatment samples at T1 (heat shock), T2 (prolonged heat stress), and T3 (recovery), respectively. Each ellipse encompasses 95% of the estimated distribution for each group. (b, d, f) Volcano plots depicting differential abundance of lipid species between treatment and control groups at each time point. The x-axis shows the log₂ fold change (log₂FC), and the y-axis shows the −log₁₀ adjusted p-value (FDR). Lipid species with FDR < 0.05 and fold change ≥ 1.2 are highlighted: red (upregulated), blue (downregulated), and grey (non-significant).

Among the 14 lipid classes, 11, including TG, MGDG, DGDG, SQDG, PI, PE, PS, LPC, Cer, HexCer, and CL, showed significant and detectable changes in abundance in response to heat stress (student t-test, p-adjusted < 0.05; Figure 5). Most lipid classes showed subtle yet consistent changes in abundance in response to heat stress, with no significant difference between treatment and control at the recovery point (Figure 5b-k). However, TGs showed a pronounced change (>2-fold) in abundance in response to heat stress (Figure 5a), with TG 54:9|TG 18:3/18:3/18:3 and TG 52:9|TG 16:3/18:3/18:3 being identified as the most abundant species (Supplemental Figures S7a & b). TG and DGDG showed a significant increase at both heat stress time points, while MGDG and LPC showed a significant increase during prolonged heat stress (Figure 5a-c & j; Supplemental Figures S7a & b). Additionally, SQDG, HexCer, Cer, PI, PE, PS, and CL mainly increased significantly under heat shock stress (Figure 5d-i & k; Supplemental Figure S7a). Overall, these findings suggest that the lipidomic response to heat stress in chia was marked by both rapid and sustained shifts across key lipid classes. The TG lipid class exhibited the most dynamic and reversible changes, whereas membrane-associated lipids remained elevated under prolonged stress. These trends highlight the significance of lipid remodelling in chia’s response to heat stress and recovery.

**Figure 5.**
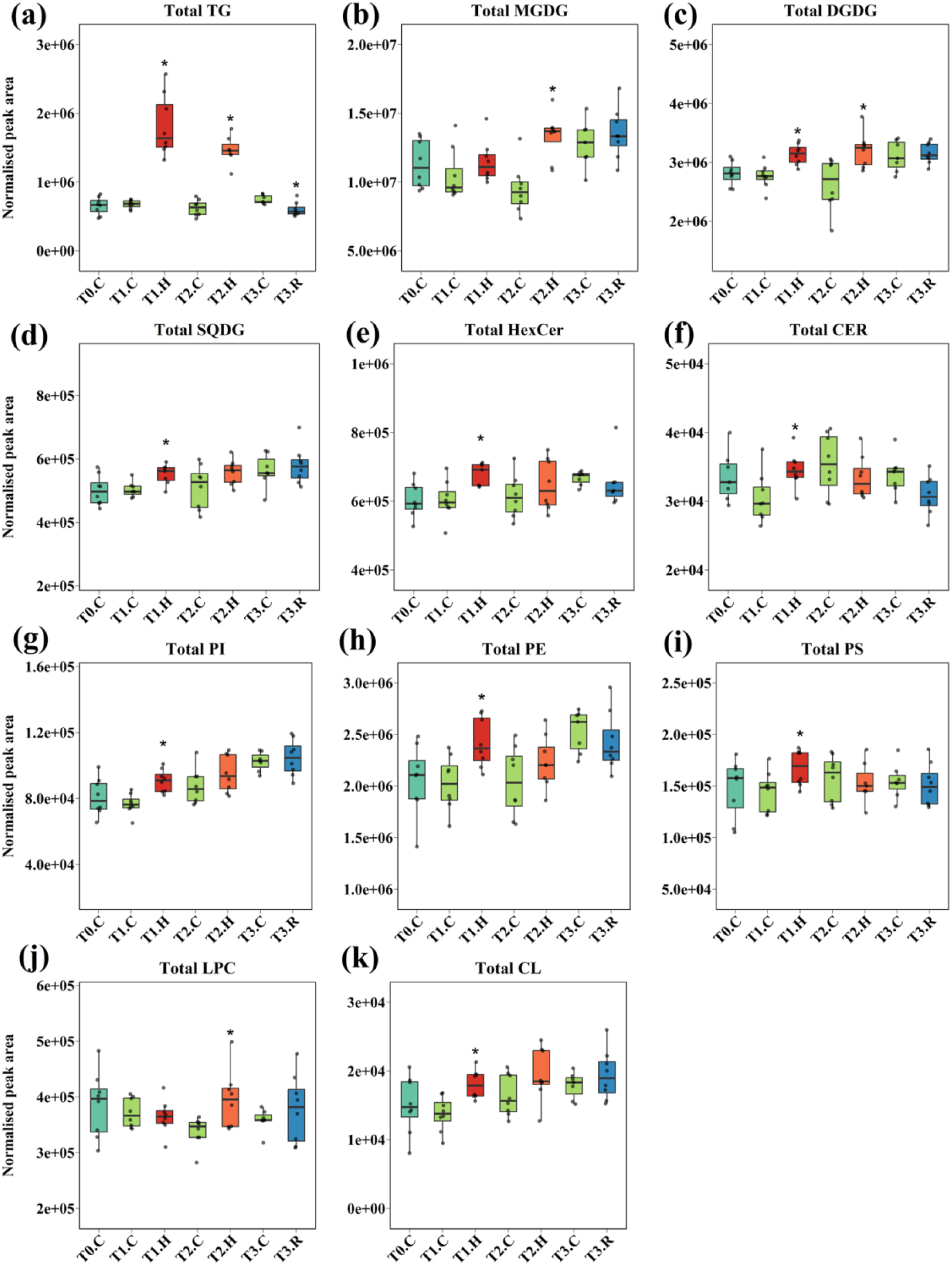
Abundance of lipid classes in chia leaves under heat stress and recovery. Boxplots illustrate the distribution of lipid peak intensities, normalised to 30 mg of fresh weight (n = 8). Horizontal lines represent medians, boxes cover the interquartile range (IQR), and whiskers extend to the most extreme values within 1.5 × IQR from the first and third quartiles. Points beyond this range are shown as outliers. Individual points signify biological replicates. Asterisks indicate statistically significant differences (FDR-adjusted p < 0.05) between paired control and treatment groups, evaluated using Student’s *t*-test. Cyan boxes indicate control samples (T0); Red boxes, 3-hour heat shock (T1.H); Orange boxes, 27-hour prolonged heat stress (T2.H); Blue boxes, 24-hour recovery (T3.R); and Green boxes, time-matched controls for each treatment (T1.C, T2.C, and T3.C). Outliers identified using the 1.5 × IQR rule that were excluded from statistical analyses and boxplots as follows: T3.C6 from all lipid classes, T1.H5 & T3.R3 from HexCer, T2.H5 from PS, T2.H4 from TG and T1.C1 & T1.H2 from SQDG. Abbreviations: TG, Triacylglycerol; MGDG, Monogalactosyldiacylglycerol; DGDG, Digalactosyldiacylglycerol; SQDG, Sulfoquinovosyldiacylglycerol; PI, Phosphatidylinositol; PS, Phosphatidylserine; PE, Phosphatidylethanolamine; LPC, Lysophosphatidylcholine; HexCer, Hexosylceramide; CL, Cardiolipin.

## DISCUSSION

Heat stress, driven by rising global temperatures, is a major environmental stressor that threatens plant productivity by disrupting cellular homeostasis and impairing physiological and molecular functions (Haider et al., 2021, Arif et al., 2025). In this study, we investigated how chia leaves respond to short-term (3 h) and prolonged (27 h) heat stress, as well as recovery following 24 hours under ambient conditions, using integrated transcriptomic and lipidomic analyses. Our findings reveal coordinated changes in gene expression and lipid composition across stress phases, with (i) broadly similar response patterns in both omics layers; (ii) distinct molecular responses induced by short versus prolonged heat exposure; and (iii) a notable recovery capacity, as transcriptomic profiles largely returned to baseline within 24 hours. These results highlight chia’s dynamic stress response and suggest key mechanisms underlying its thermotolerance.

### Heat shock and prolonged heat stress responses involve distinct biological processes

In chia leaves, heat shock and prolonged heat stress induce extensive transcriptomic and lipidomic responses, albeit with notable differences in timing and intensity. Heat shock led to a stronger immediate transcriptional response, whereas prolonged exposure resulted in a greater number of DEGs. Similar temporal transcriptional dynamics under heat stress have been reported in Chinese rose (*Rosa chinensis*) (Li et al., 2019), Arabidopsis (Wang et al., 2020), moss (*Physcomitrium patens*) (Elzanati et al., 2020), and Easter lily (*Lilium longiflorum*) (Zhou et al., 2022). The overrepresented GO terms and the four distinct temporal expression patterns indicate that short-term and prolonged heat stress engage different biological pathways, emphasising the existence of time-specific transcriptional responses in chia.

The distinct responses to heat shock and prolonged heat stress in chia were linked to different expression patterns, suggesting the involvement of separate regulatory networks, although potentially connected through shared biological processes (Abbas et al., 2022, Mondal et al., 2022). For example, in grapevine (*Vitis vinifera*) (Liu et al., 2012), field mustard (*Brassica rapa*) (Yue et al., 2021), and wheat (*Triticum aestivum*) (Mishra et al., 2021), gene expression networks responding to different heat treatments are associated with distinct biological processes. Identifying expression modules in chia supports the hypothesis that short-term and prolonged heat stress engage distinct transcriptional responses. Notably, some components of the short-term response may act as quick signals, responding more to the rate of temperature change rather than the absolute temperature. This difference could indicate varied sensing and signalling mechanisms that are activated during different phases of heat stress, highlighting the complex nature of temporal regulation in chia’s heat response.

### Mechanisms promoting homeostasis under heat stress in chia

#### Ca^2+^ signalling as an early response to heat stress

Genes involved in Ca²⁺ transport and signalling were strongly enriched among the early heat shock response in chia. Ca²⁺ plays a well-established role in abiotic stress responses, including heat stress, by activating calcium-binding proteins that initiate downstream signalling cascades (Edel and Kudla, 2016, Li et al., 2022). Ca^2+^ signalling is known to occur within minutes of the onset of heat stress (Saidi et al., 2010). Consistent with this, we observed that Ca²⁺ metabolism-related genes were repressed during the initial heat shock but returned to control levels during prolonged heat exposure (Figure 6). This pattern likely reflects a heat-induced increase in plasma membrane fluidity, allowing Ca²⁺ influx through ion channels (Liu et al., 2003). A similar transient rise in cytosolic Ca²⁺ has also been reported in moss (*P. patens*) (Saidi et al., 2009, Finka et al., 2012). The induction of Ca²⁺ signalling genes in response to heat stress has been documented in various plant species, including turfgrass (*Agrostis stolonifera*) (Xu et al., 2006), wheat (Qin et al., 2008), tobacco (*Nicotiana tabacum*) (Suri and Dhindsa, 2008), spinach (*Spinacia oleracea*) (Yan et al., 2016), and Arabidopsis (Rahmati Ishka et al., 2018). To mitigate Ca²⁺ toxicity and prevent cell death, HSFs are activated to initiate protective responses (Abdelrahman et al., 2020).

**Figure 6.**
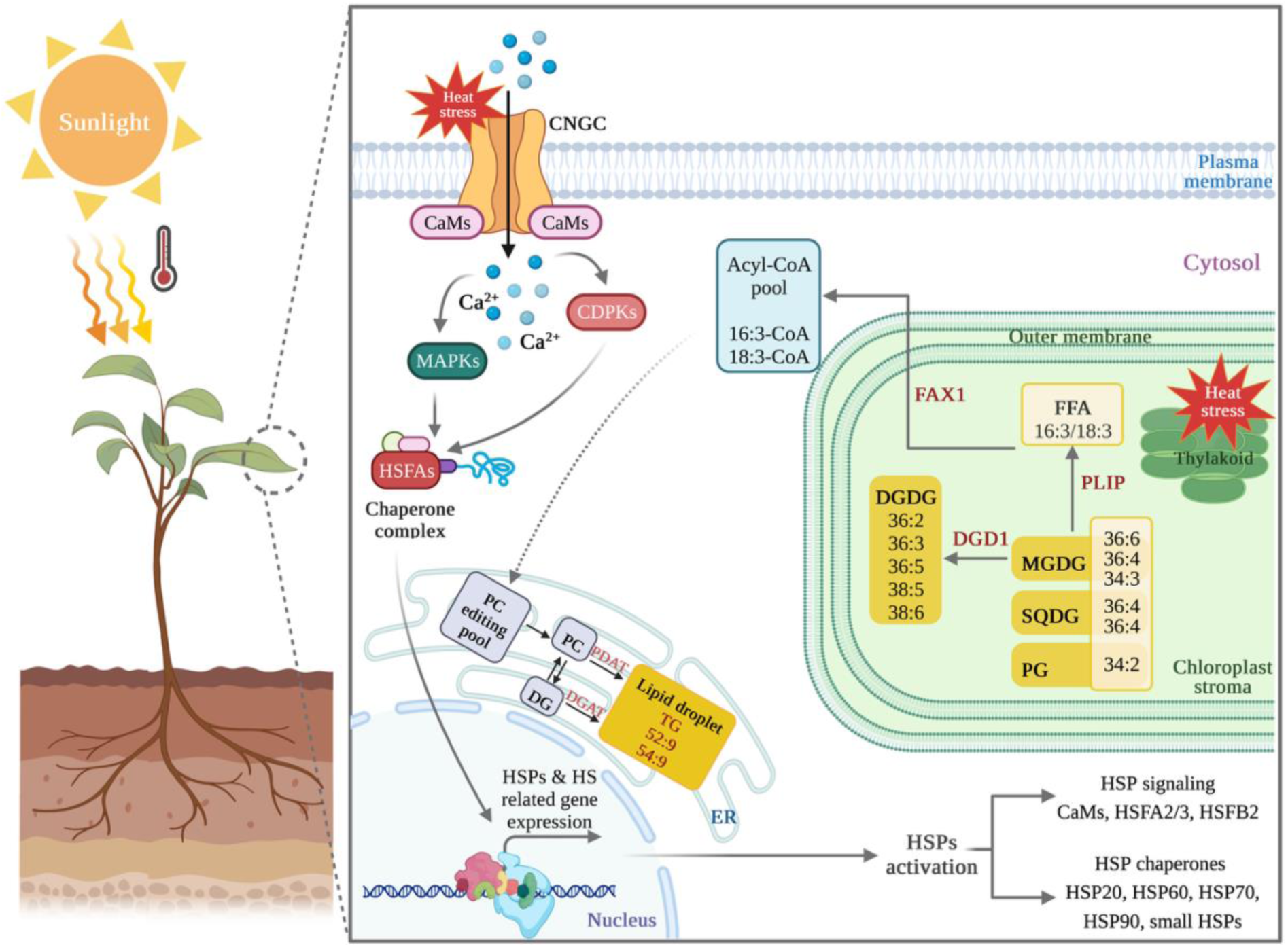
Proposed model of basal thermotolerance mechanisms in chia leaves under heat stress. Heat stress induces changes in gene expression and lipid composition in chia leaves. Heat-induced changes in membrane fluidity lead to Ca²⁺ influx into the cytosol via cyclic nucleotide-gated ion channels (CNGCs) by upregulating calmodulin (CaMs). This activates calcium-dependent protein kinases (CDPKs) and mitogen-activated protein kinases (MAPKs), which in turn induce heat shock transcription factors (HSFA and HSFB). These transcription factors translocate to the nucleus and upregulate heat shock proteins (HSP20, HSP60, HSP70, HSP90, and small HSPs). Concurrently, heat stress activates remodelling of thylakoid membrane lipids, including increased abundance of free fatty acid (FFA)-enriched monogalactosyldiacylglycerols (MGDGs). FFAs are released from MGDGs by PLASTID LIPASE1 (PLIP1), initiating lipid remodelling in response to heat stress. To mitigate FFA toxicity, 18:3-CoA and 16:3-CoA intermediates are transported to the endoplasmic reticulum (ER), where they are incorporated into phosphatidylcholines (PC) and subsequently converted into triacylglycerols (TGs). This process is supported by the heat-induced transcriptional upregulation of phospholipid diacylglycerol acyltransferase (PDAT) and diacylglycerol O-acyltransferase (DGAT) genes. Abbreviations: DG, diacylglycerol; PG, phosphatidylglycerol; DGDG, digalactosyldiacylglycerol; SQDG, sulfoquinovosyldiacylglycerol; FAX1, fatty acid export 1; DGD1, digalactosyldiacylglycerol synthase. Figure created with BioRender.com.

In chia, HSFs and chaperone-related genes were strongly upregulated during heat shock, consistent with patterns observed in heat-treated Arabidopsis seedlings (Wang et al., 2020). HSP transcripts in chia showed sustained up-regulation throughout both short-term and prolonged heat stress, highlighting their central role in protecting protein homeostasis across different stages of heat exposure. Although this mechanism is well-studied in model plants, it remains less explored in crops. A better understanding of HSF/HSP regulation in crops like chia may support breeding initiatives aimed at improving thermotolerance.

#### Transcriptomic regulation of lipid remodelling in chia under heat stress

Heat shock and prolonged heat stress triggered significant changes in the lipid composition of chia leaves. The most pronounced shift was observed in the TGs, particularly during heat shock, with several saturated lipid species showing increased abundance (Figure 5a; Supplemental Figure S7a). Such TG accumulation is a well-documented feature of heat-acclimated plant tissues, as observed in Arabidopsis (Higashi et al., 2015, Mueller et al., 2015) and wheat (Narayanan et al., 2016). In chia, the increases in both saturated and unsaturated TG species may reflect the involvement of different lipid remodelling pathways contributing to thermotolerance (Figure 6).

A shift toward more saturated FA species was most evident under heat shock, with TG species enriched in saturated FAs showing a marked increase in abundance (Supplemental Figures S7a & b). This change likely indicates decreased desaturase activity, a typical early response to heat stress that restricts membrane fluidity (Narayanan et al., 2016, Niu and Xiang, 2018). Increased membrane fluidity under heat stress can promote Ca²⁺ influx, which may contribute to cellular damage, such as apoptosis or necrosis (Conrard and Tyteca, 2019). In chia, this response coincided with the upregulation of HSFs and increased levels of saturated lipids (Supplemental Figures S7a & b). Similar observations have been reported in Arabidopsis (Tang et al., 2016, Shiva et al., 2020) and moss (*P. patens*) (Saidi et al., 2010). The increased production of stress-associated lipid species may support cellular tolerance and contribute to the adaptation of plants to high-temperature environments (Li et al., 2012, López-Marqués, 2021, Xie et al., 2021).

The accumulation of TGs in chia leaves under heat stress was associated with increased levels of both saturated and unsaturated species, including TG 52:9 (18:3/18:3/16:3), which rose 8-fold after heat shock and 16-fold after prolonged exposure (Supplemental Figures S7a & b). A similar pattern was reported in Arabidopsis (Mueller et al., 2015, Shiva et al., 2020), suggesting a potential general role for unsaturated TGs in mitigating long-term heat stress in plants. This accumulation may reflect a detoxification mechanism. During heat stress, polyunsaturated FAs such as 18:3, released from MGDGs during chloroplast membrane remodelling (Higashi et al., 2018), are transported to the ER, converted to 18:3-CoA (He et al., 2020), and incorporated into TGs by PDAT (Li-Beisson et al., 2013, Shiva et al., 2020). This response may involve the final steps of the Kennedy pathway, where PDAT or DGAT acylates DGs to form TGs (Slocombe et al., 2009, Fan et al., 2013, Tjellström et al., 2015). Our transcriptomic data show that both *PDAT* and *DGAT* were upregulated under heat stress (Supplemental Figure S10), consistent with similar observations in Arabidopsis, where *pdat1* mutants fail to accumulate TGs under elevated temperatures (Mueller et al., 2017). The return of TGs to control levels after heat treatment may indicate a role in transient lipid remodelling during heat stress adaptation in chia.

#### Brassinosteroids may contribute to thermotolerance in chia

Transcript and lipid levels in chia leaves returned to control values after a 24-hour recovery phase. Similar recovery without prior acclimation, known as basal thermotolerance, has been reported in several species, including spinach (Yan et al., 2016), Arabidopsis (Mueller et al., 2017, Serrano et al., 2019), grapevine (Venios et al., 2020), tomato (Fernández-Crespo et al., 2022), and rice (Samtani et al., 2022). This trait relies on sensing and release of signalling molecules (Chen et al., 2022, Song et al., 2022) that activate heat-responsive genes (Sajid et al., 2018, Shen et al., 2022, Yang et al., 2022). Nonetheless, a small number of DEGs and residual lipid changes remained after recovery in chia.

After recovery, the small number of remaining DEGs was mainly enriched for genes involved in brassinosteroid (BR) homeostasis. BRs are plant steroid hormones that regulate key developmental processes, including microtubule stabilisation in the cytoskeleton (Liu et al., 2018). In Arabidopsis, BR biosynthesis-deficient mutants exhibit reduced basal and acquired thermotolerance in response to heat stress (Luo et al., 2022). The upregulation of BR biosynthetic genes may support recovery and growth following heat stress in chia. However, to investigate an epigenetic thermal memory in chia, similar to what has been shown in Arabidopsis (Lämke et al., 2016, Shekhawat et al., 2022), a subsequent heat treatment would need to be applied to the offspring of heat-exposed plants.

## CONCLUSION

This study revealed that basal thermotolerance in chia involves the capacity to rapidly respond to heat shock, maintain homeostasis during prolonged heat exposure, and return to near-control levels after a 24-hour recovery period. In chia leaves, heat shock triggered the downregulation of Ca²⁺ signalling-related genes and the upregulation of HSFs, while prolonged heat stress maintained elevated levels of HSPs and restored Ca²⁺ metabolism gene expression to control levels. Lipid profiling revealed changes in TG composition, with the accumulation of both saturated and unsaturated TG species under heat stress. These changes were accompanied by the upregulation of *PDAT* and *DGAT* genes, suggesting an active lipid remodelling process potentially linked to FA detoxification. Although further experiments, such as repeated heat stress or exploring thermal memory in progenies, are needed to fully understand thermotolerance in chia, this study offers valuable insights into its molecular basis and highlights promising targets for improving heat resilience in chia and other oilseed crops through biotechnological and conventional breeding efforts.

## MATERIALS AND METHODS

### Plant materials and experimental design

Chia seeds (black commercial genotype) were supplied by the Chia Co. and the Northern Australia Crop Research Alliance Pty Ltd (NACRA) (Kununurra, Western Australia). Seeds were germinated in commercial garden soil (Osmocote^®^), and plants were grown at a temperature of 26/15 °C day/night, a photoperiod of 11.5 hr/12.5 hr light/dark with a relative humidity of 60–61% in a growth chamber. Four-week-old chia seedlings (56 homogeneous plants) were used for transcriptomic and lipidomic studies. The experiment was conducted in two temperature-controlled growth chambers: one for control and one for heat treatment. Leaf tissues were collected at four time points (T0, T1 (3 h), T2 (27 h), and T3 (24 h)) with eight biological replicates (Supplemental Figure S1).

To minimise circadian rhythm effects on molecular physiology, all samples were collected at 2:30 pm. Plants in both control and treatment conditions were watered daily, with special attention during heat stress to prevent wilting or drought stress. On day 1, T0 plants remained in the control chamber (26 °C day / 15 °C night). On day 2, heat-treated plants (T1.H, T2.H, and T3.R) were transferred to a separate chamber at 11:00 am and exposed to 38 °C day / 20°C night, while the controls (T1.C, T2.C, T3.C) remained in the original chamber. T1.H plants were sampled after 3 h of heat exposure. T2.H and T3.R plants were exposed to the same high temperatures for 27 h. On day 3, T3.R plants were returned to the control chamber for a 24 h recovery. Samples from T2.H and T3.R were collected at 2:30 pm on days 3 and 4, immediately frozen in liquid nitrogen, and stored at –80 °C until used.

### Transcriptomic analysis

#### RNA isolation and sequencing

Total RNA was extracted following the manufacturer’s instructions using the RNeasy Plant Mini Kit (Qiagen, Hilden, Germany). DNA contaminants were removed using the RNase-Free DNase Kit (Qiagen, Hilden, Germany). RNA purity was assessed using a NanoDrop^™^ spectrophotometer (Thermo Fisher Scientific, MA, USA), and concentrations were determined using the Qubit^™^ 3.0 Fluorometer and RNA High Sensitivity Assay Kit (Thermo Fisher Scientific). RNA integrity was confirmed using 1% agarose gel electrophoresis, and samples with a 260/280 ratio between 1.8 and 2.0 and RNA integrity number (RIN) > 7, were selected. Subsequently, 1 μg of total RNA was submitted to GENEWIZ (Suzhou, China) for RNAseq library preparation and sequencing.

RNA-seq libraries were prepared using the NEBNext Ultra RNA Library Prep Kit for Illumina (NEB, MA, USA). Poly(A) enrichment and rRNA removal were performed with Oligo-d(T) magnetic beads (NEBNext Poly(A) mRNA Magnetic Isolation Module). Fragmentation was conducted with NEBNext First Strand Synthesis Reaction Buffer (5X), followed by priming with NEBNext Random Hexamer Primers. Second-strand synthesis was carried out with ProtoScript II Reverse Transcriptase and NEBNext Second Strand Synthesis Enzyme Mix (NEB). Double-stranded cDNA was purified with AxyPrep MAG PCR Clean-up Kit (Axygen, CA, USA), and end-repair was completed using NEBNext End Prep Enzyme Mix (NEB). A-tailing was done with the NEBNext dA-Tailing Module (NEB), and adaptor ligation was conducted using Illumina-compatible adaptors and DNA ligase, following the NEBNext Ultra RNA Library Prep Kit protocol.

Adaptor-ligated cDNAs were size-selected using the AxyPrep MAG PCR Clean-up Kit (Axygen) to obtain fragments of approximately 360 bp (insert size ∼300 bp). The libraries underwent PCR amplification (11 cycles), were validated using an Agilent 2100 Bioanalyzer (Agilent Technologies, CA, USA), and were quantified using a Qubit^TM^ 2.0 Fluorometer (Invitrogen, CA, USA). Sequencing was conducted on an Illumina HiSeq X-ten platform in 2×150 bp paired-end mode, yielding 2,116,114,396 reads (∼635 Gb) across 56 RNA-seq libraries (Supplemental Figure S2 & Table S1).

#### Quality control and preprocessing of the raw RNAseq dataset

The quality of the raw RNA-seq reads was evaluated using FastQC v0.11.9 (Andrews, 2010). Trimmomatic v0.39 (Bolger et al., 2014) was used to remove low-quality bases and adaptor sequences using the following parameters: SLIDINGWINDOW:10:28, HEADCROP:12, MINLEN:136, and ILLUMINACLIP:TruSeq3-PE-2.fa:7:25:8:1:true. After preprocessing, 614 million clean reads were retained across 56 libraries, with each library yielding between 26 and 38 million reads. All samples met quality control criteria for downstream analysis.

#### RNAseq data analysis

Sequencing reads were aligned to the chia reference genome (Zare et al., 2024b) using STAR v2.7.10a (Dobin et al., 2013) (Table S2). Transcript expression levels were quantified with Cuffquant v2.2.1 (Trapnell et al., 2012), using the chia genome annotation file version 0.1 (Zare et al., 2024b). Transcript abundance was further calculated using FeatureCounts from the R\Subread package v2.0.3 (Liao et al., 2019).

#### Analysis of differentially expressed genes (DEGs)

Genes with zero expression across all time points were excluded from the analysis. Low-abundance genes were removed using the filterByExpr() function from the edgeR package v3.38.4 (Robinson et al., 2010), with the following parameters: min.count = 10, min.total.count = 15, large.n = 10, and min.prop = 0.7. To account for variation in sequencing depth, gene expression values were normalised by library size. Library size consistency was verified using the plotLibrarySizes() function in edgeR (Robinson et al., 2010). Normalisation efficiency was assessed using a mean-difference (MD) plot generated with the limma package v3.52.3 (Ritchie et al., 2015), which evaluated the impact of Trimmed Mean of M-values (TMM) normalisation. The distribution of gene log fold-changes (logFCs) was centred and symmetrical around zero, indicating that no systematic bias was introduced during RNA-seq library preparation or normalisation (Supplemental Figure S3).

#### Exploratory data analysis

To assess sample relationships, detect potential outliers, and evaluate group separation, hierarchical clustering and dimensionality reduction techniques were applied. Raw read counts were converted to counts per million (CPM). A distance matrix was computed using the “canberra” method, followed by hierarchical clustering with the “ward.D” linkage method. Principal component analysis (PCA) was visualised using the R/ggbiplot package v0.55 (Vu, 2011). Multidimensional scaling (MDS) plots were generated using the plotMDS() function from the limma package, with the default argument set to gene.selection = “pairwise”.

#### Differential expression modelling

Differential expression analysis was performed using the edgeR (Robinson et al., 2010) and limma-voom (Law et al., 2014) pipelines. In edgeR, a quasi-likelihood (QL) negative binomial generalised linear model (NB-GLM) was fitted using the glmQLFit() function, and QL F-tests were conducted via glmQLFTest() to identify DEGs. An empirical Bayes (EB) approach was applied, in which a mean-dependent trend was fitted to the raw QL dispersion estimates, shrinking them toward the trend to stabilise variance estimates and account for both technical and biological variability in gene expression measurements (Robinson et al., 2010, Chen et al., 2016, Lun et al., 2016).

To isolate treatment effects and account for time-dependent variation, logFCs were computed by subtracting the temporal component through comparisons with matched control groups. Specifically, heat shock at T1 (3 h heat treatment) was modelled as dT1H_T0 = (T1.H − T0.H) − (T1.C − T0.C); prolonged heat stress at T2 (an additional 24 h of heat exposure) as dT2H_T0 = (T2.H − T0.H) − (T2.C − T0.C); and recovery at T3 (24 h under optimal conditions) as dT3R_T0 = (T3.R − T0.H) − (T3.C − T0.C). Log_2_FC values were calculated using an average prior count of 2, and a common dispersion estimate of ∼0.03 was used for the NB model. DEGs were identified using the decideTests() function in the limma package, with method = “separate” and a false discovery rate (FDR) < 0.001. Mean–difference plots were generated with the plotMD() function in limma, and volcano plots were generated using the R/EnhancedVolcano package (Blighe and Lewis, 2018).

Both the edgeR and limma-voom methods identified largely overlapping sets of DEGs, with only minor differences (Supplemental Figure S4). However, edgeR employs an NB-GLM regression, which does not rely on the Poisson distribution assumptions inherent in limma-voom’s linear modelling framework. This distinction is particularly important in the context of RNA-seq data, where overdispersion is common, meaning that the variance exceeds the mean. Consequently, downstream analyses were based on the DEGs identified using edgeR.

#### K-means clustering and Gene Ontology enrichment analysis

K-means clustering was used to explore the response profiles of DEGs and their pairwise interactions, following methods adapted from the Parker Lab’s RNA-seq analysis workflow (Science Park Study Group, 2017). The regression coefficients of the fitted models for the contrasts of interest (excluding controls) were scaled and centred, then used for k-means clustering with k = 4. The optimal number of clusters was determined using the sum of squared errors, average silhouette width using cluster v2.1.4 (Maechler et al., 2021), and the Calinski– Harabasz index using vegan v2.6-4 (Oksanen et al., 2022). Genes with correlation scores < 0.8 were excluded. Heatmaps were generated using R/gplots v3.1.3 (Warnes et al., 2022). GO enrichment analysis was performed in ShinyGO v0.76.2 (Ge et al., 2020), with GO terms assigned to DEGs via Arabidopsis homologs at FDR ≤ 0.05, following Paril et al. (2023) and Zare et al. (2024b).

### Lipid analysis

#### Lipid extraction

Frozen plant material was ground using a pre-chilled mortar and pestle. Approximately 30 mg of chia leaf powder was used for lipid extraction following the protocol described by Kehelpannala et al. (2020). Eight biological replicates were prepared for each time point. For homogenisation, 400 μL of cold 2-propanol containing 0.01% butylated hydroxytoluene (BHT) and 20 μM d-cholesterol was added to each sample. Samples were homogenised using a TissueLyser II (Qiagen, USA) at 30 Hz for 3 min at –10 °C. They were then incubated at 75 °C for 15 min with constant shaking at 1400 rpm, followed by cooling to room temperature (25°C). Subsequently, 1.2 mL of a chloroform (CHCl₃)/methanol (MeOH)/water (30:41.5:3.5, v/v/v) mixture was added to each tube. Samples were shaken at 300 rpm and incubated for 24 h at 25 °C. Lipids were extracted from the upper phase following centrifugation at 13,000 rpm for 15 min at 25 °C. The extracts were dried using a rotary vacuum concentrator at room temperature without heating (Supplemental Figure S5). For quality control, pooled biological quality control (PBQC) samples were prepared by combining 10 μL aliquots from each lipid extract to assess reproducibility and account for biological variation.

#### Lipid analysis by liquid chromatography (LC-MS)

The dried lipid samples were reconstituted in 200 μL of butanol (BuOH)/MeOH (1:1, v/v) containing 10 mM ammonium formate (AF). Samples were incubated at 20 °C for 20 min with constant shaking at 1400 rpm, then centrifuged at 13,000 rpm for 15 min at 25 °C. LC-MS analysis was performed as described in Hu et al. (2008).

Briefly, samples were placed in an autosampler maintained at 12 °C. Lipid extracts (15 μL) were injected into an InfinityLab Poroshell 120 EC-C18 reversed-phase column (2.1 × 100 mm, 2.7 μm particle size; Agilent Technologies, CA, USA), operated at 55 °C using a Sciex Exion LC30AD ultra-high-performance liquid chromatography (UHPLC) system at a flow rate of 0.26 mL/min. Elution was performed over a 30-min binary gradient using eluent A [acetonitrile (ACN)/water (60:40, v/v) with 10 mM AF] and eluent B [isopropanol (IPP)/ACN (90:10, v/v) with 10 mM AF], as described by Kehelpannala et al. (2021).

Lipid analysis was performed on a Sciex TripleTOF^™^ 6600 quadrupole–quadrupole time–of– flight (QqTOF) mass spectrometer equipped with an automated calibrant delivery system (CDS) and a Turbo V^™^ dual-ion source [electrospray ionisation (ESI) and atmospheric pressure chemical ionisation (APCI)]. Data were acquired in positive ion mode using the Sequential Window Acquisition of All Theoretical Mass Spectra (SWATH-MS) method (Tsugawa et al., 2019) with the following parameters: MS1 mass range, 100–1700 m/z; SWATH scan range, 300–1700 m/z; MS/MS mass range, 100–1700 m/z; TOF-MS and TOF-MS/MS accumulation times of 50.0 ms and 10.0 ms, respectively; collision energy, +45 V; collision energy spread, 15 V; precursor ion isolation window, 15 Da; precursor window cycle time, 1042 ms. The ESI settings were as follows: source temperature, 250 °C; curtain gas, 35 psi; gas1 and gas2 at 35 psi and 25 psi, respectively; ion spray voltage floating (ISVF), 5500 V; and declustering potential (DP), +80 V. The APCI calibration solution was automatically delivered by the CDS every six samples for instrument calibration.

#### Data processing

Data were initially pre-processed by visually inspecting the raw files using PeakView software v2.2 (AB Sciex) to assess peak integrity. Raw data were converted to ABF format using the REIFYCS file converter and analysed with Mass Spectrometry–Data Independent Analysis (MS-DIAL) v2.24 (Tsugawa et al., 2015). Key processing parameters included an MS1 and MS2 tolerance of 0.01 Da and 0.05 Da, respectively; MS1 mass range, 300–1700 Da; and a minimum peak height of 1000 amplitude. For peak alignment to quality control samples, an MS1 tolerance of 0.015 Da and a retention time (RT) tolerance of 0.05 min were applied.

Lipid species were identified in positive ion mode using [M+H]+ and [M+NH₄]+ adducts. The MS-DIAL internal lipid database was used for feature annotation, with mass tolerances of 0.01 Da (MS1) and 0.05 Da (MS2) applied (Tsugawa et al., 2015). Default parameters were retained for all other SWATH-MS settings. For FC analysis, only lipid species with a coefficient of variation (CV) above 30% in quality control samples were included, provided they exhibited consistent RT patterns within lipid classes and MS/MS fragmentation profiles matched reference spectra in the MS-DIAL database.

#### Statistical analysis of lipid data

The identified lipid peak areas from MS-DIAL were analysed using Microsoft Excel (Microsoft, https://www.microsoft.com). Peak areas were normalised to the fresh weight of each sample (∼30 mg) to account for technical and biological variability. The data were then log-transformed, Pareto-scaled, and subjected to statistical analysis using MetaboAnalyst v5.0 (Pang et al., 2022) (www.metaboanalyst.ca/MetaboAnalyst). PCA was conducted to assess variability patterns across control and treatment groups. To detect potential outliers, hierarchical clustering was performed using Pearson’s correlation distance and Ward’s linkage method.

Significant differences in lipid abundances between treatment and control groups were identified using Student’s *t*-test within MetaboAnalyst. *P*-values were adjusted for multiple testing using the Benjamini–Hochberg method, with a significance threshold set at FDR-adjusted *P* < 0.001 (Chong and Xia, 2020). Fold changes (FC) were calculated based on the log ratio of lipid responses in heat-treated versus control samples. Lipid species showing both statistical significance (FDR-adjusted *P* < 0.001) and an FC > 1.5 were considered differentially abundant.

### Data statements

The associated RNA-seq data are available in the Sequence Read Archive (SRA) under accession number SRP588053 (BioProject: PRJNA1268670; BioSample: SAMN48760599). The chia (*Salvia hispanica*) genome assembly and annotation are available on the NCBI database under GenBank accession GCA_023119035.1 and RefSeq accession GCF_023119035.1.

## Acknowledgements

**TZ** was supported by the Australian Research Training Program (RTP) Scholarship and received additional support from the University of Melbourne Alfred Nicholas Fellowship, the Megan Klemm Postgraduate Research Scholarship, and the Norma Hilda Schuster (née Swift) Scholarship, all awarded by the University of Melbourne. **TZ** also acknowledges funding from Rosewood Research, which enabled the use of advanced transcriptomic and lipidomic platforms in this study. **BE** was supported by the 2020 Inaugural Botany Foundation Fellowship Award from the University of Melbourne Botany Foundation.

## Short legends for supporting information

### Supporting figures

**Figure S1.** Schematic overview of the experimental design to investigate heat stress responses in chia leaves.

**Figure S2.** Workflow for RNA isolation, RNA-seq library preparation, and transcriptomic analysis in chia leaves.

**Figure S3.** Mean-difference (MD) plots for all RNA-seq samples.

**Figure S4.** Evaluation of dispersion estimates for differential expression modelling.

**Figure S5.** Workflow of lipidomics analysis in chia leaves.

**Figure S6.** Venn diagrams showing the number of stress-related genes identified across experimental conditions.

**Figure S7.** Significantly altered lipid species in response to heat stress.

**Figure S8.** Heatmap of lipid abundance in chia leaves under heat shock (T1.H) versus control (T1.C) conditions.

**Figure S9.** Heatmap of lipid abundance in chia leaves under prolonged heat stress (T2.H) versus control (T2.C) conditions.

**Figure S10.** Temporal Expression of Lipid Biosynthesis Genes in Chia under Heat Stress.

### Supporting Tables

**Table S1.** Summary of sequencing quality metrics for raw RNA-seq reads across all libraries.

**Table S2.** Summary of RNA-seq read alignment statistics to the chia reference genome using STAR.

**Table S3.** Literature-derived lipid metabolism genes identified in chia.

**Table S4.** Differentially expressed lipid metabolism genes in chia under heat stress.

**Table S5.** Gene Ontology (GO) enrichment analysis of differentially expressed genes (DEGs) under heat shock in chia (T1.H vs. T0): Top 20 enriched biological processes.

**Table S6.** Gene Ontology (GO) enrichment analysis of differentially expressed genes (DEGs) under heat shock in chia (T1.H vs. T0): Top 20 enriched molecular functions.

**Table S7.** Gene Ontology (GO) enrichment analysis of differentially expressed genes (DEGs) under heat shock in chia (T1.H vs. T0): Top 20 enriched cellular components.

**Table S8.** Gene Ontology (GO) enrichment analysis of differentially expressed genes (DEGs) under prolonged heat stress in chia (T2.H vs. T0): Enriched biological processes.

**Table S9.** Gene Ontology (GO) enrichment analysis of DEGs under prolonged heat stress (T2.H vs T0), highlighting the enriched molecular functions (FDR <0.05).

**Table S10.** Gene Ontology (GO) enrichment analysis of DEGs under prolonged heat stress (T2.H vs T0), highlighting the enriched cellular components (FDR <0.05).

**Table S11.** Annotated lipid species and their abundance levels in chia leaves in response to heat.

## References

1. Abbas, A., Shah, A. N., Shah, A. A., Nadeem, M. A., Alsaleh, A., Javed, T., Alotaibi, S. S. & Abdelsalam, N. R. 2022. Genome-wide analysis of invertase gene family, and expression profiling under abiotic stress conditions in potato. Biology, 11(4), 539.

2. Abdelrahman, M., Ishii, T., El-Sayed, M. & Tran, L.-S. P. 2020. Heat sensing and lipid reprograming as a signaling switch for heat stress responses in wheat. Plant and Cell Physiology, 61(8), 1399–1407.

3. Alejo-Jacuinde, G., Nájera-González, H.-R., Chávez Montes, R. A., Gutierrez Reyes, C. D., Barragán-Rosillo, A. C., Perez Sanchez, B., Mechref, Y., López-Arredondo, D., Yong-Villalobos, L. & Herrera-Estrella, L. 2023. Multi-omic analyses reveal the unique properties of chia (*Salvia hispanica*) seed metabolism. Communications Biology, 6(1), 820.

4. Andrási, N., Pettkó-Szandtner, A. & Szabados, L. 2021. Diversity of plant heat shock factors: regulation, interactions, and functions. Journal of Experimental Botany, 72(5), 1558–1575.

5. Andrews, S. 2010. FastQC: a quality control tool for high throughput sequence data.

6. Arif, M., Haroon, M., Nawaz, A. F., Abbas, H., Xu, R. & Li, L. 2025. Enhancing wheat resilience: biotechnological advances in combating heat stress and environmental challenges. Plant Molecular Biology, 115(2), 41.

7. Bibi, F. & Rahman, A. 2023. An overview of climate change impacts on agriculture and their mitigation strategies. Agriculture, 13(8), 1508.

8. Blighe, K. & Lewis, M. 2018. EnhancedVolcano: Publication-ready volcano plots with enhanced colouring and labeling. R package version, 1(0).

9. Bolger, A. M., Lohse, M. & Usadel, B. 2014. Trimmomatic: a flexible trimmer for Illumina sequence data. Bioinformatics, 30(15), 2114–2120.

10. Bourgine, B. & Guihur, A. 2021. Heat shock signaling in land plants: from plasma membrane sensing to the transcription of small heat shock proteins. Frontiers in plant science, 12(710801.

11. Chaudhry, S. & Sidhu, G. P. S. 2022. Climate change regulated abiotic stress mechanisms in plants: a comprehensive review. Plant Cell Reports, 41(1), 1–31.

12. Chen, Y., Lun, A. T. & Smyth, G. K. 2016. From reads to genes to pathways: differential expression analysis of RNA-Seq experiments using Rsubread and the edgeR quasi-likelihood pipeline. F1000Research, 5(1438.

13. Chen, Z., Galli, M. & Gallavotti, A. 2022. Mechanisms of temperature-regulated growth and thermotolerance in crop species. Current opinion in plant biology, 65(102134.

14. Chong, J. & Xia, J. 2020. Using MetaboAnalyst 4.0 for metabolomics data analysis, interpretation, and integration with other omics data. Computational Methods and Data Analysis for Metabolomics, 337–360.

15. Conrard, L. & Tyteca, D. 2019. Regulation of membrane calcium transport proteins by the surrounding lipid environment. Biomolecules, 9(10), 513.

16. Crawford, F., Deards, B., Moir, B. & Thompson, N. 2012. Human consumption of hemp seed: prospects for Australian production. Canberra: Australian Bureau of Agricultural and Resource Economics and Sciences.

17. Cui, Y., Lu, S., Li, Z., Cheng, J., Hu, P., Zhu, T., Wang, X., Jin, M., Wang, X. & Li, L. 2020. CYCLIC NUCLEOTIDE-GATED ION CHANNELs 14 and 16 promote tolerance to heat and chilling in rice. Plant Physiology, 183(4), 1794–1808.

18. Ding, Y. & Yang, S. 2022. Surviving and thriving: How plants perceive and respond to temperature stress. Developmental Cell, 57(8), 947–958.

19. Djanaguiraman, M., Boyle, D., Welti, R., Jagadish, S. & Prasad, P. 2018. Decreased photosynthetic rate under high temperature in wheat is due to lipid desaturation, oxidation, acylation, and damage of organelles. BMC plant biology, 18(1-17.

20. Dobin, A., Davis, C. A., Schlesinger, F., Drenkow, J., Zaleski, C., Jha, S., Batut, P., Chaisson, M. & Gingeras, T. R. 2013. STAR: ultrafast universal RNA-seq aligner. Bioinformatics, 29(1), 15–21.

21. Dubrovina, A. S., Kiselev, K. V., Khristenko, V. S. & Aleynova, O. A. 2017. The calcium-dependent protein kinase gene VaCPK29 is involved in grapevine responses to heat and osmotic stresses. Plant Growth Regulation, 82(79-89.

22. Edel, K. H. & Kudla, J. 2016. Integration of calcium and ABA signaling. Current opinion in plant biology, 33(83-91.

23. Elzanati, O., Mouzeyar, S. & Roche, J. 2020. Dynamics of the transcriptome response to heat in the moss, *Physcomitrella patens*. International journal of molecular sciences, 21(4), 1512.

24. Fan, J., Yan, C., Zhang, X. & Xu, C. 2013. Dual role for phospholipid: diacylglycerol acyltransferase: enhancing fatty acid synthesis and diverting fatty acids from membrane lipids to triacylglycerol in Arabidopsis leaves. The Plant Cell, 25(9), 3506–3518.

25. Fernández-Crespo, E., Liu-Xu, L., Albert-Sidro, C., Scalschi, L., Llorens, E., González-Hernández, A. I., Crespo, O., Gonzalez-Bosch, C., Camañes, G. & García-Agustín, P. 2022. Exploiting tomato genotypes to understand heat stress tolerance. Plants, 11(22), 3170.

26. Finka, A., Cuendet, A. F. H., Maathuis, F. J., Saidi, Y. & Goloubinoff, P. 2012. Plasma membrane cyclic nucleotide gated calcium channels control land plant thermal sensing and acquired thermotolerance. The Plant Cell, 24(8), 3333–3348.

27. Friedrich, T., Oberkofler, V., Trindade, I., Altmann, S., Brzezinka, K., Lämke, J., Gorka, M., Kappel, C., Sokolowska, E. & Skirycz, A. 2021. Heteromeric HSFA2/HSFA3 complexes drive transcriptional memory after heat stress in Arabidopsis. Nature communications, 12(1), 3426.

28. Gao, C., Lu, S., Zhou, R., Wang, Z., Li, Y., Fang, H., Wang, B., Chen, M. & Cao, Y. 2022. The OsCBL8–OsCIPK17 module regulates seedling growth and confers resistance to heat and drought in rice. International Journal of Molecular Sciences, 23(20), 12451.

29. Gao, F., Han, X., Wu, J., Zheng, S., Shang, Z., Sun, D., Zhou, R. & Li, B. 2012. A heat-activated calcium-permeable channel–Arabidopsis cyclic nucleotide-gated ion channel 6–is involved in heat shock responses. The Plant Journal, 70(6), 1056–1069.

30. Ge, S. X., Jung, D. & Yao, R. 2020. ShinyGO: a graphical gene-set enrichment tool for animals and plants. Bioinformatics, 36(8), 2628–2629.

31. Ghosh, S., Bheri, M., Bisht, D. & Pandey, G. K. 2022. Calcium signaling and transport machinery: Potential for development of stress tolerance in plants. Current Plant Biology, 29(1), 100235.

32. Gupta, P., Geniza, M., Naithani, S., Phillips, J. L., Haq, E. & Jaiswal, P. 2021. Chia (*Salvia hispanica*) gene expression atlas elucidates dynamic spatio-temporal changes associated with plant growth and development. Frontiers in Plant Science, 12(667678.

33. Haider, S., Iqbal, J., Naseer, S., Yaseen, T., Shaukat, M., Bibi, H., Ahmad, Y., Daud, H., Abbasi, N. L. & Mahmood, T. 2021. Molecular mechanisms of plant tolerance to heat stress: current landscape and future perspectives. Plant Cell Reports, 1–25.

34. Haider, S., Raza, A., Iqbal, J., Shaukat, M. & Mahmood, T. 2022. Analyzing the regulatory role of heat shock transcription factors in plant heat stress tolerance: A brief appraisal. Molecular Biology Reports, 49(6), 5771–5785.

35. He, M., Qin, C.-X., Wang, X. & Ding, N.-Z. 2020. Plant unsaturated fatty acids: biosynthesis and regulation. Frontiers in Plant Science, 11(390.

36. Higashi, Y., Okazaki, Y., Myouga, F., Shinozaki, K. & Saito, K. 2015. Landscape of the lipidome and transcriptome under heat stress in *Arabidopsis thaliana*. Scientific reports, 5(1), 10533.

37. Higashi, Y., Okazaki, Y., Takano, K., Myouga, F., Shinozaki, K., Knoch, E., Fukushima, A. & Saito, K. 2018. HEAT INDUCIBLE LIPASE1 remodels chloroplastic monogalactosyldiacylglycerol by liberating α-linolenic acid in Arabidopsis leaves under heat stress. The Plant Cell, 30(8), 1887–1905.

38. Higashi, Y. & Saito, K. 2019. Lipidomic studies of membrane glycerolipids in plant leaves under heat stress. Progress in lipid research, 75(100990.

39. Hu, C., Van Dommelen, J., Van Der Heijden, R., Spijksma, G., Reijmers, T. H., Wang, M., Slee, E., Lu, X., Xu, G. & Van Der Greef, J. 2008. RPLC-ion-trap-FTMS method for lipid profiling of plasma: method validation and application to p53 mutant mouse model. Journal of proteome research, 7(11), 4982–4991.

40. Hu, X. J., Chen, D., Lynne Mclntyre, C., Fernanda Dreccer, M., Zhang, Z. B., Drenth, J., Kalaipandian, S., Chang, H. & Xue, G. P. 2018. Heat shock factor C2a serves as a proactive mechanism for heat protection in developing grains in wheat via an ABA-mediated regulatory pathway. Plant, cell & environment, 41(1), 79–98.

41. Jagadish, S. K., Way, D. A. & Sharkey, T. D. 2021. Plant heat stress: Concepts directing future research. Plant, cell & environment, 44(7), 1992–2005.

42. Jaimes-Miranda, F. & Chávez Montes, R. A. 2020. The plant MBF1 protein family: a bridge between stress and transcription. Journal of experimental botany, 71(6), 1782–1791.

43. Jarratt-Barnham, E., Wang, L., Ning, Y. & Davies, J. M. 2021. The complex story of plant cyclic nucleotide-gated channels. International Journal of Molecular Sciences, 22(2), 874.

44. Kan, Y., Mu, X.-R., Gao, J., Lin, H.-X. & Lin, Y. 2023. The molecular basis of heat stress responses in plants. Molecular Plant, 16(10), 1612–1634.

45. Kang, X., Zhao, L. & Liu, X. 2023. Calcium signaling and the response to heat shock in crop plants. International Journal of Molecular Sciences, 25(1), 324.

46. Kehelpannala, C., Rupasinghe, T., Pasha, A., Esteban, E., Hennessy, T., Bradley, D., Ebert, B., Provart, N. J. & Roessner, U. 2021. An Arabidopsis lipid map reveals differences between tissues and dynamic changes throughout development. The Plant Journal, 107(1), 287–302.

47. Kehelpannala, C., Rupasinghe, T. W., Hennessy, T., Bradley, D., Ebert, B. & Roessner, U. 2020. A comprehensive comparison of four methods for extracting lipids from Arabidopsis tissues. Plant Methods, 16(1-16.

48. Kumar, P., Paul, D., Jhajhriya, S., Kumar, R., Dutta, S., Siwach, P. & Das, S. 2024. Understanding heat-shock proteins’ abundance and pivotal function under multiple abiotic stresses. Journal of Plant Biochemistry and Biotechnology, 1–22.

49. Lämke, J., Brzezinka, K., Altmann, S. & Bäurle, I. 2016. A hit-and-run heat shock factor governs sustained histone methylation and transcriptional stress memory. The EMBO journal, 35(2), 162–175.

50. Law, C. W., Chen, Y., Shi, W. & Smyth, G. K. 2014. voom: Precision weights unlock linear model analysis tools for RNA-seq read counts. Genome biology, 15(1-17.

51. Lee, H., Calvin, K., Dasgupta, D., Krinner, G., Mukherji, A., Thorne, P., Trisos, C., Romero, J., Aldunce, P. & Ruane, A. C. 2024. Climate Change 2023 Synthesis Report: Summary for Policymakers. The Intergovernmental Panel on Climate Change.

52. Li, X., Moellering, E. R., Liu, B., Johnny, C., Fedewa, M., Sears, B. B., Kuo, M.-H. & Benning, C. 2012. A galactoglycerolipid lipase is required for triacylglycerol accumulation and survival following nitrogen deprivation in Chlamydomonas reinhardtii. The Plant Cell, 24(11), 4670–4686.

53. Li, Y., Liu, Y., Jin, L. & Peng, R. 2022. Crosstalk between Ca^2+^ and other regulators assists plants in responding to abiotic stress. Plants, 11(10), 1351.

54. Li, Z. Q., Xing, W., Luo, P., Zhang, F. J., Jin, X. L. & Zhang, M. H. 2019. Comparative transcriptome analysis of *Rosa chinensis* ‘Slater’s crimson China’provides insights into the crucial factors and signaling pathways in heat stress response. Plant Physiology and Biochemistry, 142(312-331.

55. Li-Beisson, Y., Shorrosh, B., Beisson, F., Andersson, M. X., Arondel, V., Bates, P. D., Baud, S., Bird, D., DeBono, A. & Durrett, T. P. 2013. Acyl-lipid metabolism. The Arabidopsis book/American Society of Plant Biologists, 11(e0161.

56. Liao, Y., Smyth, G. K. & Shi, W. 2019. The R package Rsubread is easier, faster, cheaper and better for alignment and quantification of RNA sequencing reads. Nucleic acids research, 47(8), e47–e47.

57. Lippmann, R., Babben, S., Menger, A., Delker, C. & Quint, M. 2019. Development of wild and cultivated plants under global warming conditions. Current Biology, 29(24), R1326–R1338.

58. Liu, G.-T., Wang, J.-F., Cramer, G., Dai, Z.-W., Duan, W., Xu, H.-G., Wu, B.-H., Fan, P.-G., Wang, L.-J. & Li, S.-H. 2012. Transcriptomic analysis of grape (*Vitis vinifera* L.) leaves during and after recovery from heat stress. BMC plant biology, 12(1-10.

59. Liu, H.-T., Li, B., Shang, Z.-L., Li, X.-Z., Mu, R.-L., Sun, D.-Y. & Zhou, R.-G. 2003. Calmodulin is involved in heat shock signal transduction in wheat. Plant physiology, 132(3), 1186–1195.

60. Liu, X., Yang, Q., Wang, Y., Wang, L., Fu, Y. & Wang, X. 2018. Brassinosteroids regulate pavement cell growth by mediating BIN2-induced microtubule stabilization. Journal of experimental botany, 69(5), 1037–1049.

61. López-Marqués, R. L. 2021. Lipid flippases as key players in plant adaptation to their environment. Nature Plants, 7(9), 1188–1199.

62. Lu, J., Xu, Y., Wang, J., Singer, S. D. & Chen, G. 2020. The role of triacylglycerol in plant stress response. Plants, 9(4), 472.

63. Lun, A. T., Chen, Y. & Smyth, G. K. 2016. It’s DE-licious: a recipe for differential expression analyses of RNA-seq experiments using quasi-likelihood methods in edgeR. Statistical genomics: methods and protocols, 391–416.

64. Luo, J., Jiang, J., Sun, S. & Wang, X. 2022. Brassinosteroids promote thermotolerance through releasing BIN2-mediated phosphorylation and suppression of HsfA1 transcription factors in Arabidopsis. Plant communications, 3(6).

65. Maechler, M., Rousseeuw, P., Struyf, A., Hubert, M. & Hornik, K. 2021. cluster: cluster analysis basics and extensions. R package version 2.1. 4—For new features, see the ‘Changelog’file (in the package source). R Package Version R Foundation for Statistical Computing: Vienna, Austria, 1(56.

66. Mesihovic, A., Ullrich, S., Rosenkranz, R. R., Gebhardt, P., Bublak, D., Eich, H., Weber, D., Berberich, T., Scharf, K.-D. & Schleiff, E. 2022. HsfA7 coordinates the transition from mild to strong heat stress response by controlling the activity of the master regulator HsfA1a in tomato. Cell reports, 38(2).

67. Mishra, D. C., Arora, D., Kumar, R. R., Goswami, S., Varshney, S., Budhlakoti, N., Kumar, S., Chaturvedi, K. K., Sharma, A. & Chinnusamy, V. 2021. Weighted gene co-expression analysis for identification of key genes regulating heat stress in wheat. Cereal Research Communications, 49(73-81.

68. Mishra, S. K., Tripp, J., Winkelhaus, S., Tschiersch, B., Theres, K., Nover, L. & Scharf, K.-D. 2002. In the complex family of heat stress transcription factors, HsfA1 has a unique role as master regulator of thermotolerance in tomato. Genes & development, 16(12), 1555–1567.

69. Mondal, R., Madhurya, K., Saha, P., Chattopadhyay, S., Antony, S., Kumar, A., Roy, S. & Roy, D. 2022. Expression profile, transcriptional and post-transcriptional regulation of genes involved in hydrogen sulphide metabolism connecting the balance between development and stress adaptation in plants: a data-mining bioinformatics approach. Plant Biology, 24(4), 602–617.

70. Mueller, S. P., Krause, D. M., Mueller, M. J. & Fekete, A. 2015. Accumulation of extra-chloroplastic triacylglycerols in Arabidopsis seedlings during heat acclimation. Journal of Experimental Botany, 66(15), 4517–4526.

71. Mueller, S. P., Unger, M., Guender, L., Fekete, A. & Mueller, M. J. 2017. Phospholipid: diacylglycerol acyltransferase-mediated triacylglyerol synthesis augments basal thermotolerance. Plant Physiology, 175(1), 486–497.

72. Narayanan, S., Tamura, P. J., Roth, M. R., Prasad, P. V. & Welti, R. 2016. Wheat leaf lipids during heat stress: I. High day and night temperatures result in major lipid alterations. Plant, cell & environment, 39(4), 787–803.

73. Niu, Y. & Xiang, Y. 2018. An overview of biomembrane functions in plant responses to high-temperature stress. Frontiers in plant science, 9(915.

74. Oksanen, J., Simpson, G. L., Blanchet, F. G., Kindt, R., Legendre, P., Minchin, P. R., O’Hara, R., Solymos, P., Stevens, M. & Szoecs, E. 2022. vegan: Community ecology package (2.6-4). CRAN.

75. Oranab, S., Ghaffar, A., Ahmad, A., Pasha, M., Munir, B., Arif, S., Ishaq, S., Mahfooz, S., Kousar, R. & Zakia, S. 2023. Genome-wide analysis of cyclic nucleotide-gated ion channels (CNGCS) of *Arabidopsis thaliana* under abiotic stresses.

76. Pan, Y., Chai, X., Gao, Q., Zhou, L., Zhang, S., Li, L. & Luan, S. 2019. Dynamic interactions of plant CNGC subunits and calmodulins drive oscillatory Ca^2+^ channel activities. Developmental Cell, 48(5), 710–725. e5.

77. Pang, Z., Zhou, G., Ewald, J., Chang, L., Hacariz, O., Basu, N. & Xia, J. 2022. Using MetaboAnalyst 5.0 for LC–HRMS spectra processing, multi-omics integration and covariate adjustment of global metabolomics data. Nature Protocols, 17(8), 1735–1761.

78. Paril, J., Zare, T. & Fournier-Level, A. 2023. Compare_Genomes: A Comparative Genomics Workflow to Streamline the Analysis of Evolutionary Divergence Across Eukaryotic Genomes. Current Protocols, 3(8), e876.

79. Park, C.-J. & Shin, R. 2022. Calcium channels and transporters: Roles in response to biotic and abiotic stresses. Frontiers in Plant Science, 13(964059.

80. Peláez, P., Orona-Tamayo, D., Montes-Hernández, S., Valverde, M. E., Paredes-López, O. & Cibrián-Jaramillo, A. 2019. Comparative transcriptome analysis of cultivated and wild seeds of *Salvia hispanica* (chia). Scientific reports, 9(1), 9761.

81. Poonia, A. K., Mishra, S. K., Sirohi, P., Chaudhary, R., Kanwar, M., Germain, H. & Chauhan, H. 2020. Overexpression of wheat transcription factor (TaHsfA6b) provides thermotolerance in barley. Planta, 252(1-14.

82. Qin, D., Wu, H., Peng, H., Yao, Y., Ni, Z., Li, Z., Zhou, C. & Sun, Q. 2008. Heat stress-responsive transcriptome analysis in heat susceptible and tolerant wheat (*Triticum aestivum* L.) by using Wheat Genome Array. BMC genomics, 9(1-19.

83. Quicke, A. 2019. Heatwatch: Extreme heat in the Kimberley. Canberra: The Australia Institute.

84. Rahmati Ishka, M., Brown, E., Weigand, C., Tillett, R. L., Schlauch, K. A., Miller, G. & Harper, J. F. 2018. A comparison of heat-stress transcriptome changes between wild-type Arabidopsis pollen and a heat-sensitive mutant harboring a knockout of cyclic nucleotide-gated cation channel 16 (cngc16). BMC genomics, 19(1-19.

85. Rawsthorne, S. 2002. Carbon flux and fatty acid synthesis in plants. Progress in Lipid Research, 41(2), 182–196.

86. Ritchie, M. E., Phipson, B., Wu, D., Hu, Y., Law, C. W., Shi, W. & Smyth, G. K. 2015. limma powers differential expression analyses for RNA-sequencing and microarray studies. Nucleic Acids Research, 43(7), e47–e47.

87. Robinson, M. D., McCarthy, D. J. & Smyth, G. K. 2010. edgeR: a Bioconductor package for differential expression analysis of digital gene expression data. Bioinformatics, 26(1), 139–140.

88. Rodríguez, M. E., Poza-Viejo, L., Maestro-Gaitán, I., Schneider-Teixeira, A., Deladino, L., Ixtaina, V. & Reguera, M. 2024. Shotgun proteomics profiling of chia seeds (*Salvia hispanica* L.) reveals genotypic differential responses to viability loss. Frontiers in Plant Science, 15(1441234.

89. Saidi, Y., Finka, A., Muriset, M., Bromberg, Z., Weiss, Y. G., Maathuis, F. J. & Goloubinoff, P. 2009. The heat shock response in moss plants is regulated by specific calcium-permeable channels in the plasma membrane. The Plant Cell, 21(9), 2829–2843.

90. Saidi, Y., Peter, M., Finka, A., Cicekli, C., Vigh, L. & Goloubinoff, P. 2010. Membrane lipid composition affects plant heat sensing and modulates Ca^2+^-dependent heat shock response. Plant Signaling & Behavior, 5(12), 1530–1533.

91. Sajid, M., Rashid, B., Ali, Q. & Husnain, T. 2018. Mechanisms of heat sensing and responses in plants. It is not all about Ca^2+^ ions. Biologia Plantarum, 62(409-420.

92. Samtani, H., Sharma, A. & Khurana, P. 2022. Wheat ocs-element binding factor 1 enhances thermotolerance by modulating the heat stress response pathway. Frontiers in Plant Science, 13(914363.

93. Samtani, H., Sharma, A. & Khurana, P. 2023. Ectopic overexpression of TaHsfA5 promotes thermomorphogenesis in *Arabidopsis thaliana* and thermotolerance in *Oryza sativa*. Plant Molecular Biology, 112(4), 225–243.

94. Sato, H., Mizoi, J., Shinozaki, K. & Yamaguchi-Shinozaki, K. 2024. Complex plant responses to drought and heat stress under climate change. The Plant Journal, 117(6), 1873–1892.

95. Science Park Study Group, S. 2017. 08 Cluster analysis – Introduction to RNA-seq [Online]. Parker Lab, University of Cambridge. Available: https://scienceparkstudygroup.github.io/rna-seq-lesson/08-cluster-analysis/index.html [Accessed 15 August 2022].

96. Serrano, N., Ling, Y., Bahieldin, A. & Mahfouz, M. M. 2019. Thermopriming reprograms metabolic homeostasis to confer heat tolerance. Scientific Reports, 9(1), 181.

97. Shekhawat, K., Almeida-Trapp, M., García-Ramírez, G. X. & Hirt, H. 2022. Beat the heat: plant-and microbe-mediated strategies for crop thermotolerance. Trends in Plant Science, 27(8), 802–813.

98. Shen, T.-T., Wang, L., Shang, C.-H., Zhen, Y.-C., Fang, Y.-L., Wei, L.-L., Zhou, T., Bai, J.-T. & Li, B. 2022. The Arabidopsis J-protein AtDjC5 facilitates thermotolerance likely by aiding in the ER stress response. International Journal of Molecular Sciences, 23(21), 13134.

99. Shiva, S., Samarakoon, T., Lowe, K. A., Roach, C., Vu, H. S., Colter, M., Porras, H., Hwang, C., Roth, M. R. & Tamura, P. 2020. Leaf lipid alterations in response to heat stress of *Arabidopsis thaliana*. Plants, 9(7), 845.

100. Singh, K. & Chandra, A. 2021. DREBs-potential transcription factors involve in combating abiotic stress tolerance in plants. Biologia, 76(10), 3043–3055.

101. Slocombe, S. P., Cornah, J., Pinfield-Wells, H., Soady, K., Zhang, Q., Gilday, A., Dyer, J. M. & Graham, I. A. 2009. Oil accumulation in leaves directed by modification of fatty acid breakdown and lipid synthesis pathways. Plant Biotechnology Journal, 7(7), 694–703.

102. Song, Z. T., Chen, X. J., Luo, L., Yu, F., Liu, J. X. & Han, J. J. 2022. UBA domain protein SUF1 interacts with NatA-complex subunit NAA15 to regulate thermotolerance in Arabidopsis. Journal of Integrative Plant Biology, 64(7), 1297–1302.

103. Sreedhar, R. V., Kumari, P., Rupwate, S. D., Rajasekharan, R. & Srinivasan, M. 2015. Exploring triacylglycerol biosynthetic pathway in developing seeds of Chia (*Salvia hispanica* L.): a transcriptomic approach. PloS one, 10(4), e0123580.

104. Suri, S. S. & Dhindsa, R. S. 2008. A heat-activated MAP kinase (HAMK) as a mediator of heat shock response in tobacco cells. *Plant*, Cell & Environment, 31(2), 218–226.

105. Swindell, W. R., Huebner, M. & Weber, A. P. 2007. Transcriptional profiling of Arabidopsis heat shock proteins and transcription factors reveals extensive overlap between heat and non-heat stress response pathways. BMC Genomics, 8(1-15.

106. Tang, T., Liu, P., Zheng, G. & Li, W. 2016. Two phases of response to long-term moderate heat: Variation in thermotolerance between *Arabidopsis thaliana* and its relative *Arabis paniculata*. Phytochemistry, 122(81-90.

107. Tjellström, H., Strawsine, M. & Ohlrogge, J. B. 2015. Tracking synthesis and turnover of triacylglycerol in leaves. Journal of Experimental Botany, 66(5), 1453–1461.

108. Trapnell, C., Roberts, A., Goff, L., Pertea, G., Kim, D., Kelley, D. R., Pimentel, H., Salzberg, S. L., Rinn, J. L. & Pachter, L. 2012. Differential gene and transcript expression analysis of RNA-seq experiments with TopHat and Cufflinks. Nature Protocols, 7(3), 562–578.

109. Tsugawa, H., Cajka, T., Kind, T., Ma, Y., Higgins, B., Ikeda, K., Kanazawa, M., VanderGheynst, J., Fiehn, O. & Arita, M. 2015. MS-DIAL: data-independent MS/MS deconvolution for comprehensive metabolome analysis. Nature Methods, 12(6), 523–526.

110. Tsugawa, H., Hanada, A., Ikeda, K., Isobe, Y., Senoo, Y. & Senoo, M. 2019. High-Throughput Lipid Profiling with SWATH Aquisition and MS-Dial. SCIEX: Framingham, MA, USA, 1–5.

111. Venios, X., Korkas, E., Nisiotou, A. & Banilas, G. 2020. Grapevine responses to heat stress and global warming. Plants, 9(12), 1754.

112. Vu, V. Q. 2011. ggbiplot: A ggplot2 based biplot. R package version 0.55, 755(

113. Wang, H. & Xie, Z. 2024. Cullin-conciliated regulation of plant immune responses: Implications for sustainable crop protection. Plants, 13(21), 2997.

114. Wang, L., Lee, M., Sun, F., Song, Z., Yang, Z. & Yue, G. H. 2022. A chromosome-level genome assembly of chia provides insights into high omega-3 content and coat color variation of its seeds. Plant Communications, 3(4).

115. Wang, L., Ma, K.-B., Lu, Z.-G., Ren, S.-X., Jiang, H.-R., Cui, J.-W., Chen, G., Teng, N.-J., Lam, H.-M. & Jin, B. 2020. Differential physiological, transcriptomic and metabolomic responses of Arabidopsis leaves under prolonged warming and heat shock. BMC Plant Biology, 20(1-15.

116. Warnes, G. R., Bolker, B., Bonebakker, L., Gentleman, R., Huber, W., Liaw, A., Lumley, T., Maechler, M., Magnusson, A. & Moeller, S. 2022. gplots: Various R programming tools for plotting data (R package version 3.1.3) [Online]. CRAN. Available: https://CRAN.R-project.org/package=gplots [Accessed 17 Aug 2022].

117. Weigand, C., Kim, S.-H., Brown, E., Medina, E., Mares III, M., Miller, G., Harper, J. F. & Choi, W.-G. 2021. A ratiometric calcium reporter CGf reveals calcium dynamics both in the single cell and whole plant levels under heat stress. Frontiers in Plant Science, 12(777975.

118. Wimberley, J., Cahill, J. & Atamian, H. S. 2020. De novo sequencing and analysis of Salvia hispanica tissue-specific transcriptome and identification of genes involved in terpenoid biosynthesis. Plants, 9(3), 405.

119. Wu, H.-C., Luo, D.-L., Vignols, F. & Jinn, T.-L. 2012. Heat shock-induced biphasic Ca2+ signature and OsCaM1-1 nuclear localization mediate downstream signalling in acquisition of thermotolerance in rice (*Oryza sativa* L.).

120. Xie, L.-J., Zhou, Y., Chen, Q.-F. & Xiao, S. 2021. New insights into the role of lipids in plant hypoxia responses. Progress in Lipid Research, 81(101072.

121. Xu, S., Li, J., Zhang, X., Wei, H. & Cui, L. 2006. Effects of heat acclimation pretreatment on changes of membrane lipid peroxidation, antioxidant metabolites, and ultrastructure of chloroplasts in two cool-season turfgrass species under heat stress. Environmental and Experimental Botany, 56(3), 274–285.

122. Yan, J., Yu, L., Xuan, J., Lu, Y., Lu, S. & Zhu, W. 2016. De novo transcriptome sequencing and gene expression profiling of spinach (*Spinacia oleracea* L.) leaves under heat stress. Scientific Reports, 6(1), 19473.

123. Yang, Y., Kong, Q., Lim, A. R., Lu, S., Zhao, H., Guo, L., Yuan, L. & Ma, W. 2022. Transcriptional regulation of oil biosynthesis in seed plants: Current understanding, applications, and perspectives. Plant Communications, 3(5).

124. Yu, B., Yan, S., Zhou, H., Dong, R., Lei, J., Chen, C. & Cao, B. 2018. Overexpression of CsCaM3 improves high temperature tolerance in cucumber. Frontiers in Plant Science, 9(797.

125. Yue, L., Li, G., Dai, Y., Sun, X., Li, F., Zhang, S., Zhang, H., Sun, R. & Zhang, S. 2021. Gene co-expression network analysis of the heat-responsive core transcriptome identifies hub genes in *Brassica rapa*. Planta, 253(5), 111.

126. Zare, T., Fournier-Level, A., Ebert, B. & Roessner, U. 2024a. Chia (*Salvia hispanica* L.), a functional ‘superfood’: new insights into its botanical, genetic and nutraceutical characteristics. Annals of Botany, 134(5), 725–746.

127. Zare, T., Paril, J. F., Barnett, E. M., Kaur, P., Appels, R., Ebert, B., Roessner, U. & Fournier-Level, A. 2024b. Comparative genomics points to tandem duplications of SAD gene clusters as drivers of increased α-linolenic (ω-3) content in *S. hispanica* seeds. The Plant Genome, 17(1), e20430.

128. Zeng, C., Jia, T., Gu, T., Su, J. & Hu, X. 2021. Progress in research on the mechanisms underlying chloroplast-involved heat tolerance in plants. Genes, 12(9), 1343.

129. Zhang, W., Zhou, R.-G., Gao, Y.-J., Zheng, S.-Z., Xu, P., Zhang, S.-Q. & Sun, D.-Y. 2009. Molecular and genetic evidence for the key role of AtCaM3 in heat-shock signal transduction in Arabidopsis. Plant Physiology, 149(4), 1773–1784.

130. Zhang, Y., Li, Y., Han, B., Liu, A. & Xu, W. 2022. Integrated lipidomic and transcriptomic analysis reveals triacylglycerol accumulation in castor bean seedlings under heat stress. Industrial Crops and Products, 180(114702.

131. Zhao, J., Lu, Z., Wang, L. & Jin, B. 2020. Plant responses to heat stress: physiology, transcription, noncoding RNAs, and epigenetics. International Journal of Molecular Sciences, 22(1), 117.

132. Zhao, Y., Du, H., Wang, Y., Wang, H., Yang, S., Li, C., Chen, N., Yang, H., Zhang, Y. & Zhu, Y. 2021. The calcium-dependent protein kinase ZmCDPK7 functions in heat-stress tolerance in maize. Journal of Integrative Plant Biology, 63(3), 510–527.

133. Zhou, Y., Wang, Y., Xu, F., Song, C., Yang, X., Zhang, Z., Yi, M., Ma, N., Zhou, X. & He, J. 2022. Small HSPs play an important role in crosstalk between HSF-HSP and ROS pathways in heat stress response through transcriptomic analysis in lilies (*Lilium longiflorum*). BMC Plant Biology, 22(1), 202.

134. Zoong Lwe, Z. S., Welti, R., Anco, D., Naveed, S., Rustgi, S. & Narayanan, S. 2020. Heat stress elicits remodeling in the anther lipidome of peanut. Scientific Reports, 10(1), 22163.

