## Supporting Information for "Thermotolerance in Chia (*Salvia hispanica* L.) is Mediated by Rapid Heat-Induced Transcriptomic Reprogramming and Lipid Remodelling in Leaves"

**ORCIDs**

Tannaz Zare: https://orcid.org/0000-0002-7194-6800

Cheka Kehelpannala: https://orcid.org/0000-0001-7375-1161

Atul Bhatnagar: https://orcid.org/0000-0003-2153-299X

Thusitha W. Rupasinghe: https://orcid.org/0000-0001-7229-2469

Berit Ebert: https://orcid.org/0000-0002-6914-5473

Alexandre Fournier-Level: https://orcid.org/0000-0002-6047-7164

Ute Roessner: https://orcid.org/0000-0002-6482-2615

#### Supporting Figures

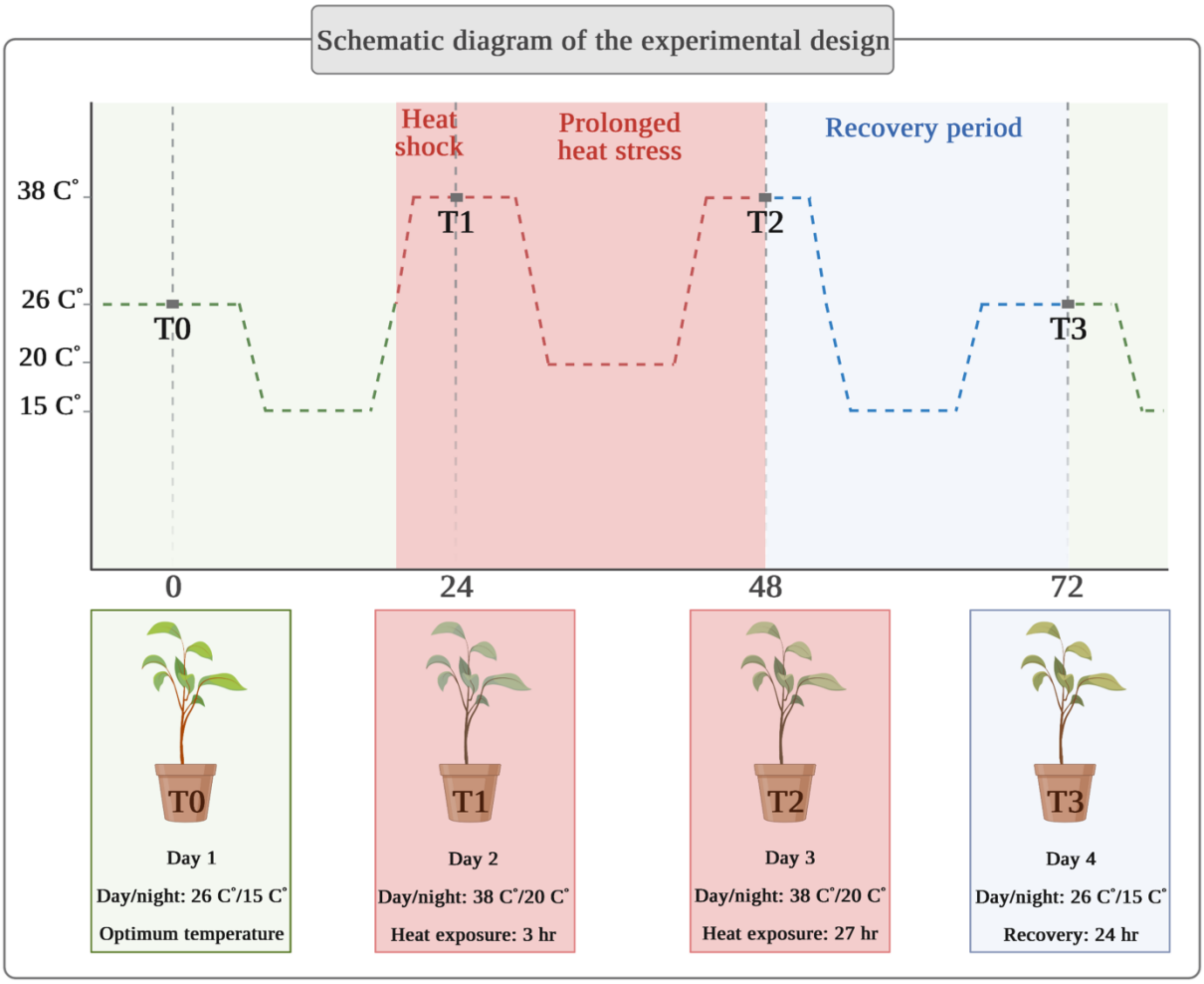


**Figure S1. Schematic overview of the experimental design to investigate heat stress responses in chia leaves.** Plants were grown under controlled conditions (green zone: 26°C/15°C day/night) until Day 1 (T0). Heat stress was applied on Day 2 (T1) with exposure to 38°C/20°C for 3 hr, followed by prolonged heat stress on Day 3 (T2) for 27 hr. Recovery conditions (blue zone), matching the control environment, were applied on Day 4 (T3) for 24 h. The red zone indicates the heat stress period, characterised by a relative humidity of 60–61%. The diagram was created using BioRender.com.

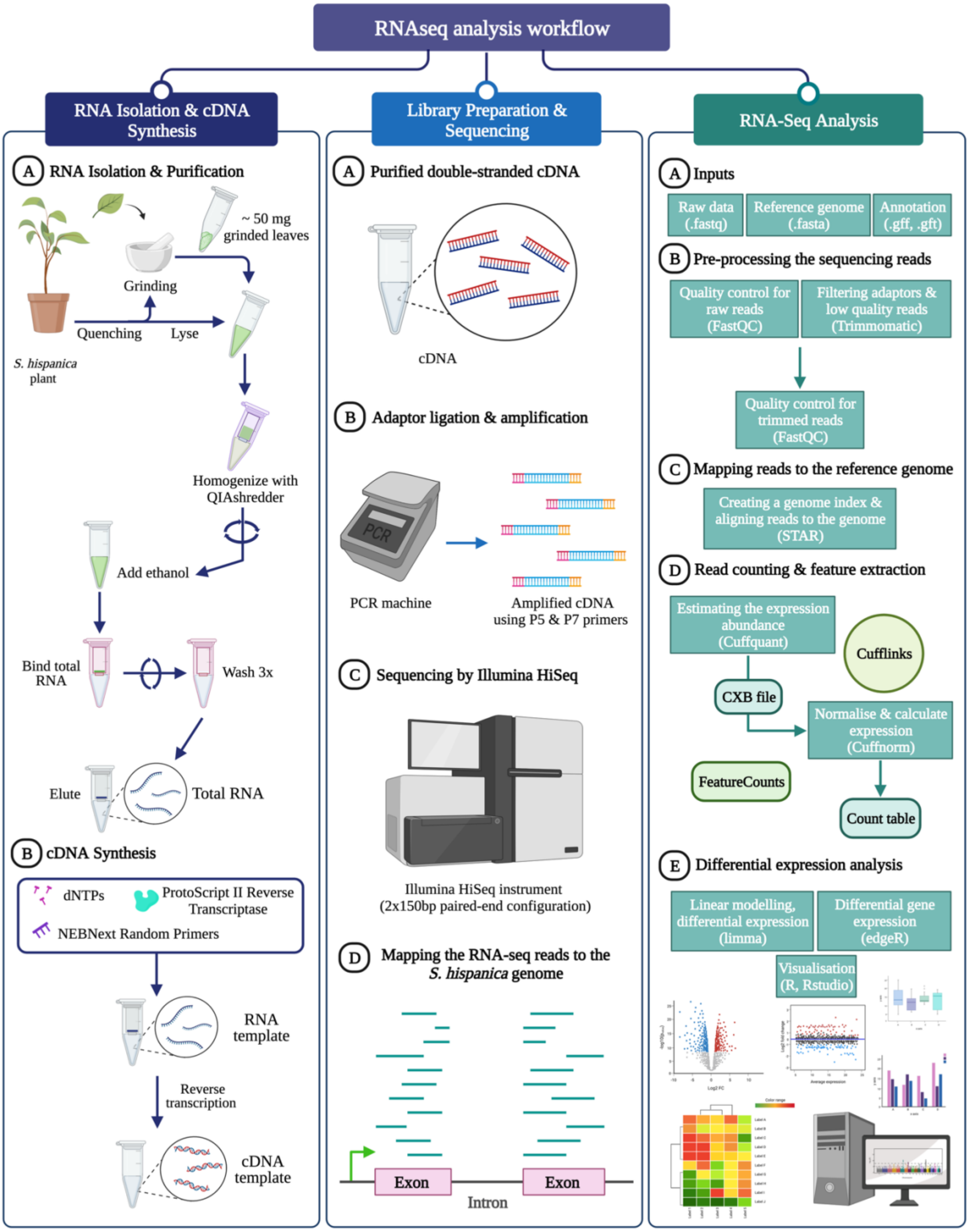


**Figure S2. Workflow for RNA isolation, RNA-seq library preparation, and transcriptomic analysis in chia leaves.** The workflow includes RNA extraction and cDNA synthesis followed by library preparation and sequencing on the Illumina HiSeq platform (2 × 150 bp). RNA-seq analysis involved quality control, expression quantification and differential expression analysis. The diagram was created using BioRender.com

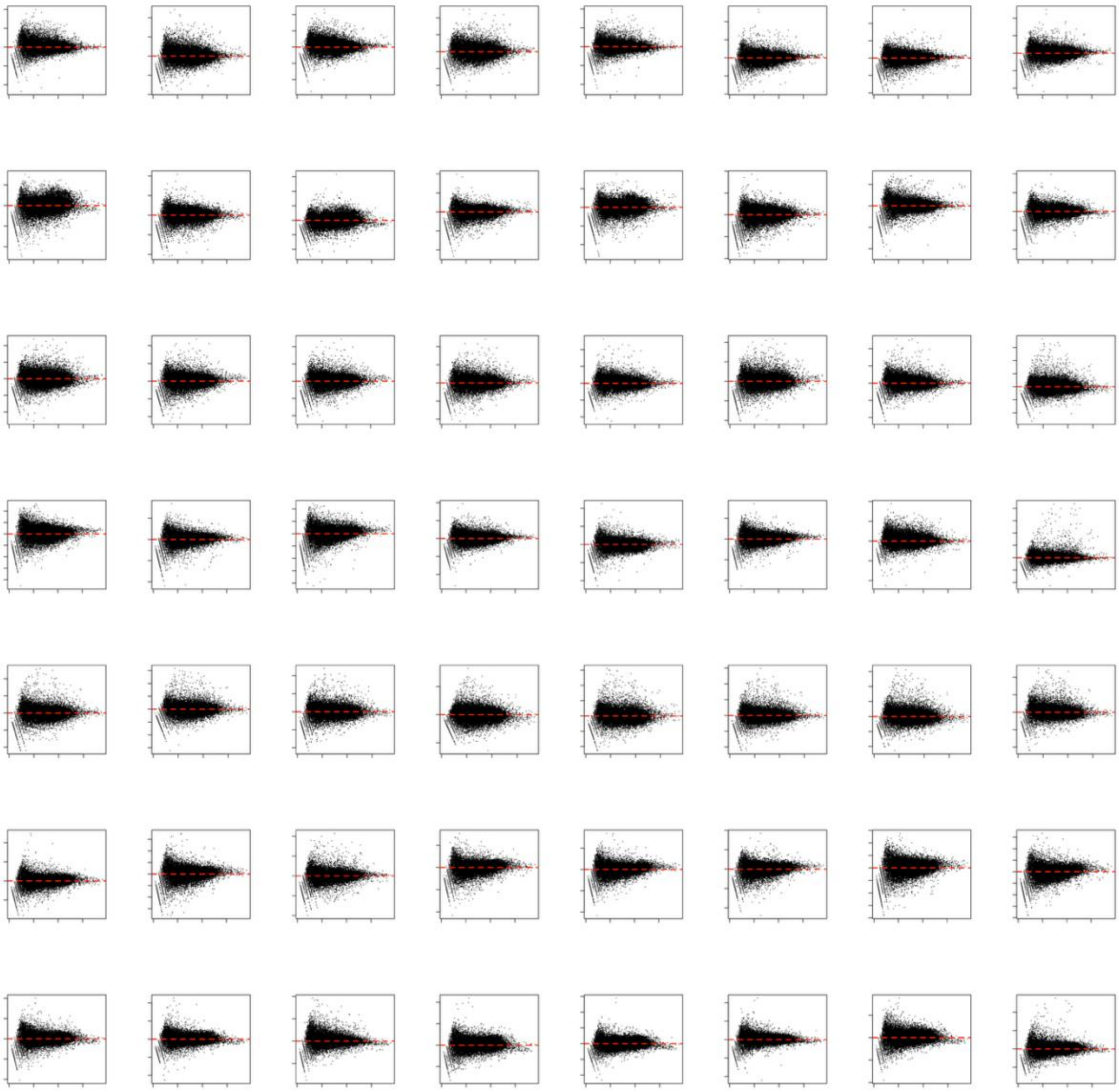

**Figure S3. Mean-difference (MD) plots for all RNA-seq samples.** MD plots were generated using the plotMD() function in the limma package to evaluate the effectiveness of Trimmed Mean of M-values (TMM) normalisation. Each plot compares the log-fold change of a sample (adjusted for library size) to the average log expression of a synthetic reference library composed of all other samples. Symmetry around the zero line indicates successful removal of composition bias across libraries.

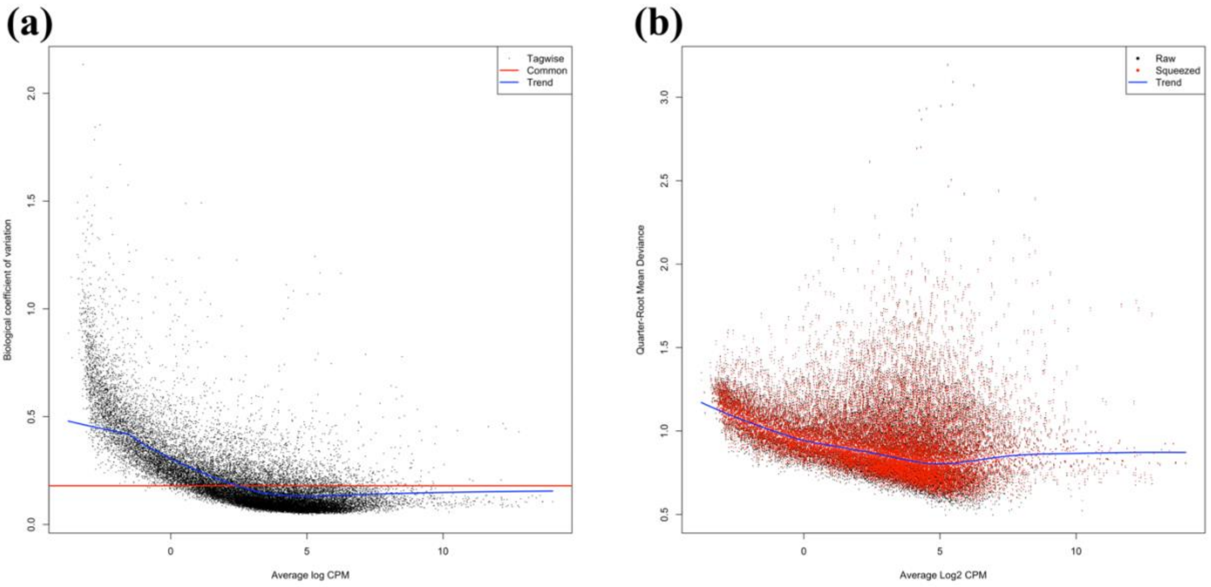


**Figure S4. Evaluation of dispersion estimates for differential expression modelling. (a)** Plot of genewise negative binomial (NB) dispersions, shown as the biological coefficient of variation (BCV), against average $\log_2$ counts per million (CPM). The common and trended dispersions are indicated by red and blue lines, respectively. **(b)** Plot of quarter-root quasi-likelihood (QL) dispersions against average $\log_2$ CPM. Black points represent raw genewise dispersions, while red points show "squeezed" dispersions after empirical Bayes shrinkage. The blue line indicates the trended QL dispersion.

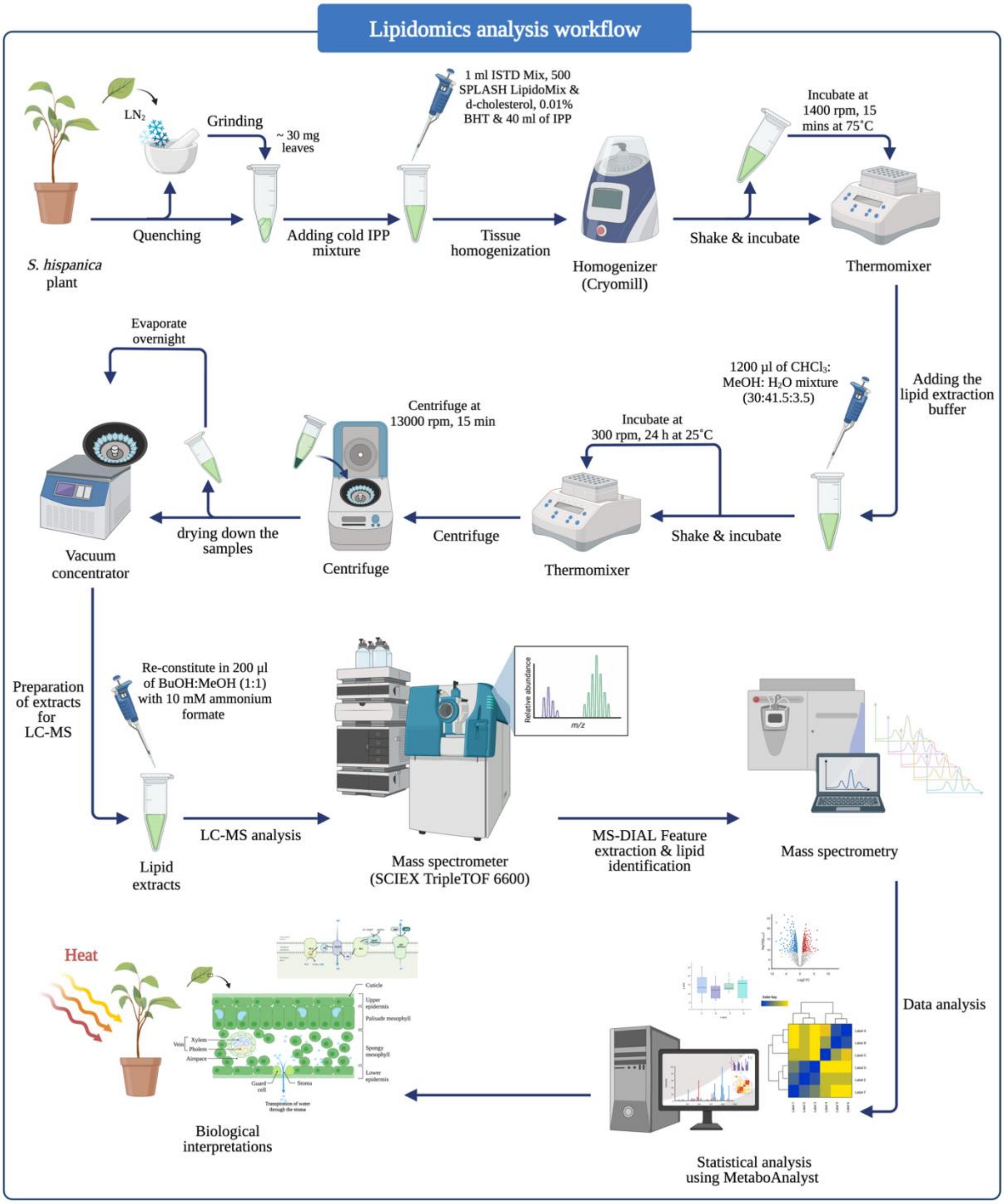


**Figure S5. Workflow of lipidomics analysis in Salvia hispanica leaves.** Lipid extraction was performed on ~30 mg of leaf tissue. Lipid extracts were analysed by LC-MS (SCIEX TripleTOF 6600), and MS-DIAL was used for feature extraction and identification. Statistical analysis was conducted in MetaboAnalyst, and results were interpreted in the context of heat stress responses. The diagram was created using BioRender.com.

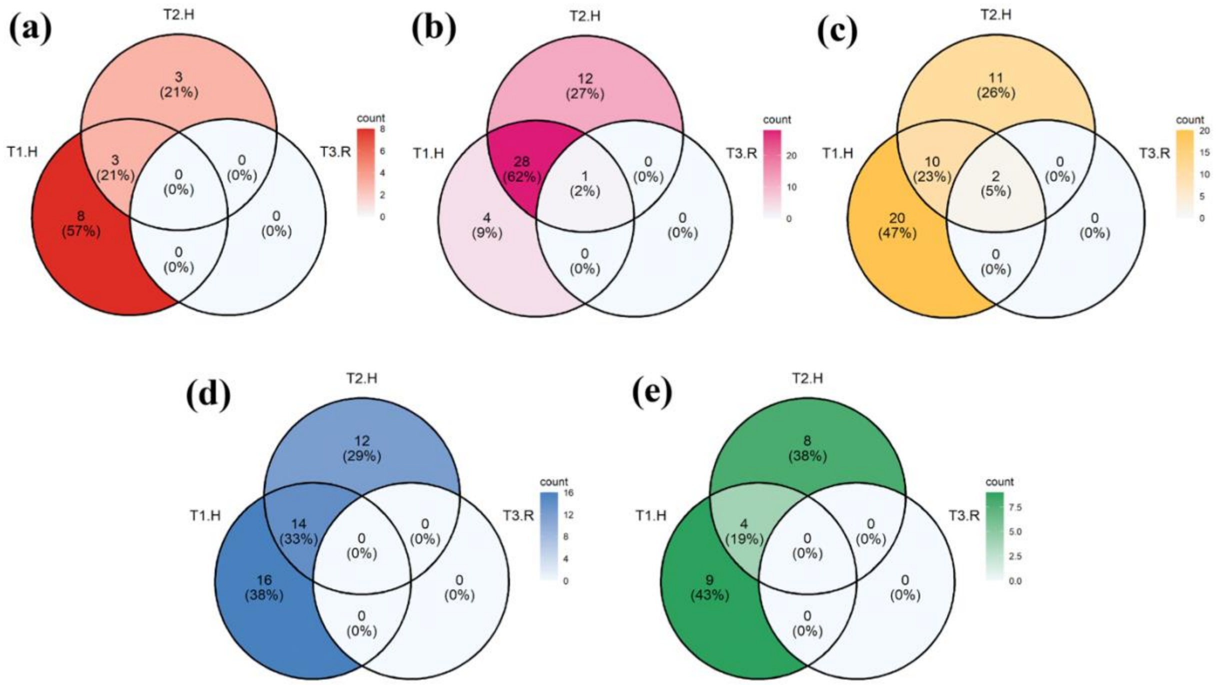


**Figure S6.Venn diagrams showing the number of stress-related genes identified across experimental conditions.** Each panel represents the distribution of genes detected under heat shock (T1.H), prolonged heat stress (T2.H), and recovery (T3.R) for specific functional categories: **(a)** heat stress transcription factors, (**b)** heat shock proteins, **(c)** chaperone proteins, **(d)** calcium ($Ca^{2+}$) transport and signalling-related genes, and **(e)** FK506-binding proteins (FKBPs). Percentages indicate the proportion of genes detected within each time point relative to the total number of genes in each category.

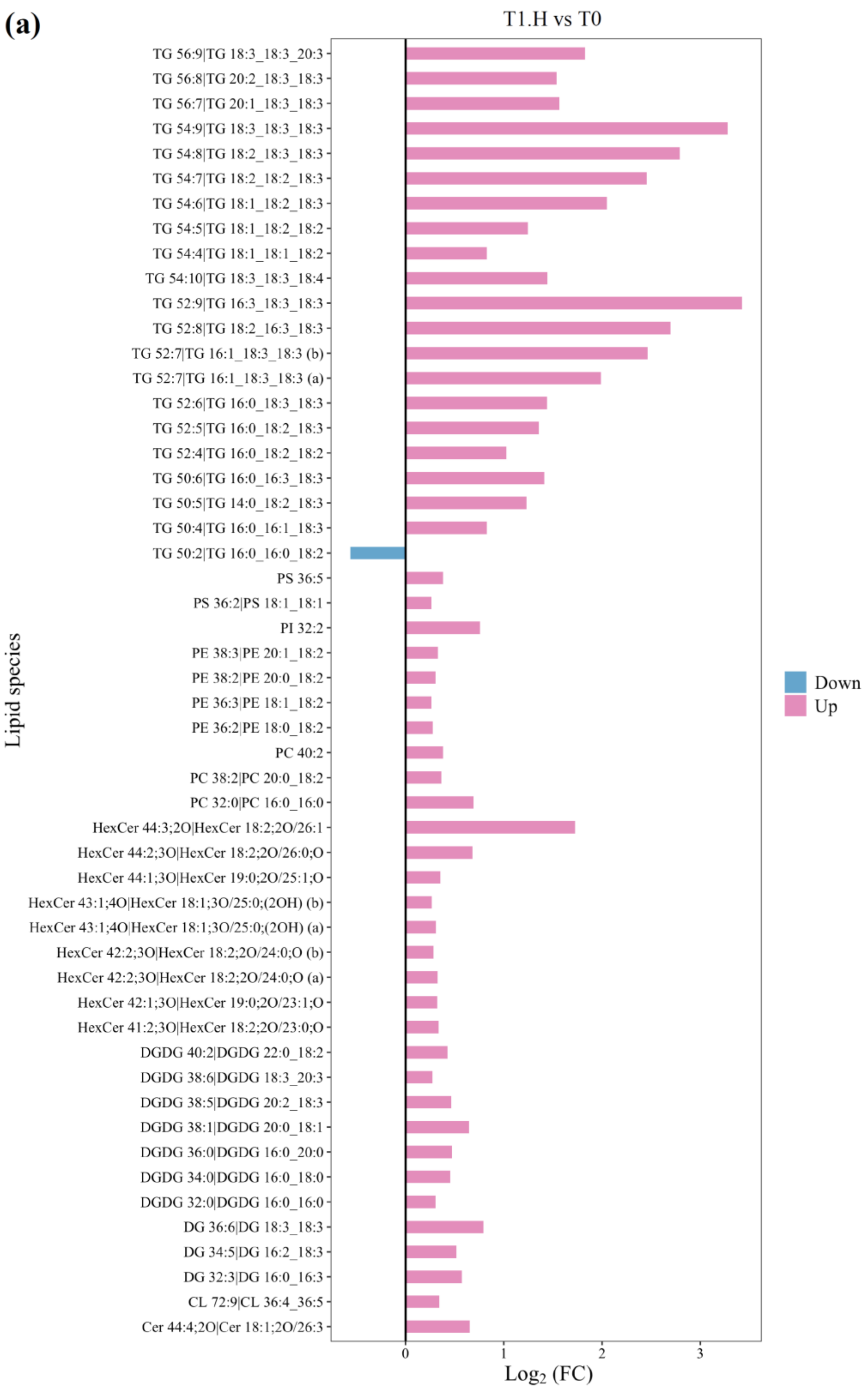
(a)
T1.H vs T0
TG 56:9|TG 18:3_18:3_20:3
TG 56:8|TG 20:2_18:3_18:3
TG 56:7|TG 20:1_18:3_18:3
TG 54:9|TG 18:3_18:3_18:3
TG 54:8|TG 18:2_18:3_18:3
TG 54:7|TG 18:2_18:2_18:3
TG 54:6|TG 18:1_18:2_18:3
TG 54:5|TG 18:1_18:2_18:2
TG 54:4|TG 18:1_18:1_18:2
TG 54:10|TG 18:3_18:3_18:4
TG 52:9|TG 16:3_18:3_18:3
TG 52:8|TG 18:2_16:3_18:3
TG 52:7|TG 16:1_18:3_18:3 (b)
TG 52:7|TG 16:1_18:3_18:3 (a)
TG 52:6|TG 16:0_18:3_18:3
TG 52:5|TG 16:0_18:2_18:3
TG 52:4|TG 16:0_18:2_18:2
TG 50:6|TG 16:0_16:3_18:3
TG 50:5|TG 14:0_18:2_18:3
TG 50:4|TG 16:0_16:1_18:3
TG 50:2|TG 16:0_16:0_18:2
PS 36:5
PS 36:2|PS 18:1_18:1
PI 32:2
PE 38:3|PE 20:1_18:2
PE 38:2|PE 20:0_18:2
PE 36:3|PE 18:1_18:2
PE 36:2|PE 18:0_18:2
PC 40:2
PC 38:2|PC 20:0_18:2
PC 32:0|PC 16:0_16:0
HexCer 44:3;2O|HexCer 18:2;2O/26:1
HexCer 44:2;3O|HexCer 18:2;2O/26:0;O
HexCer 44:1;3O|HexCer 19:0;2O/25:1;O
HexCer 43:1;4O|HexCer 18:1;3O/25:0;(2OH) (b)
HexCer 43:1;4O|HexCer 18:1;3O/25:0;(2OH) (a)
HexCer 42:2;3O|HexCer 18:2;2O/24:0;O (b)
HexCer 42:2;3O|HexCer 18:2;2O/24:0;O (a)
HexCer 42:1;3O|HexCer 19:0;2O/23:1;O
HexCer 41:2;3O|HexCer 18:2;2O/23:0;O
DGDG 40:2|DGDG 22:0_18:2
DGDG 38:6|DGDG 18:3_20:3
DGDG 38:5|DGDG 20:2_18:3
DGDG 38:1|DGDG 20:0_18:1
DGDG 36:0|DGDG 16:0_20:0
DGDG 34:0|DGDG 16:0_18:0
DGDG 32:0|DGDG 16:0_16:0
DG 36:6|DG 18:3_18:3
DG 34:5|DG 16:2_18:3
DG 32:3|DG 16:0_16:3
CL 72:9|CL 36:4_36:5
Cer 44:4;2O|Cer 18:1;2O/26:3
Lipid species
0
1
2
3
Log2 (FC)
Down
Up

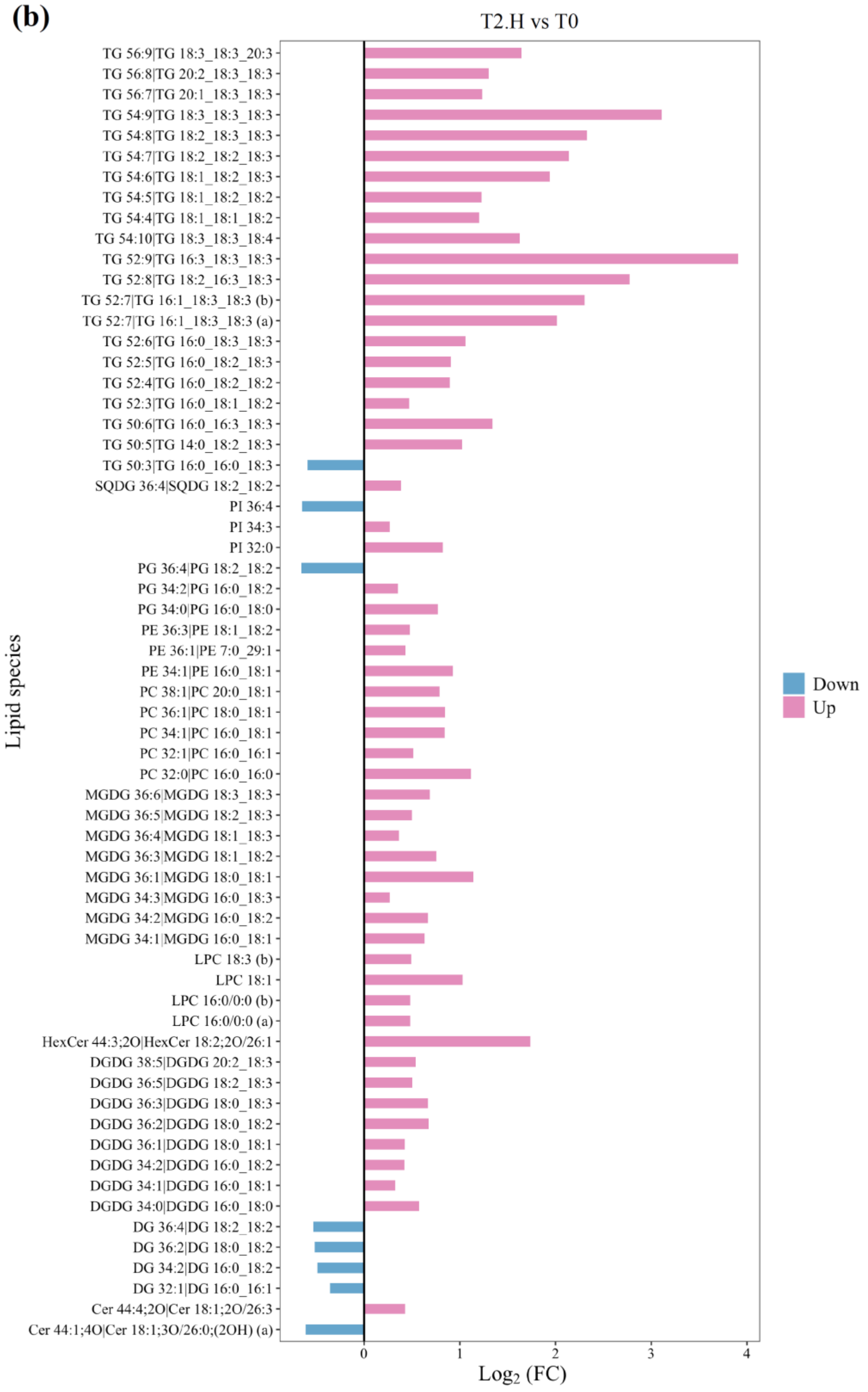


**Figure S7. Significantly altered lipid species in response to heat stress. (a)** $Log_2$ fold changes (FC) of individual lipid species following heat shock (T1.H). **(b)** $Log_2FC$ of lipid species following prolonged heat stress (T2.H). Significantly changed lipid species were identified using a false discovery rate (FDR) threshold of <0.05 and a fold change (FC) cutoff of ≥1.2.

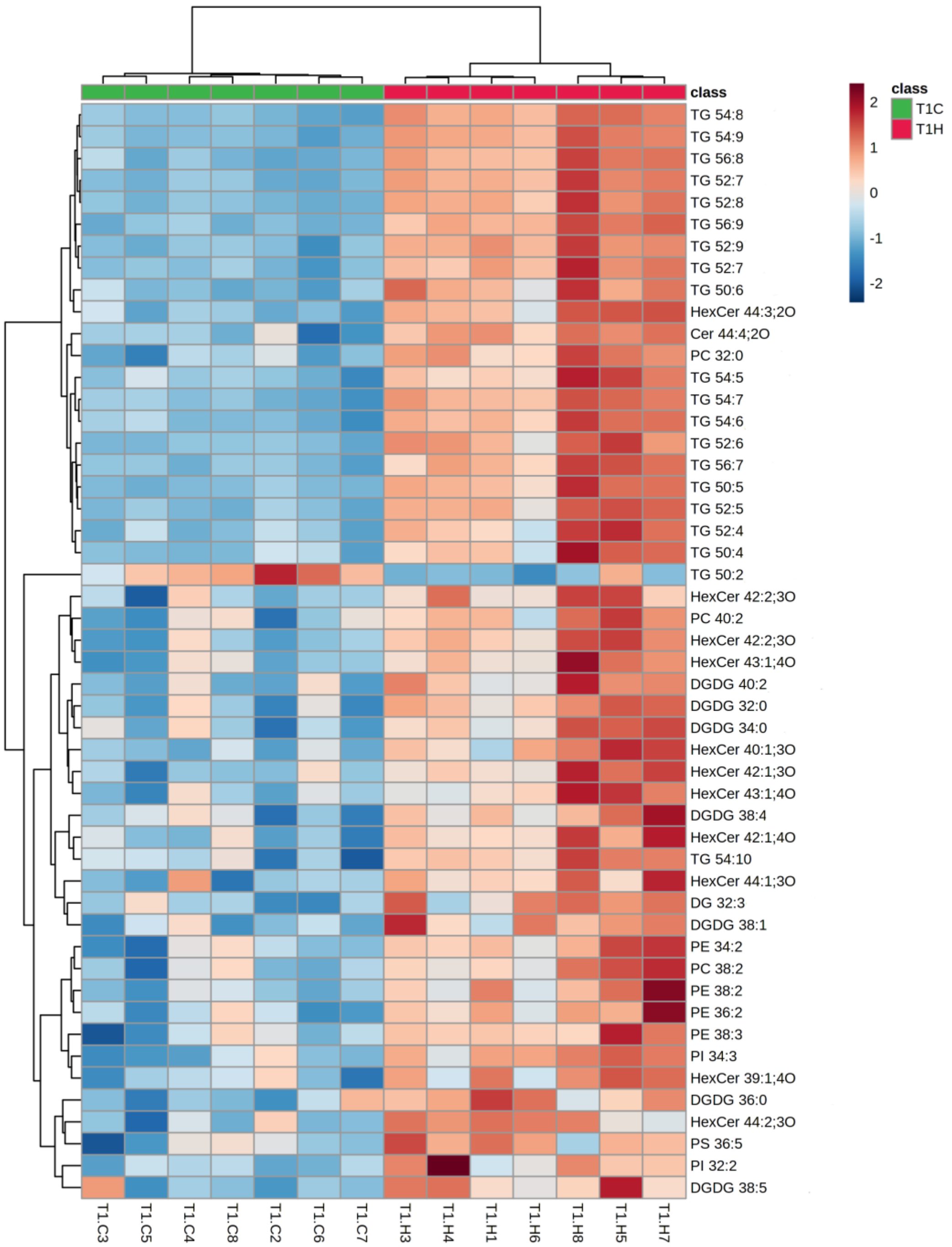


**Figure S8. Heatmap of lipid abundance in chia leaves under heat shock (T1.H) versus control (T1.C) conditions.** Each row represents a lipid species, and each column corresponds to a biological replicate. Heatmap colours reflect normalised peak area intensities. Hierarchical clustering was performed using the Ward algorithm with Pearson correlation as the distance metric. Lipid classes include triacylglycerols (TG), diacylglycerols (DG), digalactosyldiacylglycerols (DGDG), phosphatidylcholines (PC), phosphatidylethanolamines (PE), phosphatidylinositols (PI), phosphatidylserines (PS), ceramides (Cer), and hexosylceramides (HexCer).

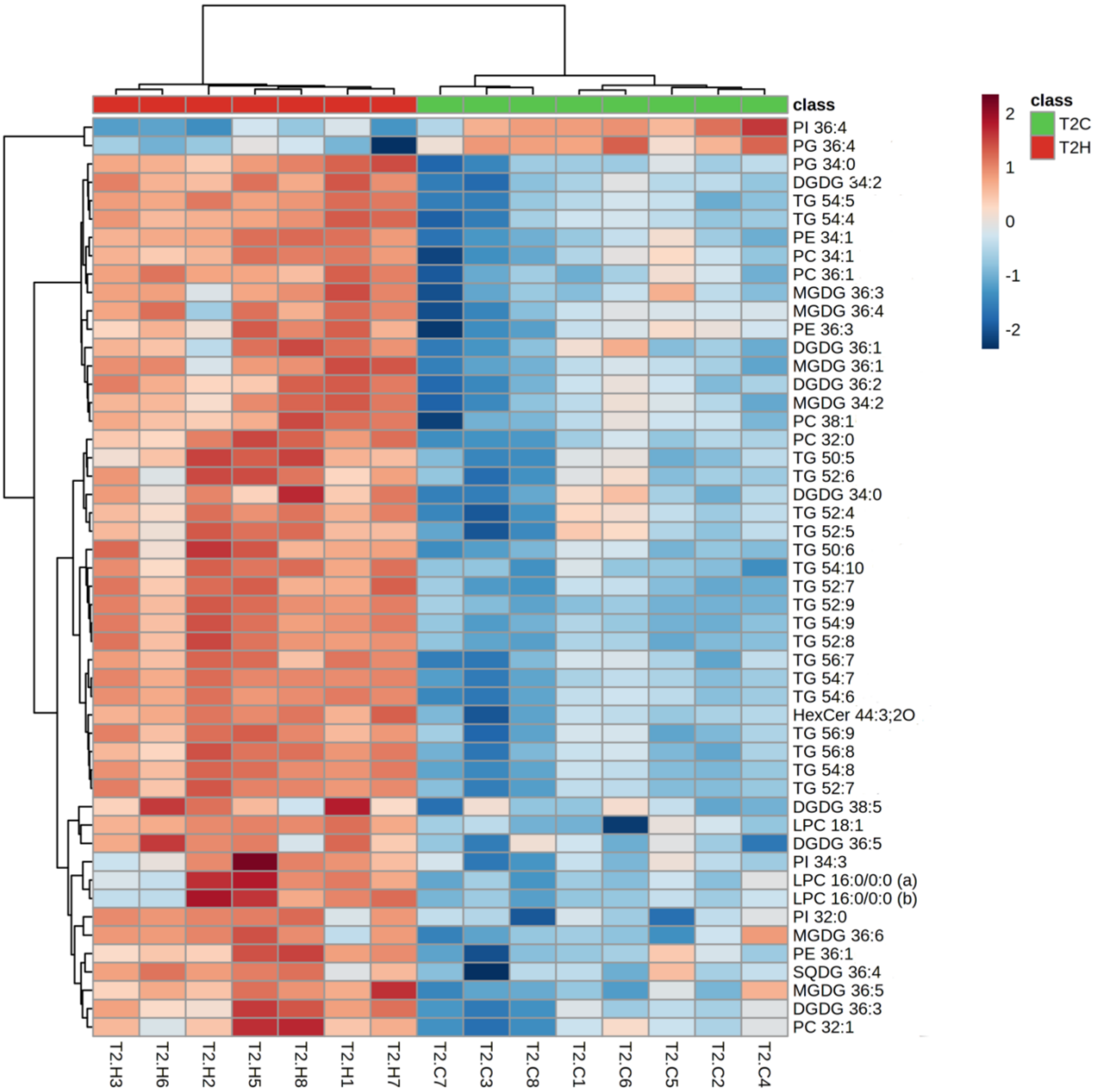


**Figure S9. Heatmap of lipid abundance in chia leaves under prolonged heat stress (T2.H) versus control (T2.C) conditions.** Each row represents a lipid species, and each column corresponds to a biological replicate. Heatmap colours indicate normalised peak area intensities. Hierarchical clustering was performed using the Ward algorithm with Pearson correlation as the distance metric. Lipid classes include triacylglycerols (TG), digalactosyldiacylglycerols (DGDG), monogalactosyldiacylglycerols (MGDG), sulfoquinovosyldiacylglycerols (SQDG), phosphatidylcholines (PC), phosphatidylethanolamines (PE), phosphatidylglycerols (PG), phosphatidylinositols (PI), lysophosphatidylcholines (LPC), and hexosylceramides (HexCer).

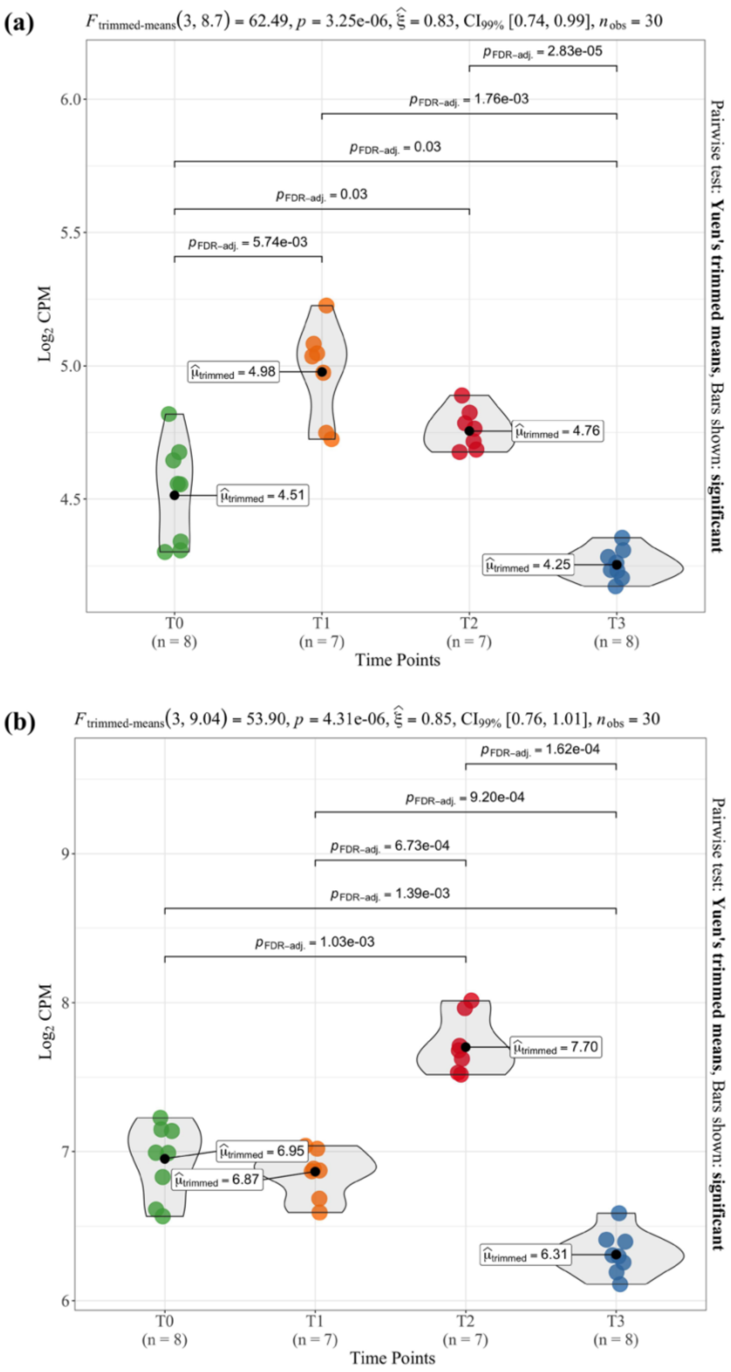


**Figure S10. Temporal Expression of Lipid Biosynthesis Genes in Chia under Heat Stress.** Violin plots show $\log_2$ counts per million (CPM) for **(a)** phospholipid diacylglycerol acyltransferase (*PDAT*) and **(b)** diacylglycerol acyltransferase (*DGAT*) genes across timepoints: T0 (control), T1 (heat shock), T2 (prolonged heat), and T3 (recovery). PDAT expression increased immediately following heat exposure, while DGAT showed delayed upregulation. Significant pairwise differences were determined using Yuen's trimmed means test with FDR-adjusted *p*-values ($\alpha = 0.05$).

### Supporting Tables

**Table S1.** Summary of sequencing quality metrics for raw RNA-seq reads across all libraries.

| Library ID | Raw reads | Raw base (G) | Error Rate (%) | Q20 (%) | Q30 (%) | GC Content (%) |
|---|---|---|---|---|---|---|
| T0-1 | 33761393 | 10.128 | 0.04 | 97.88 | 93.99 | 50.25 |
| T0-2 | 36424707 | 10.927 | 0.04 | 97.72 | 93.59 | 50.5 |
| T0-3 | 35976281 | 10.793 | 0.04 | 97.72 | 93.57 | 50.94 |
| T0-4 | 37737451 | 11.321 | 0.04 | 97.69 | 93.59 | 50.67 |
| T0-5 | 42899702 | 12.87 | 0.04 | 97.74 | 93.66 | 50.36 |
| T0-6 | 37969393 | 11.391 | 0.04 | 97.77 | 93.72 | 50.68 |
| T0-7 | 43745856 | 13.124 | 0.04 | 97.84 | 93.88 | 50.61 |
| T0-8 | 40098752 | 12.03 | 0.04 | 97.71 | 93.59 | 50.58 |
| T1-C1 | 34050338 | 10.215 | 0.04 | 97.77 | 93.74 | 50.16 |
| T1-C2 | 35114874 | 10.534 | 0.04 | 97.56 | 93.3 | 50.32 |
| T1-C3 | 39620313 | 11.886 | 0.04 | 97.76 | 93.65 | 49.83 |
| T1-C4 | 36208478 | 10.863 | 0.04 | 97.65 | 93.47 | 50.62 |
| T1-C5 | 32004846 | 9.601 | 0.04 | 97.69 | 93.48 | 50.41 |
| T1-C6 | 34910287 | 10.473 | 0.04 | 97.84 | 93.93 | 50.55 |
| T1-C7 | 33924751 | 10.177 | 0.04 | 97.75 | 93.67 | 50.72 |
| T1-C8 | 34068192 | 10.22 | 0.04 | 97.85 | 93.87 | 50.14 |
| T1-H1 | 38213422 | 11.464 | 0.04 | 97.73 | 93.62 | 50.33 |
| T1-H2 | 37961194 | 11.388 | 0.04 | 97.84 | 93.89 | 50.7 |
| T1-H3 | 34240458 | 10.272 | 0.04 | 97.91 | 94.03 | 50.83 |
| T1-H4 | 39294866 | 11.788 | 0.04 | 97.72 | 93.6 | 50.55 |
| T1-H5 | 40167361 | 12.05 | 0.04 | 97.66 | 93.47 | 50.46 |
| T1-H6 | 39268694 | 11.781 | 0.04 | 97.73 | 93.59 | 50.61 |
| T1-H7 | 31292293 | 9.388 | 0.04 | 97.84 | 93.84 | 50.45 |
| T1-H8 | 33020822 | 9.906 | 0.04 | 97.63 | 93.42 | 49.98 |
| T2-C1 | 36140998 | 10.842 | 0.04 | 97.87 | 93.94 | 50.86 |
| T2-C2 | 34909389 | 10.473 | 0.04 | 97.72 | 93.59 | 50.35 |
| T2-C3 | 39243723 | 11.773 | 0.04 | 97.68 | 93.5 | 50.8 |
| T2-C4 | 39423845 | 11.827 | 0.04 | 97.71 | 93.58 | 50.58 |
| T2-C5 | 36048962 | 10.815 | 0.04 | 97.79 | 93.78 | 50.45 |
| T2-C6 | 39290579 | 11.787 | 0.04 | 97.77 | 93.72 | 50.66 |
| T2-C7 | 47648364 | 14.295 | 0.04 | 97.87 | 93.93 | 50.34 |
| T2-C8 | 36751693 | 11.026 | 0.04 | 97.59 | 93.27 | 50.48 |
| T2-H1 | 33970556 | 10.191 | 0.04 | 97.77 | 93.72 | 50.28 |
| T2-H2 | 32040474 | 9.612 | 0.04 | 97.53 | 93.2 | 49.94 |
| T2-H3 | 36288948 | 10.887 | 0.04 | 97.75 | 93.62 | 50.08 |
| T2-H4 | 40653099 | 12.196 | 0.04 | 97.58 | 93.24 | 49.36 |
| T2-H5 | 35134069 | 10.54 | 0.04 | 97.64 | 93.32 | 50.29 |
| T2-H6 | 36519779 | 10.956 | 0.04 | 97.78 | 93.75 | 50 |
| T2-H7 | 38825919 | 11.648 | 0.04 | 97.79 | 93.72 | 49.98 |
| T2-H8 | 38795390 | 11.639 | 0.04 | 97.83 | 93.77 | 50.06 |
| T3-C1 | 40733214 | 12.22 | 0.04 | 97.72 | 93.58 | 50.41 |
| T3-C2 | 35496028 | 10.649 | 0.04 | 97.77 | 93.71 | 50.6 |
| T3-C3 | 37860211 | 11.358 | 0.04 | 97.81 | 93.79 | 50.3 |
| T3-C4 | 38559533 | 11.568 | 0.04 | 97.72 | 93.61 | 50.5 |
| T3-C5 | 41094427 | 12.328 | 0.04 | 97.67 | 93.52 | 50.66 |
| T3-C6 | 41424824 | 12.427 | 0.04 | 97.72 | 93.57 | 50.83 |
| T3-C7 | 32905388 | 9.872 | 0.04 | 97.81 | 93.78 | 50.75 |
| T3-C8 | 35726023 | 10.718 | 0.04 | 97.69 | 93.57 | 50.65 |
| T3-R1 | 35380066 | 10.614 | 0.04 | 97.84 | 93.88 | 50.2 |
| T3-R2 | 38493895 | 11.548 | 0.04 | 97.68 | 93.52 | 50.03 |
| T3-R3 | 37235826 | 11.171 | 0.04 | 97.66 | 93.44 | 50.05 |
| T3-R4 | 42145760 | 12.644 | 0.04 | 97.71 | 93.58 | 49.97 |
| T3-R5 | 46667840 | 14 | 0.04 | 97.78 | 93.7 | 49.81 |
| T3-R6 | 41147233 | 12.344 | 0.04 | 97.78 | 93.71 | 50.16 |
| T3-R7 | 46524069 | 13.957 | 0.04 | 97.88 | 93.94 | 49.99 |
| T3-R8 | 41059548 | 12.318 | 0.04 | 97.72 | 93.57 | 50.32 |

**Table S2.** Summary of RNA-seq read alignment statistics to the Salvia hispanica reference genome using STAR.

| Sample | Number of reads | Average read length | Uniquely mapped reads number | Average mapped length | Mismatch rate per base (%) | Deletion rate per base | Deletion average length | Reads mapped to multiple loci | % of reads unmapped |
|---|---|---|---|---|---|---|---|---|---|
| T0-1 | 26088668 | 275 | 21548815 | 274.42 | 0.22% | 0.01% | 2.66 | 1809500 | 10.46% |
| T0-2 | 29417773 | 275 | 24958853 | 274.52 | 0.22% | 0.01% | 2.62 | 2315256 | 7.28% |
| T0-3 | 29225266 | 275 | 24591742 | 274.56 | 0.22% | 0.01% | 2.71 | 2572917 | 7.04% |
| T0-4 | 30506784 | 275 | 26047047 | 274.6 | 0.22% | 0.01% | 2.66 | 2269121 | 7.17% |
| T0-5 | 35153219 | 275 | 30021346 | 274.59 | 0.22% | 0.01% | 2.65 | 2604382 | 7.18% |
| T0-6 | 31113115 | 275 | 26870396 | 274.63 | 0.21% | 0.01% | 2.68 | 2062736 | 7.00% |
| T0-7 | 35930706 | 275 | 31135526 | 274.65 | 0.21% | 0.01% | 2.67 | 2304245 | 6.93% |
| T0-8 | 32624316 | 275 | 28226449 | 274.64 | 0.22% | 0.01% | 2.7 | 2049004 | 7.19% |
| T1-C1 | 27767275 | 275 | 23984449 | 274.61 | 0.21% | 0.01% | 2.69 | 1759245 | 7.28% |
| T1-C2 | 28350801 | 275 | 24501088 | 274.58 | 0.22% | 0.01% | 2.68 | 1834992 | 7.10% |
| T1-C3 | 32502003 | 275 | 28104715 | 274.54 | 0.21% | 0.01% | 2.65 | 2064721 | 7.17% |
| T1-C4 | 29376919 | 275 | 24921360 | 274.58 | 0.22% | 0.01% | 2.62 | 2337488 | 7.20% |
| T1-C5 | 25977725 | 275 | 21867576 | 274.55 | 0.22% | 0.01% | 2.63 | 2214998 | 7.29% |
| T1-C6 | 28887122 | 275 | 25033652 | 274.63 | 0.21% | 0.01% | 2.67 | 1826325 | 7.01% |
| T1-C7 | 27731655 | 275 | 22972473 | 274.58 | 0.22% | 0.01% | 2.58 | 2800840 | 7.05% |
| T1-C8 | 28083084 | 275 | 24297237 | 274.60 | 0.21% | 0.01% | 2.68 | 1771225 | 7.17% |
| T1-H1 | 31066033 | 275 | 26513846 | 274.61 | 0.22% | 0.01% | 2.59 | 2259112 | 7.37% |
| T1-H2 | 30982474 | 275 | 26388573 | 274.63 | 0.21% | 0.01% | 2.64 | 2403477 | 7.06% |
| T1-H3 | 27996300 | 275 | 23773693 | 274.58 | 0.22% | 0.01% | 2.67 | 2203704 | 7.20% |
| T1-H4 | 31905646 | 275 | 27101800 | 274.59 | 0.22% | 0.01% | 2.59 | 2520965 | 7.15% |
| T1-H5 | 32701837 | 275 | 27921970 | 274.55 | 0.22% | 0.01% | 2.59 | 2458478 | 7.09% |
| T1-H6 | 31980838 | 275 | 27477919 | 274.60 | 0.22% | 0.01% | 2.62 | 2197273 | 7.20% |
| T1-H7 | 25628095 | 275 | 21894452 | 274.59 | 0.22% | 0.01% | 2.61 | 1903781 | 7.13% |
| T1-H8 | 26719848 | 275 | 22904080 | 274.56 | 0.21% | 0.01% | 2.64 | 1824361 | 7.45% |
| T2-C1 | 29548280 | 275 | 25447236 | 274.61 | 0.21% | 0.01% | 2.67 | 1993314 | 7.13% |
| T2-C2 | 28603455 | 275 | 24714479 | 274.64 | 0.21% | 0.01% | 2.69 | 1903876 | 6.93% |
| T2-C3 | 31856343 | 275 | 27443737 | 274.57 | 0.22% | 0.01% | 2.66 | 2198565 | 6.94% |
| T2-C4 | 32131120 | 275 | 27570924 | 274.60 | 0.22% | 0.01% | 2.67 | 2236840 | 7.22% |
| T2-C5 | 29528096 | 275 | 25252023 | 274.61 | 0.22% | 0.01% | 2.66 | 2072533 | 7.45% |
| T2-C6 | 32327354 | 275 | 27879321 | 274.77 | 0.21% | 0.01% | 2.65 | 2277656 | 6.71% |
| T2-C7 | 38954778 | 275 | 33507745 | 274.60 | 0.21% | 0.01% | 2.66 | 2509255 | 7.53% |
| T2-C8 | 29678944 | 275 | 25247821 | 274.61 | 0.22% | 0.01% | 2.59 | 2302769 | 7.16% |
| T2-H1 | 27532144 | 275 | 23453024 | 274.59 | 0.21% | 0.01% | 2.59 | 1960925 | 7.69% |
| T2-H2 | 25514110 | 275 | 21717498 | 274.56 | 0.22% | 0.01% | 2.59 | 1837761 | 7.67% |
| T2-H3 | 29593903 | 275 | 25423894 | 274.59 | 0.22% | 0.01% | 2.56 | 1983512 | 7.38% |
| T2-H4 | 32908077 | 275 | 27784575 | 274.67 | 0.22% | 0.01% | 2.58 | 2573854 | 7.74% |
| T2-H5 | 28336727 | 275 | 23613972 | 274.59 | 0.22% | 0.01% | 2.59 | 2601633 | 7.48% |
| T2-H6 | 29915376 | 275 | 25593991 | 274.58 | 0.21% | 0.01% | 2.60 | 2094501 | 7.44% |
| T2-H7 | 31754671 | 275 | 27038308 | 274.62 | 0.21% | 0.01% | 2.59 | 2281247 | 7.66% |
| T2-H8 | 31726990 | 275 | 27132917 | 274.59 | 0.21% | 0.01% | 2.58 | 2201314 | 7.53% |
| T3-C1 | 33071964 | 275 | 28299599 | 274.59 | 0.22% | 0.01% | 2.58 | 2335700 | 7.36% |
| T3-C2 | 29040984 | 275 | 24881791 | 274.59 | 0.21% | 0.01% | 2.60 | 2108862 | 7.05% |
| T3-C3 | 30768141 | 275 | 26570807 | 274.63 | 0.21% | 0.01% | 2.63 | 2024577 | 7.05% |
| T3-C4 | 31464014 | 275 | 27133806 | 274.58 | 0.22% | 0.01% | 2.62 | 1976004 | 7.47% |
| T3-C5 | 33487484 | 275 | 28703547 | 274.61 | 0.22% | 0.01% | 2.63 | 2373932 | 7.19% |
| T3-C6 | 33766221 | 275 | 29113325 | 274.62 | 0.22% | 0.01% | 2.60 | 2247027 | 7.12% |
| T3-C7 | 27047994 | 275 | 23337126 | 274.63 | 0.21% | 0.01% | 2.62 | 1809079 | 7.02% |
| T3-C8 | 29172470 | 275 | 25238511 | 274.64 | 0.21% | 0.01% | 2.62 | 1856237 | 7.11% |
| T3-R1 | 29090283 | 275 | 25079409 | 274.58 | 0.21% | 0.01% | 2.61 | 1882249 | 7.31% |
| T3-R2 | 31397769 | 275 | 27139376 | 274.60 | 0.21% | 0.01% | 2.60 | 2010016 | 7.15% |
| T3-R3 | 30333219 | 275 | 26264255 | 274.57 | 0.22% | 0.01% | 2.62 | 1902275 | 7.14% |
| T3-R4 | 34458378 | 275 | 29798775 | 274.62 | 0.21% | 0.01% | 2.64 | 2204020 | 7.12% |
| T3-R5 | 38331564 | 275 | 33000993 | 274.59 | 0.21% | 0.01% | 2.60 | 2528378 | 7.30% |
| T3-R6 | 33771872 | 275 | 28880557 | 274.57 | 0.21% | 0.01% | 2.64 | 2468213 | 7.17% |
| T3-R7 | 38456269 | 275 | 33163683 | 274.62 | 0.21% | 0.01% | 2.64 | 2530156 | 7.18% |
| T3-R8 | 33435391 | 275 | 28817499 | 274.62 | 0.21% | 0.01% | 2.63 | 2168810 | 7.32% |

**Table S3.** Literature-derived lipid metabolism genes identified in chia.

| Description (based on chia's annotated RefSeq) | Abbreviation | Pathway | Locus ID |
|---|---|---|---|
| pyruvate dehydrogenase E1 component subunit alpha-3, chloroplastic-like | *ShPDH E1 alpha* | Plastidial Fatty Acid Synthesis | LOC125188973 |
| pyruvate dehydrogenase E1 component subunit alpha-3, chloroplastic-like | *ShPDH E1 alpha* | Plastidial Fatty Acid Synthesis | LOC125197323 |
| pyruvate dehydrogenase E1 component subunit beta-3, chloroplastic-like | *ShPDH E1 beta* | Plastidial Fatty Acid Synthesis | LOC125196267 |
| pyruvate dehydrogenase E1 component subunit beta-3, chloroplastic-like | *ShPDH E1 beta* | Plastidial Fatty Acid Synthesis | LOC125202643 |
| pyruvate dehydrogenase E1 component subunit beta-3, chloroplastic-like | *ShPDH E1 beta* | Plastidial Fatty Acid Synthesis | LOC125208933 |
| acetyl-CoA carboxylase carboxyltransferase alpha subunit, chloroplastic | *ShACC α CT/CAC3* | Plastidial Fatty Acid Synthesis | LOC125222552 |
| acetyl-CoA carboxylase carboxyltransferase beta subunit, chloroplastic | *ShACC β-CT/ACCD* | Plastidial Fatty Acid Synthesis | LOC125213838 |
| acetyl-CoA carboxylase 1-like | *ShACC* | Plastidial Fatty Acid Synthesis | LOC125223679 |
| biotin carboxyl carrier protein of acetyl-CoA carboxylase | *ShACC BC/CAC2* | Plastidial Fatty Acid Synthesis | LOC125210332 |
| biotin carboxylase 1, chloroplastic-like | *ShACC BC1* | Plastidial Fatty Acid Synthesis | LOC125213153 |
| biotin carboxylase 2, chloroplastic-like | *ShACC BC2* | Plastidial Fatty Acid Synthesis | LOC125210767 |
| biotin carboxyl carrier protein of acetyl-CoA carboxylase 1, chloroplastic | *ShACC BCCP1/CAC1A* | Plastidial Fatty Acid Synthesis | LOC125210279 |
| biotin carboxyl carrier protein of acetyl-CoA carboxylase 2, chloroplastic | *ShACC BCCP2/CAC1B* | Plastidial Fatty Acid Synthesis | LOC125189241 |
| biotin carboxyl carrier protein of acetyl-CoA carboxylase 2 isoform X1 | *ShACC BCCP2/CAC1B* | Plastidial Fatty Acid Synthesis | LOC125213190 |
| biotin carboxyl carrier protein of acetyl-CoA carboxylase 2 isoform X2 | *ShACC BCCP2/CAC1B* | Plastidial Fatty Acid Synthesis | LOC125213190 |
| Malonyl-CoA : ACP Malonyltransferase | *ShMCMT* | Plastidial Fatty Acid Synthesis | LOC125211095 |
| 3-oxoacyl-[acyl-carrier-protein] synthase III, chloroplastic | *ShKAS III* | Plastidial Fatty Acid Synthesis | LOC125186086 |
| 3-oxoacyl-[acyl-carrier-protein] synthase III, chloroplastic | *ShKAS III* | Plastidial Fatty Acid Synthesis | LOC125199315 |
| 3-oxoacyl-[acyl-carrier-protein] synthase III, chloroplastic | *ShKAS III* | Plastidial Fatty Acid Synthesis | LOC125199728 |
| 3-oxoacyl-[acyl-carrier-protein] reductase FabG-like | *ShKAR/FabG* | Plastidial Fatty Acid Synthesis | LOC125185153 |
| 3-oxoacyl-[acyl-carrier-protein] reductase FabG-like | *ShKAR/FabG* | Plastidial Fatty Acid Synthesis | LOC125197366 |
| 3-oxoacyl-[acyl-carrier-protein] reductase FabG-like | *ShKAR/FabG* | Plastidial Fatty Acid Synthesis | LOC125187197 |
| 3-oxoacyl-[acyl-carrier-protein] reductase FabG-like | *ShKAR/FabG* | Plastidial Fatty Acid Synthesis | LOC125187634 |
| 3-oxoacyl-[acyl-carrier-protein] reductase FabG-like | *ShKAR/FabG* | Plastidial Fatty Acid Synthesis | LOC125187767 |
| 3-oxoacyl-[acyl-carrier-protein] reductase FabG-like | *ShKAR/FabG* | Plastidial Fatty Acid Synthesis | LOC125190879 |

| | | | |
|---|---|---|---|
| 3-oxoacyl-[acyl-carrier-protein] reductase FabG-like | *ShKAR/FabG* | Plastidial Fatty Acid Synthesis | LOC125191177 |
| 3-oxoacyl-[acyl-carrier-protein] reductase FabG-like | *ShKAR/FabG* | Plastidial Fatty Acid Synthesis | LOC125192402 |
| 3-hydroxyacyl-[acyl-carrier-protein] dehydratase FabZ-like isoform X1 | *ShHAD/FabZ* | Plastidial Fatty Acid Synthesis | LOC125222702 |
| 3-hydroxyacyl-[acyl-carrier-protein] dehydratase FabZ-like isoform X2 | *ShHAD/FabZ* | Plastidial Fatty Acid Synthesis | LOC125222702 |
| 3-hydroxyacyl-[acyl-carrier-protein] dehydratase FabZ-like | *ShHAD/FabZ* | Plastidial Fatty Acid Synthesis | LOC125211668 |
| 3-hydroxyacyl-[acyl-carrier-protein] dehydratase FabZ-like | *ShHAD/FabZ* | Plastidial Fatty Acid Synthesis | LOC125219044 |
| enoyl-[acyl-carrier-protein] reductase | *ShENR* | Plastidial Fatty Acid Synthesis | LOC125220956 |
| enoyl-[acyl-carrier-protein] reductase | *ShENR* | Plastidial Fatty Acid Synthesis | LOC125210931 |
| enoyl-[acyl-carrier-protein] reductase | *ShENR* | Plastidial Fatty Acid Synthesis | LOC125187061 |
| 3-oxoacyl-[acyl-carrier-protein] synthase I, chloroplastic-like isoform X1 | *ShKAS I* | Plastidial Fatty Acid Synthesis | LOC125215740 |
| 3-oxoacyl-[acyl-carrier-protein] synthase I, chloroplastic-like isoform X2 | *ShKAS I* | Plastidial Fatty Acid Synthesis | LOC125215740 |
| 3-oxoacyl-[acyl-carrier-protein] synthase I, chloroplastic-like isoform X3 | *ShKAS I* | Plastidial Fatty Acid Synthesis | LOC125215740 |
| 3-oxoacyl-[acyl-carrier-protein] synthase II, chloroplastic-like isoform X1 | *ShKAS II/FAB1* | Elongation, Desaturation and Export | LOC125216292 |
| 3-oxoacyl-[acyl-carrier-protein] synthase II, chloroplastic-like isoform X2 | *ShKAS II/FAB1* | Elongation, Desaturation and Export | LOC125215511 |
| 3-oxoacyl-[acyl-carrier-protein] synthase II, chloroplastic-like isoform X3 | *ShKAS II/FAB1* | Elongation, Desaturation and Export | LOC125215511 |
| 3-oxoacyl-[acyl-carrier-protein] synthase II, chloroplastic-like | *ShKAS II/FAB1* | Elongation, Desaturation and Export | LOC125188702 |
| stearoyl-[acyl-carrier-protein] 9-desaturase, chloroplastic-like isoform X1 | *ShSAD/FAB2* | Elongation, Desaturation and Export | LOC125214070 |
| stearoyl-[acyl-carrier-protein] 9-desaturase, chloroplastic-like isoform X2 | *ShSAD/FAB2* | Elongation, Desaturation and Export | LOC125214070 |
| stearoyl-[acyl-carrier-protein] 9-desaturase, chloroplastic-like isoform X1 | *ShSAD/FAB2* | Elongation, Desaturation and Export | LOC125201215 |
| stearoyl-[acyl-carrier-protein] 9-desaturase, chloroplastic-like isoform X2 | *ShSAD/FAB2* | Elongation, Desaturation and Export | LOC125201215 |
| stearoyl-[acyl-carrier-protein] 9-desaturase, chloroplastic-like | *ShSAD/FAB2* | Elongation, Desaturation and Export | LOC125214093 |
| stearoyl-[acyl-carrier-protein] 9-desaturase, chloroplastic-like | *ShSAD/FAB2* | Elongation, Desaturation and Export | LOC125214082 |
| stearoyl-[acyl-carrier-protein] 9-desaturase, chloroplastic-like | *ShSAD/FAB2* | Elongation, Desaturation and Export | LOC125212987 |
| stearoyl-[acyl-carrier-protein] 9-desaturase, chloroplastic-like | *ShSAD/FAB2* | Elongation, Desaturation and Export | LOC125211105 |
| stearoyl-[acyl-carrier-protein] 9-desaturase, chloroplastic-like | *ShSAD/FAB2* | Elongation, Desaturation and Export | LOC125201496 |
| stearoyl-[acyl-carrier-protein] 9-desaturase, chloroplastic-like | *ShSAD/FAB2* | Elongation, Desaturation and Export | LOC125201429 |
| stearoyl-[acyl-carrier-protein] 9-desaturase, chloroplastic-like | *ShSAD/FAB2* | Elongation, Desaturation and Export | LOC125200966 |
| stearoyl-[acyl-carrier-protein] 9-desaturase, chloroplastic-like | *ShSAD/FAB2* | Elongation, Desaturation and Export | LOC125186413 |

| | | | |
|---|---|---|---|
| stearoyl-[acyl-carrier-protein] 9-desaturase, chloroplastic-like | *ShSAD/FAB2* | Elongation, Desaturation and Export | LOC125214680 |
| stearoyl-[acyl-carrier-protein] 9-desaturase, chloroplastic-like | *ShSAD/FAB2* | Elongation, Desaturation and Export | LOC125223686 |
| palmitoyl-acyl carrier protein thioesterase, chloroplastic-like | *ShFATA/B* | Elongation, Desaturation and Export | LOC125191012 |
| palmitoyl-acyl carrier protein thioesterase, chloroplastic-like | *ShFATA/B* | Elongation, Desaturation and Export | LOC125206644 |
| palmitoyl-acyl carrier protein thioesterase, chloroplastic-like | *ShFATA/B* | Elongation, Desaturation and Export | LOC125210040 |
| palmitoyl-acyl carrier protein thioesterase, chloroplastic-like | *ShFATA/B* | Elongation, Desaturation and Export | LOC125215586 |
| palmitoyl-acyl carrier protein thioesterase, chloroplastic-like | *ShFATA/B* | Elongation, Desaturation and Export | LOC125188909 |
| palmitoyl-acyl carrier protein thioesterase, chloroplastic-like | *ShFATA/B* | Elongation, Desaturation and Export | LOC125197186 |
| long chain acyl-CoA synthetase 1 | *ShLACS1* | Elongation, Desaturation and Export | LOC125216324 |
| long chain acyl-CoA synthetase 8 isoform X1 | *ShLACS8* | Elongation, Desaturation and Export | LOC125214579 |
| long chain acyl-CoA synthetase 8 isoform X2 | *ShLACS8* | Elongation, Desaturation and Export | LOC125214579 |
| long chain acyl-CoA synthetase 9 | *ShLACS9* | Elongation, Desaturation and Export | LOC125223859 |
| long chain acyl-CoA synthetase 4-like | *ShLACS4* | Elongation, Desaturation and Export | LOC125196331 |
| long chain acyl-CoA synthetase 4-like | *ShLACS4* | Elongation, Desaturation and Export | LOC125200993 |
| long chain acyl-CoA synthetase 4-like | *ShLACS4* | Elongation, Desaturation and Export | LOC125211764 |
| long chain acyl-CoA synthetase 4-like | *ShLACS4* | Elongation, Desaturation and Export | LOC125213009 |
| 3-oxoacyl-[acyl-carrier-protein] synthase II, chloroplastic-like isoform X1 | *ShKAS II/FAB1* | Elongation, Desaturation and Export | LOC125216292 |
| 3-oxoacyl-[acyl-carrier-protein] synthase II, chloroplastic-like isoform X2 | *ShKAS II/FAB1* | Elongation, Desaturation and Export | LOC125215511 |
| 3-oxoacyl-[acyl-carrier-protein] synthase II, chloroplastic-like isoform X3 | *ShKAS II/FAB1* | Elongation, Desaturation and Export | LOC125215511 |
| 3-oxoacyl-[acyl-carrier-protein] synthase II, chloroplastic-like | *ShKAS II/FAB1* | Elongation, Desaturation and Export | LOC125188702 |
| stearoyl-[acyl-carrier-protein] 9-desaturase, chloroplastic-like isoform X1 | *ShSAD/FAB2* | Elongation, Desaturation and Export | LOC125214070 |
| stearoyl-[acyl-carrier-protein] 9-desaturase, chloroplastic-like isoform X2 | *ShSAD/FAB2* | Elongation, Desaturation and Export | LOC125214070 |
| stearoyl-[acyl-carrier-protein] 9-desaturase, chloroplastic-like isoform X1 | *ShSAD/FAB2* | Elongation, Desaturation and Export | LOC125201215 |
| stearoyl-[acyl-carrier-protein] 9-desaturase, chloroplastic-like isoform X2 | *ShSAD/FAB2* | Elongation, Desaturation and Export | LOC125201215 |
| stearoyl-[acyl-carrier-protein] 9-desaturase, chloroplastic-like | *ShSAD/FAB2* | Elongation, Desaturation and Export | LOC125214093 |
| stearoyl-[acyl-carrier-protein] 9-desaturase, chloroplastic-like | *ShSAD/FAB2* | Elongation, Desaturation and Export | LOC125214082 |
| stearoyl-[acyl-carrier-protein] 9-desaturase, chloroplastic-like | *ShSAD/FAB2* | Elongation, Desaturation and Export | LOC125212987 |
| glycerol-3-phosphate dehydrogenase [NAD(+)] isoform X1 | *ShGPD* | Glycerolipid/Galactolipid Synthesis | LOC125196206 |

| | | | |
|---|---|---|---|
| glycerol-3-phosphate dehydrogenase [NAD(+)] isoform X2 | *ShGPD* | Glycerolipid/Galactolipid Synthesis | LOC125196206 |
| glycerol-3-phosphate dehydrogenase [NAD(+)] 2, chloroplastic | *ShGPD2* | Glycerolipid/Galactolipid Synthesis | LOC125186718 |
| glycerol-3-phosphate acyltransferase ATS11, chloroplastic-like isoform X1 | *ShGPAT/ATS11* | Glycerolipid/Galactolipid Synthesis | LOC125221688 |
| glycerol-3-phosphate acyltransferase ATS11, chloroplastic-like isoform X2 | *ShGPAT/ATS11* | Glycerolipid/Galactolipid Synthesis | LOC125221688 |
| glycerol-3-phosphate acyltransferase ATS11, chloroplastic-like isoform X3 | *ShGPAT/ATS11* | Glycerolipid/Galactolipid Synthesis | LOC125221688 |
| glycerol-3-phosphate acyltransferase ATS11, chloroplastic-like | *ShGPAT/ATS11* | Glycerolipid/Galactolipid Synthesis | LOC125214421 |
| probable 1-acylglycerol-3-phosphate O-acyltransferase | *ShLPAAT* | Glycerolipid/Galactolipid Synthesis | LOC125207826 |
| 1-acyl-sn-glycerol-3-phosphate acyltransferase BAT2, chloroplastic-like isoform1 | *ShLPAT/BAT2* | Glycerolipid/Galactolipid Synthesis | LOC125192077 |
| 1-acyl-sn-glycerol-3-phosphate acyltransferase BAT2, chloroplastic-like isoform2 | *ShLPAT/BAT2* | Glycerolipid/Galactolipid Synthesis | LOC125192077 |
| 1-acyl-sn-glycerol-3-phosphate acyltransferase BAT2, chloroplastic-like isoform3 | *ShLPAT/BAT2* | Glycerolipid/Galactolipid Synthesis | LOC125192077 |
| 1-acyl-sn-glycerol-3-phosphate acyltransferase BAT2, chloroplastic-like isoform4 | *ShLPAT/BAT2* | Glycerolipid/Galactolipid Synthesis | LOC125192077 |
| 1-acyl-sn-glycerol-3-phosphate acyltransferase BAT2, chloroplastic-like isoform5 | *ShLPAT/BAT2* | Glycerolipid/Galactolipid Synthesis | LOC125192077 |
| 1-acyl-sn-glycerol-3-phosphate acyltransferase BAT2, chloroplastic-like isoform1 | *ShLPAT/BAT2* | Glycerolipid/Galactolipid Synthesis | LOC125190351 |
| 1-acyl-sn-glycerol-3-phosphate acyltransferase LPAT1, chloroplastic isoform2 | *ShLPAT1* | Glycerolipid/Galactolipid Synthesis | LOC125190351 |
| 1-acyl-sn-glycerol-3-phosphate acyltransferase 2-like | *ShLPAT2* | Glycerolipid/Galactolipid Synthesis | LOC125210432 |
| 1-acyl-sn-glycerol-3-phosphate acyltransferase 2-like | *ShLPAT2* | Glycerolipid/Galactolipid Synthesis | LOC125189421 |
| 1-acyl-sn-glycerol-3-phosphate acyltransferase 2 | *ShLPAT2* | Glycerolipid/Galactolipid Synthesis | LOC125187020 |
| probable 1-acyl-sn-glycerol-3-phosphate acyltransferase 4 | *ShLPAT4* | Glycerolipid/Galactolipid Synthesis | LOC125191737 |
| phosphatidate cytidylyltransferase 1 | *ShCDS1/CDP-DAGS* | Glycerolipid/Galactolipid Synthesis | LOC125193967 |
| phosphatidate cytidylyltransferase 1-like | *ShCDS1/CDP-DAGS* | Glycerolipid/Galactolipid Synthesis | LOC125200768 |
| phosphatidate cytidylyltransferase 1-like | *ShCDS1/CDP-DAGS* | Glycerolipid/Galactolipid Synthesis | LOC125208324 |
| phosphatidate cytidylyltransferase 4, chloroplastic-like | *ShCDS4/CDP-DAGS* | Glycerolipid/Galactolipid Synthesis | LOC125221392 |
| phosphatidate cytidylyltransferase 4, chloroplastic-like isoform X1 | *ShCDS4/CDP-DAGS* | Glycerolipid/Galactolipid Synthesis | LOC125196263 |
| phosphatidate cytidylyltransferase 4, chloroplastic-like isoform X2 | *ShCDS4/CDP-DAGS* | Glycerolipid/Galactolipid Synthesis | LOC125196263 |
| phosphatidate cytidylyltransferase 3-like | *ShCDS3/CDP-DAGS* | Glycerolipid/Galactolipid Synthesis | LOC125190529 |
| probable monogalactosyldiacylglycerol synthase, chloroplastic | *ShMGD1* | Glycerolipid/Galactolipid Synthesis | LOC125214688 |
| monogalactosyldiacylglycerol synthase 2, chloroplastic-like | *ShMGD2* | Glycerolipid/Galactolipid Synthesis | LOC125194428 |
| monogalactosyldiacylglycerol synthase 2, chloroplastic-like | *ShMGD2* | Glycerolipid/Galactolipid Synthesis | LOC125193157 |

| | | | |
|---|---|---|---|
| monogalactosyldiacylglycerol synthase 2, chloroplastic-like | *ShMGD2* | Glycerolipid/Galactolipid Synthesis | LOC125187479 |
| digalactosyldiacylglycerol synthase 1, chloroplastic | *ShDGD1* | Glycerolipid/Galactolipid Synthesis | LOC125221554 |
| digalactosyldiacylglycerol synthase 2, chloroplastic | *ShDGD2* | Glycerolipid/Galactolipid Synthesis | LOC125224292 |
| UTP--glucose-1-phosphate uridylyltransferase 3 | *ShUGP3* | Glycerolipid/Galactolipid Synthesis | LOC125189095 |
| UDP-sulfoquinovose synthase, chloroplastic | *ShSQD1* | Glycerolipid/Galactolipid Synthesis | LOC125213940 |
| sulfoquinovosyl transferase SQD2-like | *ShSQD2* | Glycerolipid/Galactolipid Synthesis | LOC125221525 |
| sulfoquinovosyl transferase SQD2-like | *ShSQD2* | Glycerolipid/Galactolipid Synthesis | LOC125220069 |
| fatty acid desaturase 4, chloroplastic-like | *Shfad4* | Glycerolipid/Galactolipid Synthesis | LOC125192766 |
| fatty acid desaturase 4-like 1, chloroplastic | *Shfad4* | Glycerolipid/Galactolipid Synthesis | LOC125209570 |
| palmitoyl-monogalactosyldiacylglycerol delta-7 desaturase | *Shfad5* | Glycerolipid/Galactolipid Synthesis | LOC125211395 |
| palmitoyl-monogalactosyldiacylglycerol delta-7 desaturase | *Shfad5* | Glycerolipid/Galactolipid Synthesis | LOC125208135 |
| palmitoyl-monogalactosyldiacylglycerol delta-7 desaturase | *Shfad5* | Glycerolipid/Galactolipid Synthesis | LOC125205438 |
| omega-6 fatty acid desaturase, chloroplastic | *Shfad6* | Glycerolipid/Galactolipid Synthesis | LOC125191028 |
| omega-3 fatty acid desaturase, chloroplastic-like | *Shfad7* | Glycerolipid/Galactolipid Synthesis | LOC125201775 |
| omega-3 fatty acid desaturase, chloroplastic-like | *Shfad8* | Glycerolipid/Galactolipid Synthesis | LOC125207882 |
| non-specific phospholipase C1 | *ShNPC1* | Glycerolipid/Galactolipid Synthesis | LOC125200788 |
| non-specific phospholipase C2 | *ShNPC2* | Glycerolipid/Galactolipid Synthesis | LOC125188758 |
| non-specific phospholipase C4-like | *ShNPC4* | Glycerolipid/Galactolipid Synthesis | LOC125202640 |
| non-specific phospholipase C6-like | *ShNPC6* | Glycerolipid/Galactolipid Synthesis | LOC125186867 |
| non-specific phospholipase C6-like | *ShNPC6* | Glycerolipid/Galactolipid Synthesis | LOC125202083 |
| phospholipase D zeta 1 | *ShPLDzeta 1* | Glycerolipid/Galactolipid Synthesis | LOC125205998 |
| phospholipase D zeta 1-like | *ShPLDzeta 1* | Glycerolipid/Galactolipid Synthesis | LOC125205600 |
| phosphatidate phosphatase PAH1-like | *ShPAH1* | Glycerolipid/Galactolipid Synthesis | LOC125201171 |
| phosphatidate phosphatase PAH1-like | *ShPAH1* | Glycerolipid/Galactolipid Synthesis | LOC125203258 |
| phosphatidate phosphatase PAH1-like isoform X1 | *ShPAH1* | Glycerolipid/Galactolipid Synthesis | LOC125214169 |
| phosphatidate phosphatase PAH1-like isoform X2 | *ShPAH1* | Glycerolipid/Galactolipid Synthesis | LOC125214169 |
| phosphatidate phosphatase PAH1-like isoform X1 | *ShPAH1* | Glycerolipid/Galactolipid Synthesis | LOC125222089 |
| phosphatidate phosphatase PAH1-like isoform X2 | *ShPAH1* | Glycerolipid/Galactolipid Synthesis | LOC125222089 |

| | | | |
|---|---|---|---|
| phosphatidate phosphatase PAH2-like | *ShPAH2* | Glycerolipid/Galactolipid Synthesis | LOC125212424 |
| lipid phosphate phosphatase epsilon 1, chloroplastic-like isoform X1 | *ShLPPepsilon 1* | Glycerolipid/Galactolipid Synthesis | LOC125217146 |
| lipid phosphate phosphatase epsilon 2, chloroplastic-like isoform X2 | *ShLPPepsilon 2* | Glycerolipid/Galactolipid Synthesis | LOC125217146 |
| lipid phosphate phosphatase epsilon 2, chloroplastic-like isoform X1 | *ShLPPepsilon 2* | Glycerolipid/Galactolipid Synthesis | LOC125214051 |
| lipid phosphate phosphatase epsilon 2, chloroplastic-like isoform X2 | *ShLPPepsilon 2* | Glycerolipid/Galactolipid Synthesis | LOC125214051 |
| probable lipid phosphate phosphatase beta | *ShLPPbeta* | Glycerolipid/Galactolipid Synthesis | LOC125219896 |
| lipid phosphate phosphatase gamma | *ShLPPgamma* | Glycerolipid/Galactolipid Synthesis | LOC125224217 |
| lipid phosphate phosphatase 2-like | *ShPAP2* | Glycerolipid/Galactolipid Synthesis | LOC125200353 |
| lipid phosphate phosphatase 2-like | *ShPAP2* | Glycerolipid/Galactolipid Synthesis | LOC125190836 |
| lipid phosphate phosphatase 2-like | *ShPAP2* | Glycerolipid/Galactolipid Synthesis | LOC125190836 |
| lipid phosphate phosphatase 2-like | *ShPAP2* | Glycerolipid/Galactolipid Synthesis | LOC125200323 |
| lipid phosphate phosphatase 2 | *ShPAP2* | Glycerolipid/Galactolipid Synthesis | LOC125204016 |
| lipid phosphate phosphatase 2 | *ShPAP2* | Glycerolipid/Galactolipid Synthesis | LOC125204016 |
| lipid phosphate phosphatase 2-like isoform X1 | *ShPAP2* | Glycerolipid/Galactolipid Synthesis | LOC125215764 |
| lipid phosphate phosphatase 2-like isoform X2 | *ShPAP2* | Glycerolipid/Galactolipid Synthesis | LOC125215764 |
| lipid phosphate phosphatase 2-like isoform X1 | *ShPAP2* | Glycerolipid/Galactolipid Synthesis | LOC125186282 |
| lipid phosphate phosphatase 2-like isoform X1 | *ShPAP2* | Glycerolipid/Galactolipid Synthesis | LOC125211545 |
| lipid phosphate phosphatase 2-like isoform X2 | *ShPAP2* | Glycerolipid/Galactolipid Synthesis | LOC125211545 |
| probable choline kinase 1 | *ShCK1* | Eukaryotic Phospholipid Synthesis | LOC125219769 |
| probable choline kinase 2 | *ShCK2* | Eukaryotic Phospholipid Synthesis | LOC125204129 |
| probable choline kinase 2 isoform X1 | *ShCK2* | Eukaryotic Phospholipid Synthesis | LOC125214444 |
| probable choline kinase 2 isoform X2 | *ShCK2* | Eukaryotic Phospholipid Synthesis | LOC125214444 |
| choline-phosphate cytidylyltransferase 1-like | *ShCCT1* | Eukaryotic Phospholipid Synthesis | LOC125214651 |
| choline-phosphate cytidylyltransferase 2-like | *ShCCT2* | Eukaryotic Phospholipid Synthesis | LOC125190883 |
| choline-phosphate cytidylyltransferase 2-like | *ShCCT2* | Eukaryotic Phospholipid Synthesis | LOC125193389 |
| choline/ethanolaminephosphotransferase 1 | *ShAPPT1* | Eukaryotic Phospholipid Synthesis | LOC125205505 |
| probable ethanolamine kinase | *ShETNK* | Eukaryotic Phospholipid Synthesis | LOC125201919 |
| probable ethanolamine kinase | *ShETNK* | Eukaryotic Phospholipid Synthesis | LOC125214905 |

| | | | |
|---|---|---|---|
| ethanolamine-phosphate cytidylyltransferase | *ShPECT* | Eukaryotic Phospholipid Synthesis | LOC125221953 |
| probable CDP-diacylglycerol--inositol 3-phosphatidyltransferase 2 | *ShPIS2* | Eukaryotic Phospholipid Synthesis | LOC125200569 |
| CDP-diacylglycerol--glycerol-3-phosphate 3-phosphatidyltransferase 2-like | *ShPGPS2* | Eukaryotic Phospholipid Synthesis | LOC125216065 |
| CDP-diacylglycerol--glycerol-3-phosphate 3-phosphatidyltransferase 2-like | *ShPGPS2* | Eukaryotic Phospholipid Synthesis | LOC125212846 |
| CDP-diacylglycerol--glycerol-3-phosphate 3-phosphatidyltransferase 2-isoform1 | *ShPGPS2* | Eukaryotic Phospholipid Synthesis | LOC125215010 |
| CDP-diacylglycerol--glycerol-3-phosphate 3-phosphatidyltransferase 2-isoform2 | *ShPGPS2* | Eukaryotic Phospholipid Synthesis | LOC125215010 |
| CDP-diacylglycerol--glycerol-3-phosphate 3-phosphatidyltransferase 2-isoform1 | *ShPGPS2* | Eukaryotic Phospholipid Synthesis | LOC125201212 |
| lysophospholipid acyltransferase 1-like | *ShLPCAT1* | Eukaryotic Phospholipid Synthesis | LOC125192484 |
| lysophospholipid acyltransferase 1-like | *ShLPEAT1* | Eukaryotic Phospholipid Synthesis | LOC125212344 |
| lysophospholipid acyltransferase LPEAT1-like isoform X1 | *ShLPEAT1* | Eukaryotic Phospholipid Synthesis | LOC125222801 |
| lysophospholipid acyltransferase LPEAT1-like isoform X2 | *ShLPEAT1* | Eukaryotic Phospholipid Synthesis | LOC125222801 |
| lysophospholipid acyltransferase LPEAT2-like | *ShLPEAT2* | Eukaryotic Phospholipid Synthesis | LOC125195695 |
| probable glycerol-3-phosphate dehydrogenase [NAD(+)] 1, cytosolic | *ShGPDHC1* | Eukaryotic Phospholipid Synthesis | LOC125196926 |
| probable glycerol-3-phosphate dehydrogenase [NAD(+)] 1, cytosolic | *ShGPDHC1* | Eukaryotic Phospholipid Synthesis | LOC125205961 |
| glycerol kinase | *ShGK* | Eukaryotic Phospholipid Synthesis | LOC125188838 |
| phosphoethanolamine N-methyltransferase 1 isoform X1 | *ShPEAMT* | Eukaryotic Phospholipid Synthesis | LOC125188971 |
| delta(12)-fatty-acid desaturase FAD2-like | *ShFAD2* | Eukaryotic Phospholipid Synthesis | LOC125205310 |
| delta(12)-fatty-acid desaturase FAD2-like | *ShFAD2* | Eukaryotic Phospholipid Synthesis | LOC125188161 |
| acyl-lipid omega-3 desaturase (cytochrome b5), endoplasmic reticulum-like | *ShFAD3* | Eukaryotic Phospholipid Synthesis | LOC125202091 |
| acyl-lipid omega-3 desaturase (cytochrome b5), endoplasmic reticulum-like | *ShFAD3* | Eukaryotic Phospholipid Synthesis | LOC125215068 |
| acyl-CoA-binding protein | *ShACBP* | Eukaryotic Phospholipid Synthesis | LOC125224105 |
| acyl-CoA-binding protein-like | *ShACBP* | Eukaryotic Phospholipid Synthesis | LOC125211122 |
| CDP-diacylglycerol--serine O-phosphatidyltransferase 1-like | *ShPSS1* | Eukaryotic Phospholipid Synthesis | LOC125204320 |
| CDP-diacylglycerol--serine O-phosphatidyltransferase 1-like isoform X1 | *ShPSS1* | Eukaryotic Phospholipid Synthesis | LOC125215583 |
| CDP-diacylglycerol--serine O-phosphatidyltransferase 1-like isoform X2 | *ShPSS1* | Eukaryotic Phospholipid Synthesis | LOC125215583 |
| CDP-diacylglycerol--serine O-phosphatidyltransferase 1-like isoform X3 | *ShPSS1* | Eukaryotic Phospholipid Synthesis | LOC125215583 |
| CDP-diacylglycerol--serine O-phosphatidyltransferase 1-like isoform X4 | *ShPSS1* | Eukaryotic Phospholipid Synthesis | LOC125215583 |
| phosphatidylserine decarboxylase proenzyme 2-like | *ShPSD2* | Eukaryotic Phospholipid Synthesis | LOC125220128 |

| | | | |
|---|---|---|---|
| phosphatidylcholine:diacylglycerol cholinephosphotransferase 1-like | *ShPDAT* | TG Synthesis | LOC125208152 |
| phospholipid:diacylglycerol acyltransferase 1-like isoform X1 | *ShPDAT* | TG Synthesis | LOC125186420 |
| phospholipid:diacylglycerol acyltransferase 1-like isoform X2 | *ShPDAT* | TG Synthesis | LOC125186420 |
| phospholipid:diacylglycerol acyltransferase 1-like | *ShPDAT* | TG Synthesis | LOC125205938 |
| phospholipid:diacylglycerol acyltransferase 1-like isoform X1 | *ShPDAT* | TG Synthesis | LOC125208495 |
| phospholipid:diacylglycerol acyltransferase 1-like isoform X2 | *ShPDAT* | TG Synthesis | LOC125208495 |
| phospholipid:diacylglycerol acyltransferase 1-like isoform X3 | *ShPDAT* | TG Synthesis | LOC125208495 |
| diacylglycerol O-acyltransferase 1A-like | *ShDGAT1* | TG Synthesis | LOC125187649 |
| diacylglycerol O-acyltransferase 1A-like | *ShDGAT1* | TG Synthesis | LOC125194449 |
| diacylglycerol O-acyltransferase 2D isoform X1 | *ShDGAT2* | TG Synthesis | LOC125211455 |
| diacylglycerol O-acyltransferase 2D isoform X2 | *ShDGAT2* | TG Synthesis | LOC125211455 |
| diacylglycerol O-acyltransferase 3 | *ShDGAT3* | TG Synthesis | LOC125196152 |
| diacylglycerol O-acyltransferase 3-like | *ShDGAT3* | TG Synthesis | LOC125216115 |
| triacylglycerol lipase 1 isoform X1 | *ShLIPASE1* | TG Degradation | LOC125223798 |
| triacylglycerol lipase 1 isoform X2 | *ShLIPASE1* | TG Degradation | LOC125223798 |
| triacylglycerol lipase 2-like | *ShLIPASE2* | TG Degradation | LOC125191219 |
| triacylglycerol lipase 2-like | *ShLIPASE2* | TG Degradation | LOC125188085 |
| triacylglycerol lipase SDP1-like | *ShSDP1* | TG Degradation | LOC125189115 |
| triacylglycerol lipase SDP1-like | *ShSDP1* | TG Degradation | LOC125190680 |
| triacylglycerol lipase OBL1-like | *ShOBL1* | TG Degradation | LOC125219294 |
| triacylglycerol lipase OBL1-like | *ShOBL1* | TG Degradation | LOC125217055 |
| triacylglycerol lipase OBL1-like | *ShOBL1* | TG Degradation | LOC125214023 |
| triacylglycerol lipase OBL1-like | *ShOBL1* | TG Degradation | LOC125208989 |
| triacylglycerol lipase OBL1-like | *ShOBL1* | TG Degradation | LOC125186766 |
| glyoxysomal fatty acid beta-oxidation multifunctional protein MFP isoform X1 | *ShMFP* | TG beta-Oxidation | LOC125208198 |
| glyoxysomal fatty acid beta-oxidation multifunctional protein MFP isoform X2 | *ShMFP* | TG beta-Oxidation | LOC125208198 |
| glyoxysomal fatty acid beta-oxidation multifunctional protein MFP-a | *ShMFP* | TG beta-Oxidation | LOC125203706 |
| probable enoyl-CoA hydratase 1, peroxisomal | *ShECH1* | TG beta-Oxidation | LOC125194356 |

| | | | |
|---|---|---|---|
| enoyl-CoA hydratase 2, peroxisomal-like | *ShECH2* | TG beta-Oxidation | LOC125204119 |
| enoyl-CoA hydratase 2, peroxisomal-like | *ShECH2* | TG beta-Oxidation | LOC125186330 |
| 3-ketoacyl CoA thiolase 1, peroxisomal-like | *ShKAT1* | TG beta-Oxidation | LOC125211318 |
| 3-ketoacyl CoA thiolase 1, peroxisomal-like | *ShKAT1* | TG beta-Oxidation | LOC125207798 |
| 3-ketoacyl CoA thiolase 2, peroxisomal-like | *ShKAT2* | TG beta-Oxidation | LOC125192223 |
| 3-ketoacyl CoA thiolase 2, peroxisomal-like | *ShKAT2* | TG beta-Oxidation | LOC125195696 |
| enoyl-CoA delta isomerase 1, peroxisomal-like | *ShECI1* | TG beta-Oxidation | LOC125208388 |
| enoyl-CoA delta isomerase 1, peroxisomal-like | *ShECI1* | TG beta-Oxidation | LOC125201303 |
| enoyl-CoA delta isomerase 1, peroxisomal-like | *ShECI1* | TG beta-Oxidation | LOC125201014 |
| enoyl-CoA delta isomerase 2, peroxisomal-like isoform X1 | *ShECI2* | TG beta-Oxidation | LOC125200563 |
| enoyl-CoA delta isomerase 2, peroxisomal-like isoform X2 | *ShECI2* | TG beta-Oxidation | LOC125200563 |
| delta(3,5)-Delta(2,4)-dienoyl-CoA isomerase, peroxisomal | *ShDCI1* | TG beta-Oxidation | LOC125203834 |
| peroxisomal 2,4-dienoyl-CoA reductase [(3E)-enoyl-CoA-producing]-like | *ShDECR* | TG beta-Oxidation | LOC125216932 |
| peroxisomal 2,4-dienoyl-CoA reductase [(3E)-enoyl-CoA-producing]-like | *ShDECR* | TG beta-Oxidation | LOC125193671 |
| peroxisomal 2,4-dienoyl-CoA reductase [(3E)-enoyl-CoA-producing]-like | *ShDECR* | TG beta-Oxidation | LOC125187272 |
| ethylene-responsive transcription factor WRI1-like | *ShWRI1* | TFs involved in TG Synthesis | LOC125202281 |
| ethylene-responsive transcription factor WRI1-like | *ShWRI1* | TFs involved in TG Synthesis | LOC125197031 |
| ethylene-responsive transcription factor WRI1-like | *ShWRI1* | TFs involved in TG Synthesis | LOC125197030 |
| ethylene-responsive transcription factor WRI1-like | *ShWRI1* | TFs involved in TG Synthesis | LOC125198608 |
| B3 domain-containing transcription factor ABI3 | *ShABI3* | TFs involved in TG Synthesis | LOC125218188 |
| B3 domain-containing transcription factor ABI3-like isoform X1 | *ShABI3* | TFs involved in TG Synthesis | LOC125211724 |

**Table S4.** Differentially expressed lipid metabolism genes in chia under heat stress.

| Locus ID | Abbreviation | Pathway | Treatment | $Log_2FC$ | $Log_2CPM$ | *p*-value |
|---|---|---|---|---|---|---|
| LOC125188702 | *ShKAS II* | Elongation, Desaturation/Export | Heat Shock (T1.H vs T0) | -0.393 | 5.950 | 4.79E-05 |
| LOC125215511 | *ShKAS II* | Elongation, Desaturation/Export | Heat Shock (T1.H vs T0) | -0.936 | 4.857 | 1.35E-10 |
| LOC125216292 | *ShKAS II* | Elongation, Desaturation/Export | Heat Shock (T1.H vs T0) | -0.750 | 4.489 | 5.38E-09 |
| LOC125214680 | *ShSAD/FAB2* | Elongation, Desaturation/Export | Heat Shock (T1.H vs T0) | -0.411 | 4.966 | 4.95E-08 |
| LOC125200966 | *ShSAD/FAB2* | Elongation, Desaturation/Export | Prolonged Heat (T2.H vs T0) | -0.296 | 4.966 | 2.48E-05 |
| LOC125214579 | *ShLACS8* | Elongation, Desaturation/Export | Prolonged Heat (T2.H vs T0) | 0.478 | 5.641 | 6.30E-06 |
| LOC125214680 | *ShSAD/FAB2* | Elongation, Desaturation/Export | Prolonged Heat (T2.H vs T0) | -0.521 | 7.512 | 2.93E-06 |
| LOC125223686 | *ShSAD/FAB2* | Elongation, Desaturation/Export | Prolonged Heat (T2.H vs T0) | -0.491 | 5.954 | 6.37E-06 |
| LOC125186718 | *ShGPD2* | Glycerolipid/Galactolipid Synthesis | Heat Shock (T1.H vs T0) | 0.393 | 4.393 | 1.72E-06 |
| LOC125186867 | *ShNPC6* | Glycerolipid/Galactolipid Synthesis | Heat Shock (T1.H vs T0) | 0.811 | 2.617 | 2.11E-05 |
| LOC125187479 | *ShMGD2* | Glycerolipid/Galactolipid Synthesis | Heat Shock (T1.H vs T0) | -0.674 | 2.386 | 9.79E-05 |
| LOC125193157 | *ShMGD2* | Glycerolipid/Galactolipid Synthesis | Heat Shock (T1.H vs T0) | -1.323 | 1.014 | 2.02E-07 |
| LOC125196263 | *ShCDP-DAGS* | Glycerolipid/Galactolipid Synthesis | Heat Shock (T1.H vs T0) | -0.626 | 4.920 | 3.45E-15 |
| LOC125201775 | *Shfad7* | Glycerolipid/Galactolipid Synthesis | Heat Shock (T1.H vs T0) | -0.772 | 7.734 | 8.35E-05 |
| LOC125205438 | *Shfad5* | Glycerolipid/Galactolipid Synthesis | Heat Shock (T1.H vs T0) | -0.733 | 8.107 | 1.10E-05 |
| LOC125208135 | *Shfad5* | Glycerolipid/Galactolipid Synthesis | Heat Shock (T1.H vs T0) | -1.107 | 0.421 | 1.57E-04 |
| LOC125210432 | *ShLPAT2* | Glycerolipid/Galactolipid Synthesis | Heat Shock (T1.H vs T0) | 0.360 | 5.496 | 2.89E-06 |
| LOC125211545 | *ShPAP2* | Glycerolipid/Galactolipid Synthesis | Heat Shock (T1.H vs T0) | -0.749 | 3.930 | 3.77E-08 |
| LOC125221392 | *ShCDP-DAGS* | Glycerolipid/Galactolipid Synthesis | Heat Shock (T1.H vs T0) | -0.575 | 3.637 | 1.29E-08 |
| LOC125186718 | *ShGPD2* | Glycerolipid/Galactolipid Synthesis | Prolonged Heat (T2.H vs T0) | 0.349 | 23.237 | 1.07E-05 |
| LOC125187479 | *ShMGD2* | Glycerolipid/Galactolipid Synthesis | Prolonged Heat (T2.H vs T0) | -0.737 | 21.310 | 2.21E-05 |
| LOC125191028 | *Shfad6* | Glycerolipid/Galactolipid Synthesis | Prolonged Heat (T2.H vs T0) | -1.157 | 44.917 | 9.65E-09 |
| LOC125192766 | *Shfad4* | Glycerolipid/Galactolipid Synthesis | Prolonged Heat (T2.H vs T0) | -1.130 | 17.923 | 8.49E-05 |
| LOC125193157 | *ShMGD2* | Glycerolipid/Galactolipid Synthesis | Prolonged Heat (T2.H vs T0) | -1.202 | 29.201 | 1.28E-06 |
| LOC125196206 | *ShGPD* | Glycerolipid/Galactolipid Synthesis | Prolonged Heat (T2.H vs T0) | -1.250 | 24.422 | 6.94E-06 |

| | | | | | | |
|---|---|---|---|---|---|---|
| LOC125196263 | *ShCDP-DAGS* | Glycerolipid/Galactolipid Synthesis | Prolonged Heat (T2.H vs T0) | -0.352 | 36.689 | 1.10E-07 |
| LOC125200768 | *ShCDP-DAGS* | Glycerolipid/Galactolipid Synthesis | Prolonged Heat (T2.H vs T0) | 0.775 | 31.327 | 6.24E-07 |
| LOC125204016 | *ShPAP2* | Glycerolipid/Galactolipid Synthesis | Prolonged Heat (T2.H vs T0) | -0.683 | 19.364 | 4.70E-05 |
| LOC125205998 | *ShPLDzeta 1* | Glycerolipid/Galactolipid Synthesis | Prolonged Heat (T2.H vs T0) | 0.491 | 22.613 | 1.35E-05 |
| LOC125207882 | *Shfad8* | Glycerolipid/Galactolipid Synthesis | Prolonged Heat (T2.H vs T0) | -1.799 | 78.193 | 2.79E-12 |
| LOC125211545 | *ShPAP2* | Glycerolipid/Galactolipid Synthesis | Prolonged Heat (T2.H vs T0) | -0.809 | 48.119 | 3.75E-09 |
| LOC125214688 | *ShMGD1* | Glycerolipid/Galactolipid Synthesis | Prolonged Heat (T2.H vs T0) | -0.402 | 19.700 | 4.12E-05 |
| LOC125221392 | *ShCDP-DAGS* | Glycerolipid/Galactolipid Synthesis | Prolonged Heat (T2.H vs T0) | -0.649 | 57.399 | 3.16E-10 |
| LOC125221525 | *ShSQD2* | Glycerolipid/Galactolipid Synthesis | Prolonged Heat (T2.H vs T0) | -0.369 | 19.198 | 5.01E-05 |
| LOC125222089 | *ShPAH1* | Glycerolipid/Galactolipid Synthesis | Prolonged Heat (T2.H vs T0) | 0.543 | 21.675 | 1.93E-05 |
| LOC125224217 | *ShLPPgamma* | Glycerolipid/Galactolipid Synthesis | Prolonged Heat (T2.H vs T0) | 0.408 | 16.821 | 1.30E-04 |
| LOC125188161 | *ShFAD2* | Eukaryotic Phospholipid Synthesis | Heat Shock (T1.H vs T0) | -0.437 | 6.278 | 2.75E-05 |
| LOC125188838 | *ShGK* | Eukaryotic Phospholipid Synthesis | Heat Shock (T1.H vs T0) | 1.123 | 5.137 | 8.47E-08 |
| LOC125188971 | *ShPEAMT* | Eukaryotic Phospholipid Synthesis | Heat Shock (T1.H vs T0) | -2.277 | 8.863 | 2.93E-08 |
| LOC125190883 | *ShCCT2* | Eukaryotic Phospholipid Synthesis | Heat Shock (T1.H vs T0) | -0.466 | 4.446 | 8.13E-07 |
| LOC125201919 | *ShETNK* | Eukaryotic Phospholipid Synthesis | Heat Shock (T1.H vs T0) | -0.597 | 4.171 | 4.10E-08 |
| LOC125204129 | *ShCK2* | Eukaryotic Phospholipid Synthesis | Heat Shock (T1.H vs T0) | 1.110 | 1.732 | 8.16E-05 |
| LOC125205310 | *ShFAD2* | Eukaryotic Phospholipid Synthesis | Heat Shock (T1.H vs T0) | -0.972 | 7.358 | 4.01E-08 |
| LOC125205505 | *ShAPPT1* | Eukaryotic Phospholipid Synthesis | Heat Shock (T1.H vs T0) | -0.745 | 4.737 | 4.74E-13 |
| LOC125215010 | *ShPGPS2* | Eukaryotic Phospholipid Synthesis | Heat Shock (T1.H vs T0) | -0.461 | 3.584 | 1.17E-05 |
| LOC125215068 | *ShFAD3* | Eukaryotic Phospholipid Synthesis | Heat Shock (T1.H vs T0) | -0.892 | 6.035 | 8.26E-06 |
| LOC125216065 | *ShPGPS2* | Eukaryotic Phospholipid Synthesis | Heat Shock (T1.H vs T0) | -0.564 | 4.487 | 1.03E-10 |
| LOC125219769 | *ShCK1* | Eukaryotic Phospholipid Synthesis | Heat Shock (T1.H vs T0) | 1.505 | 5.471 | 6.36E-12 |
| LOC125220128 | *ShPSD2* | Eukaryotic Phospholipid Synthesis | Heat Shock (T1.H vs T0) | 0.712 | 5.185 | 1.18E-09 |
| LOC125188161 | *ShFAD2* | Eukaryotic Phospholipid Synthesis | Prolonged Heat (T2.H vs T0) | -0.513 | 6.278 | 1.14E-06 |
| LOC125192484 | *ShLPCAT1* | Eukaryotic Phospholipid Synthesis | Prolonged Heat (T2.H vs T0) | -0.377 | 5.410 | 6.74E-07 |
| LOC125201919 | *ShETNK* | Eukaryotic Phospholipid Synthesis | Prolonged Heat (T2.H vs T0) | -0.470 | 4.171 | 4.58E-06 |
| LOC125202091 | *ShFAD3* | Eukaryotic Phospholipid Synthesis | Prolonged Heat (T2.H vs T0) | -1.019 | 6.085 | 1.46E-06 |

| | | | | | | |
|---|---|---|---|---|---|---|
| LOC125205505 | *ShAPPT1* | Eukaryotic Phospholipid Synthesis | Prolonged Heat (T2.H vs T0) | -0.372 | 4.737 | 1.60E-05 |
| LOC125215068 | *ShFAD3* | Eukaryotic Phospholipid Synthesis | Prolonged Heat (T2.H vs T0) | -1.717 | 6.035 | 2.68E-13 |
| LOC125222801 | *ShLPEAT1* | Eukaryotic Phospholipid Synthesis | Prolonged Heat (T2.H vs T0) | -0.335 | 4.752 | 5.72E-05 |
| LOC125186101 | *ShACP1* | Plastidial Fatty Acid Synthesis | Heat Shock (T1.H vs T0) | 1.418 | 8.088 | 6.33E-09 |
| LOC125187197 | *ShKAR/FabG* | Plastidial Fatty Acid Synthesis | Heat Shock (T1.H vs T0) | -0.955 | 1.161 | 2.26E-04 |
| LOC125188973 | *ShPDH E1 alpha* | Plastidial Fatty Acid Synthesis | Heat Shock (T1.H vs T0) | -0.727 | 6.695 | 2.07E-13 |
| LOC125192402 | *ShKAR/FabG* | Plastidial Fatty Acid Synthesis | Heat Shock (T1.H vs T0) | 0.470 | 2.732 | 1.46E-04 |
| LOC125193072 | *ShACP1* | Plastidial Fatty Acid Synthesis | Heat Shock (T1.H vs T0) | -0.676 | 4.849 | 3.18E-08 |
| LOC125194305 | *ShACP1* | Plastidial Fatty Acid Synthesis | Heat Shock (T1.H vs T0) | 0.730 | 5.869 | 1.88E-04 |
| LOC125202643 | *ShPDH E1 beta* | Plastidial Fatty Acid Synthesis | Heat Shock (T1.H vs T0) | -0.688 | 3.416 | 2.64E-06 |
| LOC125210931 | *ShENR* | Plastidial Fatty Acid Synthesis | Heat Shock (T1.H vs T0) | -0.611 | 4.289 | 1.07E-04 |
| LOC125211095 | *ShMCMT* | Plastidial Fatty Acid Synthesis | Heat Shock (T1.H vs T0) | -0.678 | 5.042 | 1.63E-06 |
| LOC125218497 | *ShACP1* | Plastidial Fatty Acid Synthesis | Heat Shock (T1.H vs T0) | -0.769 | 5.354 | 1.38E-07 |
| LOC125222552 | *ShACC αCT/CAC3* | Plastidial Fatty Acid Synthesis | Heat Shock (T1.H vs T0) | 0.398 | 7.925 | 4.96E-08 |
| LOC125187061 | *ShENR* | Plastidial Fatty Acid Synthesis | Prolonged Heat (T2.H vs T0) | -0.752 | 4.299 | 8.16E-05 |
| LOC125188973 | *ShPDH E1 alpha* | Plastidial Fatty Acid Synthesis | Prolonged Heat (T2.H vs T0) | -0.564 | 6.695 | 4.41E-10 |
| LOC125192402 | *ShKAR/FabG* | Plastidial Fatty Acid Synthesis | Prolonged Heat (T2.H vs T0) | 0.548 | 2.732 | 1.08E-05 |
| LOC125193072 | *ShACP1* | Plastidial Fatty Acid Synthesis | Prolonged Heat (T2.H vs T0) | -0.768 | 4.849 | 7.71E-10 |
| LOC125197323 | *ShPDH E1 alpha* | Plastidial Fatty Acid Synthesis | Prolonged Heat (T2.H vs T0) | -0.764 | 4.287 | 3.89E-09 |
| LOC125202643 | *ShPDH E1 beta* | Plastidial Fatty Acid Synthesis | Prolonged Heat (T2.H vs T0) | -0.852 | 3.416 | 1.95E-08 |
| LOC125208933 | *ShPDH E1 beta* | Plastidial Fatty Acid Synthesis | Prolonged Heat (T2.H vs T0) | -1.103 | 2.992 | 1.37E-09 |
| LOC125210332 | *ShACC BC/CAC2* | Plastidial Fatty Acid Synthesis | Prolonged Heat (T2.H vs T0) | -0.769 | 4.342 | 2.29E-06 |
| LOC125210931 | *ShENR* | Plastidial Fatty Acid Synthesis | Prolonged Heat (T2.H vs T0) | -0.737 | 4.289 | 4.17E-06 |
| LOC125211095 | *ShMCMT* | Plastidial Fatty Acid Synthesis | Prolonged Heat (T2.H vs T0) | -0.712 | 5.042 | 4.63E-07 |
| LOC125211668 | *ShHAD/FabZ* | Plastidial Fatty Acid Synthesis | Prolonged Heat (T2.H vs T0) | -0.537 | 4.469 | 4.73E-05 |
| LOC125213153 | *ShACC BC1* | Plastidial Fatty Acid Synthesis | Prolonged Heat (T2.H vs T0) | -0.890 | 5.797 | 2.82E-08 |
| LOC125218497 | *ShACP1* | Plastidial Fatty Acid Synthesis | Prolonged Heat (T2.H vs T0) | -0.603 | 5.354 | 1.27E-05 |
| LOC125186420 | *ShPDAT* | TG Synthesis | Heat Shock (T1.H vs T0) | 0.860 | 6.835 | 1.18E-06 |

| | | | | | | |
|---|---|---|---|---|---|---|
| LOC125205938 | *ShPDAT* | TG Synthesis | Heat Shock (T1.H vs T0) | 0.809 | 4.491 | 2.73E-11 |
| LOC125208152 | *ShPDAT* | TG Synthesis | Heat Shock (T1.H vs T0) | -0.982 | 3.674 | 6.23E-05 |
| LOC125208152 | *ShPDAT* | TG Synthesis | Prolonged Heat (T2.H vs T0) | -1.156 | 3.674 | 3.92E-06 |
| LOC125205938 | *ShPDAT* | TG Synthesis | Prolonged Heat (T2.H vs T0) | 0.408 | 4.491 | 8.47E-05 |
| LOC125194449 | *ShDGAT1* | TG Synthesis | Prolonged Heat (T2.H vs T0) | 0.402 | 4.092 | 1.97E-06 |
| LOC125186420 | *ShPDAT* | TG Synthesis | Prolonged Heat (T2.H vs T0) | 0.902 | 6.835 | 2.89E-07 |
| LOC125186766 | *ShOBL1* | TG Degradation | Prolonged Heat (T2.H vs T0) | 2.285 | 1.138 | 4.29E-07 |
| LOC125223798 | *ShLIPASE1* | TG Degradation | Heat Shock (T1.H vs T0) | 1.225 | 6.384 | 4.35E-19 |
| LOC125189115 | *ShSDP1* | TG Degradation | Heat Shock (T1.H vs T0) | 0.721 | 3.884 | 1.29E-04 |
| LOC125193671 | *ShDECR* | TG beta-Oxidation | Prolonged Heat (T2.H vs T0) | -0.317 | 5.891 | 1.01E-04 |
| LOC125204119 | *ShECH2* | TG beta-Oxidation | Prolonged Heat (T2.H vs T0) | -0.659 | 4.710 | 2.46E-09 |
| LOC125194356 | *ShECH1* | TG beta-Oxidation | Prolonged Heat (T2.H vs T0) | 0.777 | 3.213 | 1.49E-06 |
| LOC125195696 | *ShKAT2* | TG beta-Oxidation | Heat Shock (T1.H vs T0) | 0.783 | 6.343 | 1.52E-13 |
| LOC125203834 | *ShDCI1* | TG beta-Oxidation | Heat Shock (T1.H vs T0) | 0.816 | 5.984 | 2.67E-08 |
| LOC125208198 | *ShMFP* | TG beta-Oxidation | Heat Shock (T1.H vs T0) | -0.480 | 4.418 | 1.79E-04 |

**Table S5.** Gene Ontology (GO) enrichment analysis of differentially expressed genes (DEGs) under heat shock in chia (T1.H vs. T0): Top 20 enriched biological processes.

| GO Biological process | GO ID | FDR | Fold enrichment |
|---|---|---|---|
| Calcium import into the mitochondrion | GO:0036444 | 2.13E-02 | 4.81 |
| Mitochondrial calcium ion homeostasis | GO:0051560 | 2.13E-02 | 4.81 |
| Cytosolic calcium ion transport | GO:0060401 | 2.13E-02 | 4.81 |
| Mitochondrial calcium ion transmembrane transport | GO:0006851 | 2.13E-02 | 4.81 |
| RRNA pseudouridine synthesis | GO:0031118 | 1.68E-02 | 4.08 |
| Formaldehyde metabolic proc. | GO:0046292 | 3.47E-02 | 3.93 |
| Formaldehyde catabolic proc. | GO:0046294 | 3.47E-02 | 3.93 |
| Cellular detoxification of aldehyde | GO:0110095 | 3.47E-02 | 3.93 |
| Cellulose catabolic proc. | GO:0030245 | 2.09E-04 | 3.41 |
| Beta-glucan catabolic proc. | GO:0051275 | 2.09E-04 | 3.41 |
| Regulation of transcription elongation of RNA polymerase II promoter | GO:0034243 | 1.86E-03 | 3.10 |
| Post regulation transcription elongation of RNA polymerase II promoter | GO:0032968 | 7.19E-03 | 3.02 |
| Phosphorylation of RNA polymerase II C-terminal domain | GO:0070816 | 7.19E-03 | 3.02 |
| Post regulation of DNA-templated transcription, elongation | GO:0032786 | 1.88E-02 | 2.57 |
| DNA-templated transcription, termination | GO:0006353 | 3.47E-02 | 2.16 |
| Glucan catabolic proc. | GO:0009251 | 3.47E-02 | 2.10 |
| Lipid catabolic proc. | GO:0016042 | 6.86E-07 | 1.86 |
| Carbohydrate transport | GO:0008643 | 7.35E-03 | 1.76 |
| Protein phosphorylation | GO:0006468 | 1.54E-15 | 1.55 |
| Protein dephosphorylation | GO:0006470 | 3.16E-02 | 1.51 |

**Table S6.** Gene Ontology (GO) enrichment analysis of differentially expressed genes (DEGs) under heat shock in chia (T1.H vs. T0): Top 20 enriched molecular functions.

| GO Molecular function | GO ID | FDR | Fold enrichment |
|---|---|---|---|
| Pectate lyase activity | GO:0030570 | 2.98E-12 | 4.78 |
| Alcohol dehydrogenase (NAD+) activity | GO:0004022 | 1.23E-06 | 4.37 |
| Glucan endo-1,3-beta-D-glucosidase activity | GO:0042973 | 4.85E-14 | 3.89 |
| Polygalacturonase activity | GO:0004650 | 8.06E-20 | 3.84 |
| Carbon-oxygen lyase activity, acting on polysaccharides | GO:0016837 | 9.09E-09 | 3.80 |
| D-threo-aldose 1-dehydrogenase activity | GO:0047834 | 5.61E-05 | 3.57 |
| Cellulase activity | GO:0008810 | 2.51E-05 | 3.29 |
| ADP binding | GO:0043531 | 1.86E-21 | 2.75 |
| NAD+ nucleosidase activity | GO:0003953 | 8.84E-14 | 2.74 |
| NAD(P)+ nucleosidase activity | GO:0050135 | 8.84E-14 | 2.74 |
| NAD+ nucleotidase, cyclic ADP-ribose generating | GO:0061809 | 8.84E-14 | 2.74 |
| Aspartic-type endopeptidase activity | GO:0004190 | 3.87E-08 | 2.70 |
| Aspartic-type peptidase activity | GO:0070001 | 3.87E-08 | 2.70 |
| Protein serine phosphatase activity | GO:0106306 | 5.90E-08 | 2.27 |
| Protein serine/threonine phosphatase activity | GO:0004722 | 4.14E-08 | 2.22 |
| Hydrolase activity, hydrolysing N-glycosyl compounds | GO:0016799 | 3.32E-08 | 2.12 |
| Ubiquitin protein ligase activity | GO:0061630 | 7.05E-10 | 2.00 |
| Hydrolase activity, acting on glycosyl bonds | GO:0016798 | 3.28E-15 | 1.80 |
| Protein kinase activity | GO:0004672 | 2.29E-24 | 1.72 |
| Methyltransferase activity | GO:0008168 | 1.24E-08 | 1.72 |

**Table S7.** Gene Ontology (GO) enrichment analysis of differentially expressed genes (DEGs) under heat shock in chia (T1.H vs. T0): Top 20 enriched cellular components.

| GO Cellular component | GO ID | FDR | Fold enrichment |
|---|---|---|---|
| Cul4A-RING E3 ubiquitin ligase complex | GO:0031464 | 7.50E-04 | 4.99 |
| Calcium channel complex | GO:0034704 | 1.06E-02 | 4.81 |
| Uniplex complex | GO:1990246 | 1.06E-02 | 4.81 |
| Cation channel complex | GO:0034703 | 2.08E-02 | 4.21 |
| Endoplasmic reticulum-Golgi intermediate compartment | GO:0005793 | 6.71E-04 | 3.42 |
| Nucleosome | GO:0000786 | 3.87E-08 | 3.03 |
| DNA packaging complex | GO:0044815 | 2.83E-07 | 2.81 |
| Histone deacetylase complex | GO:0000118 | 1.36E-03 | 2.73 |
| Cul4-RING E3 ubiquitin ligase complex | GO:0080008 | 2.08E-02 | 1.78 |
| Protein-DNA complex | GO:0032993 | 2.08E-02 | 1.73 |
| Anchored component of plasma membrane | GO:0046658 | 2.08E-02 | 1.59 |
| Cullin-RING ubiquitin ligase complex | GO:0031461 | 2.08E-02 | 1.53 |
| Chromatin | GO:0000785 | 2.91E-02 | 1.51 |
| Ubiquitin ligase complex | GO:0000151 | 2.08E-02 | 1.44 |

**Table S8.** Gene Ontology (GO) enrichment analysis of differentially expressed genes (DEGs) under prolonged heat stress in chia (T2.H vs. T0): Enriched biological processes.

| GO Biological process | GO ID | FDR | Fold enrichment |
|---|---|---|---|
| Formaldehyde metabolic proc. | GO:0046292 | 1.85E-04 | 8.87 |
| Formaldehyde catabolic proc. | GO:0046294 | 1.85E-04 | 8.87 |
| Cellular detoxification of aldehyde | GO:0110095 | 1.85E-04 | 8.87 |
| Cellular response to aldehyde | GO:0110096 | 3.24E-04 | 8.06 |
| Reg. of transcription elongation from RNA polymerase II promoter | GO:0034243 | 2.37E-08 | 6.99 |
| Post regulation transcription elongation from RNA polymerase II promoter | GO:0032968 | 2.98E-07 | 6.82 |
| Phosphorylation of RNA polymerase II C-terminal domain | GO:0070816 | 2.98E-07 | 6.82 |
| Pos. reg. of DNA-templated transcription, elongation | GO:0032786 | 3.84E-07 | 5.79 |
| DNA-templated transcription, termination | GO:0006353 | 2.84E-07 | 4.87 |
| DNA-templated transcription, elongation | GO:0006354 | 2.06E-04 | 3.75 |
| Peptidyl-tyrosine phosphorylation | GO:0018108 | 2.06E-04 | 3.59 |
| Peptidyl-tyrosine modification | GO:0018212 | 2.06E-04 | 3.59 |
| Protein phosphorylation | GO:0006468 | 1.77E-93 | 3.38 |
| Recognition of pollen | GO:0048544 | 2.38E-04 | 2.68 |
| Phosphorylation | GO:0016310 | 3.98E-67 | 2.66 |
| Protein autophosphorylation | GO:0046777 | 1.98E-06 | 2.64 |
| Cell recognition | GO:0008037 | 3.08E-04 | 2.63 |
| Pollen-pistil interaction | GO:0009875 | 8.03E-04 | 2.44 |

**Table S9.** Gene Ontology (GO) enrichment analysis of DEGs under prolonged heat stress (T2.H vs T0), highlighting the enriched molecular functions (FDR <0.05).

| GO Molecular function | GO ID | FDR | Fold enrichment |
|---|---|---|---|
| Clathrin heavy chain binding | GO:0032050 | 4.11E-11 | 5.10 |
| 1-phosphatidylinositol binding | GO:0005545 | 9.21E-10 | 5.05 |
| Pectate lyase activity | GO:0030570 | 2.52E-12 | 4.80 |
| Glucan endo-1,3-beta-D-glucosidase activity | GO:0042973 | 3.62E-15 | 4.01 |
| Polygalacturonase activity | GO:0004650 | 5.82E-21 | 3.93 |
| Carbon-oxygen lyase activity, acting on polysaccharides | GO:0016837 | 6.18E-09 | 3.81 |
| Aspartic-type endopeptidase activity | GO:0004190 | 1.84E-09 | 2.86 |
| Aspartic-type peptidase activity | GO:0070001 | 1.84E-09 | 2.86 |
| NAD+ nucleosidase activity | GO:0003953 | 6.60E-14 | 2.75 |
| NAD(P)+ nucleosidase activity | GO:0050135 | 6.60E-14 | 2.75 |
| NAD+ nucleotidase, cyclic ADP-ribose generating | GO:0061809 | 6.60E-14 | 2.75 |
| ADP binding | GO:0043531 | 5.48E-21 | 2.74 |
| Ubiquitin protein ligase activity | GO:0061630 | 1.15E-09 | 1.99 |
| Hydrolase activity, acting on glycosyl bonds | GO:0016798 | 3.62E-15 | 1.80 |
| Protein kinase activity | GO:0004672 | 4.12E-25 | 1.73 |
| Phosphotransferase activity, alcohol group as acceptor | GO:0016773 | 6.61E-19 | 1.58 |
| Protein serine/threonine kinase activity | GO:0004674 | 7.99E-11 | 1.53 |
| Kinase activity | GO:0016301 | 3.56E-15 | 1.48 |
| Transferase activity, transferring phosphorus-containing groups | GO:0016772 | 6.18E-09 | 1.32 |

**Table S10.** Gene Ontology (GO) enrichment analysis of DEGs under prolonged heat stress (T2.H vs T0), highlighting the enriched cellular components (FDR <0.05).

| GO cellular components | GO ID | FDR | Fold enrichment |
|---|---|---|---|
| Cul4A-RING E3 ubiquitin ligase complex | GO:0031464 | 4.59E-04 | 5.01 |
| Clathrin-coated pit | GO:0005905 | 3.51E-06 | 3.57 |
| Nucleosome | GO:0000786 | 1.61E-08 | 3.04 |
| DNA packaging complex | GO:0044815 | 1.58E-07 | 2.82 |
| Anchored component of plasma membrane | GO:0046658 | 1.61E-08 | 2.18 |
| Plasma membrane region | GO:0098590 | 2.39E-02 | 2.00 |
| Cul4-RING E3 ubiquitin ligase complex | GO:0080008 | 2.39E-02 | 1.78 |
| Protein-DNA complex | GO:0032993 | 2.39E-02 | 1.74 |
| Anchored component of membrane | GO:0031225 | 7.67E-05 | 1.60 |
